# Identification of topoisomerase as a precision-medicine target in chromatin reader SP140-driven Crohn’s disease

**DOI:** 10.1101/2021.09.20.461083

**Authors:** Hajera Amatullah, Sreehaas Digumarthi, Isabella Fraschilla, Fatemeh Adiliaghdam, Gracia Bonilla, Lai Ping Wong, Ruslan I. Sadreyev, Kate L. Jeffrey

## Abstract

How mis-regulated chromatin directly impacts human immunological disease is poorly understood. Speckled Protein 140 (SP140) is an immune-restricted PHD and bromodomain-containing chromatin ‘reader’ whose loss-of-function associates with Crohn’s disease (CD), multiple sclerosis (MS) and chronic lymphocytic leukemia (CLL). However, mechanisms underlying SP140-driven pathogenicity and therapeutic approaches that rescue SP140 remain unexplored. Using a global proteomic strategy, we identified SP140 as a repressor of topoisomerases (TOP) that maintains heterochromatin and immune cell fate. In humans and mice, SP140 loss resulted in unleashed TOP activity, genome instability, severely compromised lineage-defining and microbe-inducible innate transcriptional programs and defective bacterial killing. Pharmacological inhibition of TOP1 or TOP2 rescued these defects. Furthermore, exacerbated colitis was restored with TOP1 or TOP2 inhibitors in Sp140^−/−^ mice, but not wild-type mice, *in vivo.* Collectively, we identify SP140 as a repressor of topoisomerases and reveal repurposing of TOP inhibition as a precision strategy for reversing SP140-driven immune disease.

## Introduction

Chromatin regulatory proteins are indispensable for both maintenance of immune cell identity and integration of environmental cues. In particular, recognition of dynamic posttranslational modifications on histones by chromatin ‘readers’ is a critical step in this process (Smale et al., 2014; Soshnev et al., 2016; Wan et al., 2020; Wen et al., 2014). Dysregulated chromatin readers and aberrant chromatin architecture are central events in cancer (Filippakopoulos et al., 2010; Wan et al., 2020; Wen et al., 2014), therefore, targeting such pathways holds clinical promise (Dawson et al., 2011; Filippakopoulos et al., 2010; Nicodeme et al., 2010). Yet little is known about how altered chromatin readers may directly contribute to and facilitate tailored therapies for human immune diseases.

The complex immune disorders Crohn’s disease (CD), multiple sclerosis (MS) and chronic lymphocytic leukemia (CLL) result from both genetic susceptibility and environmental cues and despite their clinical heterogeneity, share many identified genetic risk variants, suggesting common pathogenic mechanisms (Cotsapas and Hafler, 2013; Franke et al., 2010; International Multiple Sclerosis Genetics et al., 2013; Jostins et al., 2012; Sille et al., 2012). Mutations within chromatin reader, SP140, is one such common genetic risk factor for these three immune diseases (Fraschilla and Jeffrey, 2020; International Multiple Sclerosis Genetics et al., 2013; Jostins et al., 2012; Sille et al., 2012). SP140 belongs to the Speckled Protein (SP) family consisting of SP100, SP110, and SP140L, which have high homology with Autoimmune Regulator (AIRE) (Fraschilla and Jeffrey, 2020; Gibson et al., 1998). SPs all contain an N-terminal CARD domain, an intrinsically disordered region (IDR) and multiple ‘reader’ modules including a SAND domain, a plant homeodomain (PHD) and a bromodomain (BRD) (Fraschilla and Jeffrey, 2020) (**Fig 1a**) indicative of a role in chromatin regulation. Moreover, SP family members are components of promyelocytic leukemia nuclear bodies (PML-NBs), which are ill-defined subnuclear structures regulated by a variety of cellular stresses such as virus infection, interferon (IFN) and DNA damage, with implicated roles in many cellular processes, including gene repression and higher order chromatin organization (Huoh et al., 2020; Lallemand-Breitenbach and de The, 2010)

**Figure 1.**
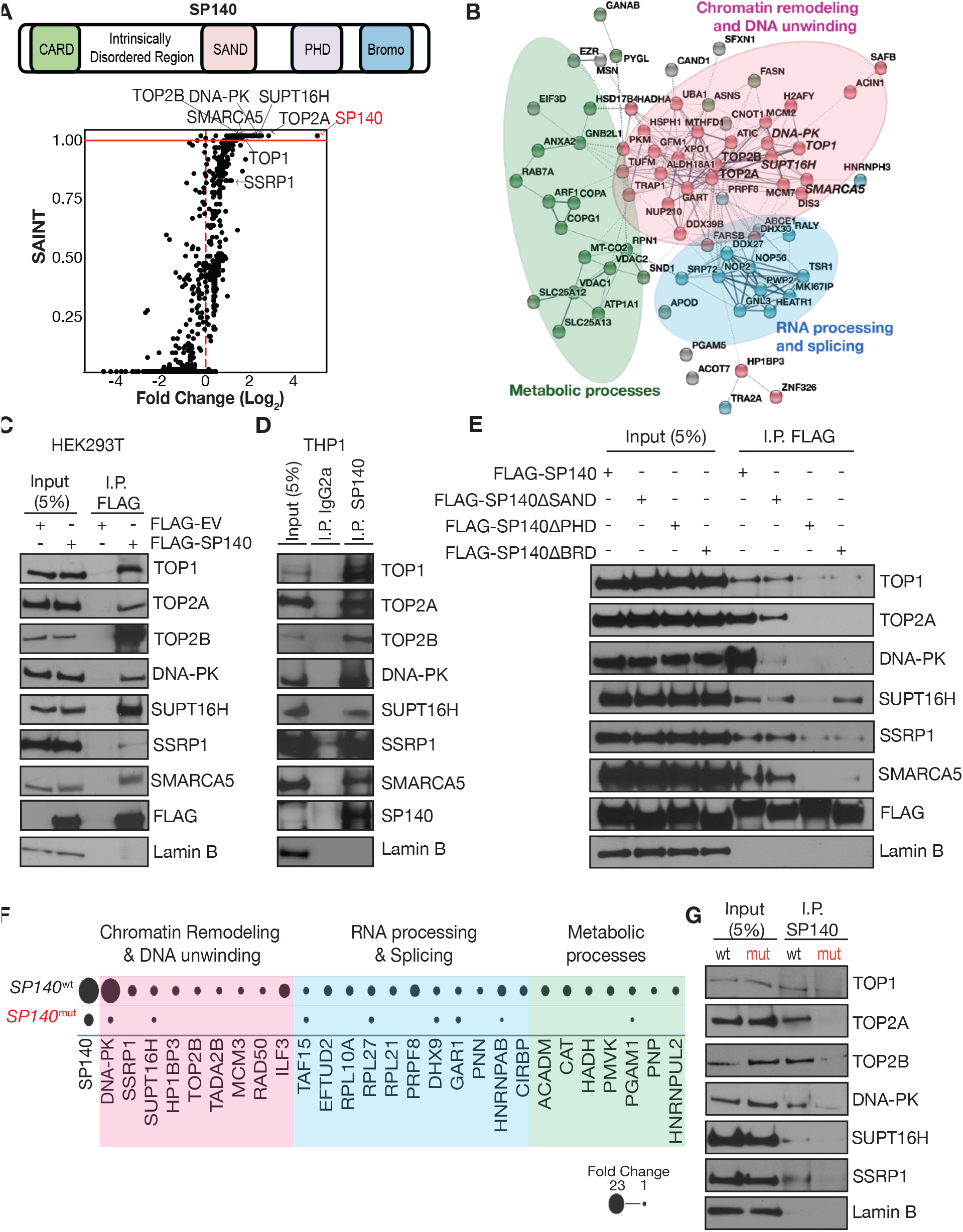
SP140 interactome includes DNA unwinding and chromatin remodeling proteins, including topoisomerases. **A,** Schematic of SP140 protein domains. SP140 interacting proteins identified using Mass Spectrometry (MS) of HEK293T nuclear lysates overexpressing FLAG-SP140 plotted as log_2_ fold change (over FLAG empty vector control) versus Significance Analysis of Interactome (SAINT) values. SAINT score ≥ 0.99 and FC>2 is indicated above the red line. Data is generated from n=2. **B,** Visual representation of SP140 interacting proteins (with SAINT score ≥ 0.99 and FC>2) using k-means clustering on STRING database (https://string-db.org/). **C,** Immunoprecipitation (IP) of FLAG-EV and FLAG-SP140 and immunoblot for endogenous Topoisomerase I (TOP1), Topoisomerase II alpha (TOP2A) and beta (TOP2B), DNA-dependent protein kinase (DNA-PK), Facilitates chromatin transcription (FACT) complex subunit SPT16 (SUPT16H), FACT complex subunit SSRP1 (SSRP1), and SWI/SNF-related matrix-associated actin-dependent regulator of chromatin subfamily A member 5 (SMARCA5) in HEK293T nuclear lysates. Lamin b is loading control. **D,** IP of endogenous SP140 and immunoblot for indicated endogenous proteins in human THP1 monocyte nuclear lysates. **E,** IP of FLAG-SP140 or FLAG-SP140 mutants lacking reader modules SAND (SP140ΔSAND), plant homeobox domain (PHD, SP140ΔPHD), or bromodomain (SP140ΔBRD) and immunoblot for indicated endogenous proteins in HEK293T nuclear lysates. **F,** Mass Spectrometry identification of SP140 interactors in patient-derived lymphoblastoid cell lines (LBL) with control SP140 (SP140^wt^) and SP140 SNP (SP140^mut^). Data are fold change of endogenous SP140 IP mass spectrometry (MS) peptide hits over IgG IP control. All proteins identified are in Extended Fig 4. **G,** IP of endogenous SP140 and immunoblot for indicated proteins in LBL nuclear lysates with indicated SP140 genotypes. **C-E** are representative of 3 experiments, **F** is mean of two biological replicates of each SP140 genotype, **G** is representative of 2 experiments.

Of the SPs, SP140 expression is uniquely restricted to immune cells (Bloch et al., 1996), with highest levels in activated macrophages and mature B cells (Mehta et al., 2017). Initial studies found SP140 to be essential for macrophage cytokine responses to bacteria or virus (Ji et al., 2021; Karaky et al., 2018; Mehta et al., 2017). Moreover, we recently demonstrated that SP140 acts as a co-repressor of lineage inappropriate genes such as *HOX* in mature human macrophages, preferentially occupying promoters of silenced genes within facultative heterochromatin regions marked by H3K27me3 in order to maintain macrophage identity and functional responses (Mehta et al., 2017). The CD, MS and CLL-associated variants of SP140 alter SP140 mRNA splicing and diminish SP140 protein (Matesanz et al., 2015; Mehta et al., 2017). Therefore, we asked mechanistically how this immune chromatin reader normally acts and how perturbations lead to altered SP140 structure and function on chromatin and eventually immune disease, with the intent to identify therapeutic interventions.

### Elucidation of the SP140 interactome: DNA unwinders and chromatin remodelers

To gain molecular insights into the function of SP140, we elucidated the SP140 interactome. We ectopically expressed FLAG-tagged human SP140 or FLAG-Empty Vector (FLAG-EV) in HEK293T cells (**Figure S1A, Related to Figure 1**) and subsequently performed Mass Spectrometry on FLAG co-immunoprecipitants. To efficiently discriminate confident interacting proteins from false-positive or contaminant proteins, the probability of bona fide protein-protein interactions was analyzed using the contaminant repository of affinity matrix (CRAPome) and Significance Analysis of INTeractome scores (SAINT) (Mellacheruvu et al., 2013) (**Figure 1A, Figure S1B**, **Supplementary Table 1**). This approach identified the top significant interactors of SP140 to be: Topoisomerases (TOP)1, TOP2A, TOP2B, DNA-dependent Protein Kinase (DNA-PK), Facilitates Chromatin Transactions (FACT) subunits SUPT16H and SSRP1, as well as the ATPase subunit of the ISWI family of chromatin remodelers, SWI/SNF Related, Matrix Associated, Actin Dependent Regulator of Chromatin, Subfamily A, member 5 (SMARCA5, **Figure 1A**). The top 1% of SP140 interacting partners (with False Discovery Rate, FDR, of less than 1%) broadly clustered under three functional groups (Szklarczyk et al., 2019): ‘chromatin organization and DNA unwinding’, ‘RNA processing and splicing’, and ‘metabolic processes’ (**Figure 1B**). Gene Ontology analyses confirmed the SP140 interactome was associated with ‘DNA topological change’, ‘nucleosome organization’ and ‘DNA unwinding’ (**Figure S1C, Related to Figure 1**).

### Topoisomerases (TOP) 1 and 2 are SP140 protein partners

We previously demonstrated that SP140 represses chromatin accessibility in human macrophages (Mehta et al., 2017), so we explored SP140 interactions with chromatin organization and DNA unwinding proteins TOP1, TOP2A, TOP2B, DNA-PK SUPT16H, SSRP1, and SMARCA5. Through immunoprecipitation of FLAG-SP140 in HEK293T expressing FLAG-SP140 or FLAG-EV and immunoblot of FLAG, we confirmed SP140 interacted with endogenous TOP1, TOP2A, TOP2B, DNA-PK, SUPT16H, SSRP1 and SMARCA5 (**Figure 1C**). Reciprocal immunoprecipitation of endogenous TOP1, TOP2A, DNA-PK, SUPT16H, SSRP1 and SMARCA5 and immunoblot of SP140 validated these interactions (**Figure S1D, Related to Figure 1**). Moreover, immunoprecipitation of endogenous SP140 in human monocyte THP1 cells confirmed these interactions exist in relevant immune cells (**Figure 1D**). SP140 complexes with TOP1, TOP2A, TOP2B, DNA-PK, SUPT16H, SSRP1 and SMARCA5 required structured DNA, since addition of the intercalating agent ethidium bromide (EtBr) disrupted their interactions (**Figure S2A,B, Related to Figure 1**). Furthermore, the chromatin “reading” PHD and bromodomain of SP140 were also necessary for these protein-protein interactions (**Figure 1e**). We next assessed whether topoisomerases or DNA-PK may be primary interacting proteins upon which the complex formed. Exposure of cells to TOP1 inhibitor Topotecan (TPT) or TOP2 inhibitor Etoposide (ETO, **Figure S2C, Related to Figure 1**) or siRNA-mediated knockdown of *TOP1* or *TOP2A* resulted in a loss of SP140 and TOP interaction but did not affect SP140 interactions with DNA-PK, SUPT16H or SMARCA5 (**Figure S2D, Related to Figure 1**). Similarly, DNA-PK (*PRKDC*) siRNA led to partial loss of SP140 binding to TOP1 but all other interactions remained intact (**Figure S2E, Related to Figure 1**).Thus, SP140 forms multiple distinct protein complexes, characteristic of flexible IDR-containing proteins that form phase-separated condensates (Sabari et al., 2018). Many of the SP140 interacting proteins we identified were previously found to interact with AIRE, which shares protein homology with SP140 (**Figure S3A, Related to Figure 1**) (Abramson et al., 2010; Bansal et al., 2017). However, whereas AIRE induces transcription of peripheral-tissue antigens in medullary thymic epithelial cells (mTECs) (Mathis and Benoist, 2009), our current working model is one in which SP140 functions as a transcriptional repressor. Consistently, the common SP140 and AIRE interacting partners included chromatin remodeling and DNA unwinding proteins (TOP1, TOP2A, TOP2B, DNA-PK and SUPT16H) while transcriptional elongation machinery (PTEFb and BRD4) were exclusively associated with AIRE and not SP140 (**Figure S3B, Related to Figure 1**). Furthermore, although SP140 and AIRE are generally expressed in different cell types (Fraschilla and Jeffrey, 2020), we co-transfected AIRE and SP140 and found that SP140 and AIRE did not compete for binding of endogenous TOP1 or TOP2A (**Figure S3C, Related to Figure 1**) demonstrating that SP140 and AIRE interact with distinct pools of topoisomerases.

### Loss of SP140-TOP interactions in patients bearing SP140 mutants

We next examined the consequence of disease risk loss-of-function SP140 on its ability to form the identified protein complexes. Similar to observations in HEK293T and THP1 cells, the dominant interactors of endogenous SP140 in patient-derived lymphoblastoid B cell lines (LBLs) identified by Mass Spectrometry, and confirmed by co-IP, were chromatin remodeling and DNA unwinding proteins TOP1, TOP2A, TOP2B, DNA-PK, SUPT16H and SSRP1 (**Figure 1F,G**). Importantly, these interactions were lost in LBLs derived from patients homozygous for loss-of-function immune disease-associated genetic variants of SP140 (SP140^mut^, **Figure 1F,G**). Notably, there was a re-wiring of the SP140 proteome in SP140^mut^ immune cells, with enhanced interactions between SP140 and proteins associated RNA processing and splicing factors (**Figure S4, Related to Figure 1**) possibly skewing SP140 toward these functions. Together, these data reveal that SP140 interacts with essential DNA unwinding and chromatin remodeling proteins, and these complexes are disrupted with CD, MS or CLL-risk SP140 loss-of-function.

### SP140 is a negative regulator of TOP 1 and 2 activity

The topoisomerase enzymes TOP1 and TOP2A/B cleave one or both DNA strands, respectively to resolve DNA topological constraints for replication, transcription, and chromatin organization (Pommier et al., 2016). For transcription, topoisomerases are indispensable for the relief of torsional strain as a result of chromatin remodeling, nucleosome depletion at enhancers or promoters, as well as RNA Polymerase (Pol) II pausing and elongation (Pommier et al., 2016). Furthermore, in addition to RNA Pol II itself (Baranello et al., 2016), multiple chromatin regulators and remodelers alter TOP by promoting or shielding TOP1 or 2 activity and binding (Abramson et al., 2010; Baranello et al., 2016; Dykhuizen et al., 2013; Husain et al., 2016). SP140 predominantly occupies facultative heterochromatin (Mehta et al., 2017) so we postulated that SP140 negatively regulates topoisomerases to prevent unsolicited transcription or chromatin decompaction at dynamic silent regions. Intriguingly, purified recombinant SP140 protein suppressed the ability of recombinant TOP1 to nick and unwind a supercoiled DNA substrate *in vitro* (**Figure 2A**). Furthermore, recombinant SP140 significantly decreased TOP2A decatenatory activity in a dose-dependent manner (**Figure 2B**). Upon SP140 depletion in THP1 monocytes or primary human peripheral blood derived macrophages, we observed significantly increased TOP1 activity in nuclear lysates, both at baseline and upon lipopolysaccharide (LPS) stimulation (**Figure 2C,E**) despite unchanged TOP1 protein levels (**Figure 2D,F**). Conversely, overexpression of SP140 in HEK293T cells suppressed nuclear TOP1 activity, in a dose-dependent manner to levels of repression observed by Topotecan (TPT), an FDA-approved TOP1 inhibitor that intercalates between cleaved DNA ends in the active site of the enzyme (Vos et al., 2011) (**Figure 2G,H**). Importantly, immune cells bearing loss-of-function SP140 displayed significantly increased TOP1 activity that could be reversed with TPT administration (**Figure 2I,J**). Thus, our results identified SP140 as a novel suppressor of TOP1 and TOP2 activity and unmask a specific compromise in this hallmark function of SP140 in patients bearing CD, MS and CLL-associated genetic variants of SP140.

**Figure 2.**
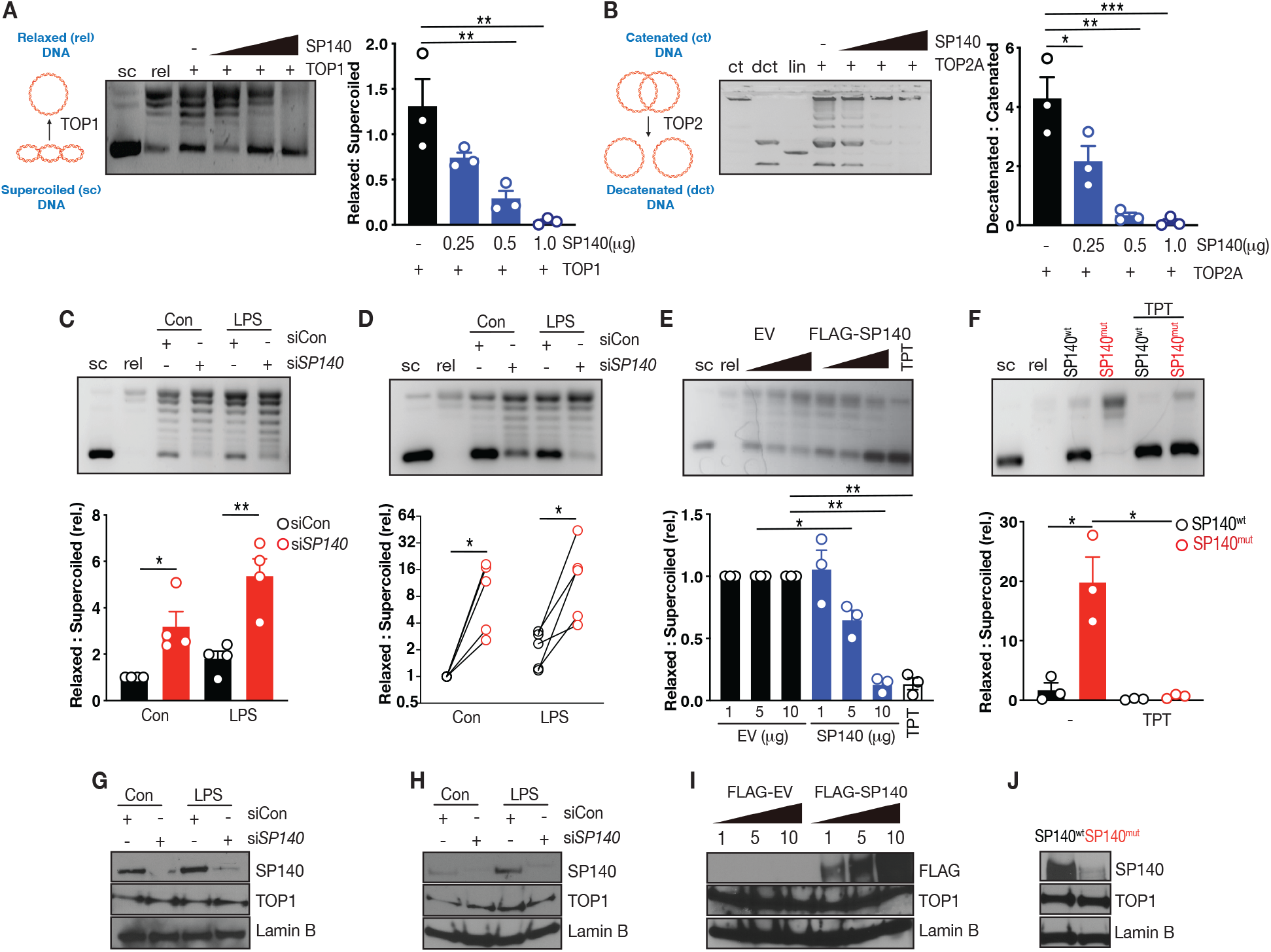
SP140 is a suppressor of topoisomerases TOP1 and TOP2. **A,** Left, schematic of topoisomerase 1 (TOP1) activity assay using migration and quantification of supercoiled (sc) and relaxed (rel) DNA following incubation of TOP1 with supercoiled pHOT DNA. Right, representative gel image and quantification of recombinant TOP1 activity (10 activity units, ~120ng) in the presence of indicated concentrations of recombinant full length SP140. **B,** Left, Schematic of topoisomerase 2 (TOP2) activity assay using migration and quantification of catenated (ct) and decatenated (dct) DNA following incubation of TOP2A with catenated DNA. Right, representative image and quantification of recombinant TOP2A (8 activity units, ~110ng) activity in the presence of indicated concentrations of recombinant full length SP140. Linear DNA (lin) serves as an assay control. **C,** Representative gel image and quantification of TOP1 activity assay in nuclear lysates from control (black bars) or siRNA-mediated SP140 knockdown (KD, red bars) naive or LPS (100 ng/mL, 4h)-stimulated THP1 monocytes. **D,** Representative gel image and quantification of TOP1 activity assay in nuclear lysates from control (black circles) or SP140 KD (red circles) naive or LPS (100 ng/mL, 4h)-stimulated primary human peripheral blood-derived macrophages. Connecting lines indicate individual healthy blood donors. **E,** Representative gel image and quantification of TOP1 activity assay in nuclear lysates from HEK293T cells transfected with indicated concentrations of FLAG empty vector (EV, black bars) or FLAG-SP140 (blue bars) or treated with Topotecan (TPT, 100μM). **F,** Representative gel image and quantification of TOP1 activity assay in lymphoblastoid cell lines (LBL) bearing wild-type SP140 (SP140^wt^) or CD-risk SP140 genetic variants (SP140^mut^) in the presence or absence of TPT (100μM). Data are mean of 3-4 biological replicates. Immunoblot of SP140 and TOP1 in control or SP140 siRNA-mediated knockdown **G,** THP1 cells or **H,** primary human macrophage nuclear lysates. **I,** Immunoblot of FLAG and TOP1 in HEK293T nuclear lysates transfected with indicated concentrations of FLAG empty vector (EV) and FLAG-SP140 (μg). **J,** Immunoblot of SP140 and TOP1 in lymphoblastoid cell lines (LBL) with wild-type SP140 (SP140^wt^) or Crohn’s disease (CD)-risk SP140 mutations (SP140^mut^). Lamin B is loading control. Data are representative of two independent experiments. Errors bars are s.e.m. \**P*< 0.05, \*\**P*<0.01, \*\*\**P*<0.001; two-tailed, unpaired t test.

### Loss of SP140 increases DNA breaks preferentially within heterochromatin

Recruitment of TOP1 and TOP2 at gene regulatory elements leads to transcriptional associated DNA breaks that activates DNA damage response pathways (Puc et al., 2017). Certainly, TOP-mediated double-stranded breaks (DSBs) are required for transcription initiation and elongation (Bunch et al., 2015; Ju et al., 2006). We therefore assessed levels of phosphorylation of the Serine 139 residue of the histone variant H2AX (gamma-H2AX, γH2AX), as a specific and sensitive molecular marker of DSBs and transcriptional activation. CRISPR-mediated deletion of Sp140 in Cas9 transgenic mouse macrophages or siRNA-mediated knockdown of SP140 in THP1 human monocytes resulted in significantly higher levels of γH2AX (**Figure 3A,B)**. The enhanced γH2AX observed in SP140 knockdown cells were not due to differences in DNA DSBs from canonical DNA damage as cell viability, cell cycle phases and levels of phosphorylated check point kinase (CHK2), were unaffected by SP140 depletion (**Figure S5, Related to Figure 3**), consistent with previous findings (Kim et al., 2019). Moreover, patient derived LBLs bearing disease-associated SP140 loss-of-function variants displayed significantly increased γH2AX, which was concomitant with the amount of SP140 protein loss (**Figure 3C)**. We next asked if gained γH2AX in the absence of SP140 was preferentially occurring at facultative heterochromatin marked by H3K27me3, where SP140 chiefly resides (Mehta et al., 2017). Indeed, confocal microscopy of murine or human macrophages revealed that elevated γH2AX primarily occurred at the nuclear periphery upon SP140 deletion or knockdown (**Figure 3D,E)**, where compacted H3K27me3+ facultative heterochromatin and low transcription activity is located (Buchwalter et al., 2019). Moreover, Chromatin Immunoprecipitation (ChIP) of γH2AX in primary human macrophages found a preferential gain of γH2AX at normally silenced loci *HOXA7*, *HOXB*9 and *FOXB1*, with unaltered γH2AX occupancy at the constitutively transcribed housekeeping gene, β-actin (*ACTB*) following siRNA-mediated knockdown of SP140 (**Figure 3F**). Thus, in the absence of SP140, transcriptional breaks and genome instability occur at normally silenced regions. To assess if levels of H3K27me3 influenced SP140 and TOP interactions, we treated macrophages with GSK343, an inhibitor of the H3K27me3 ‘writer’ Polycomb Repressive Complex 2 Subunit Enhancer of Zeste 2 (EZH2) and found that interactions between SP140 and TOP1 or TOP2A were diminished (**Figure 3G**) when H3K27me3 was reduced.

**Figure 3.**
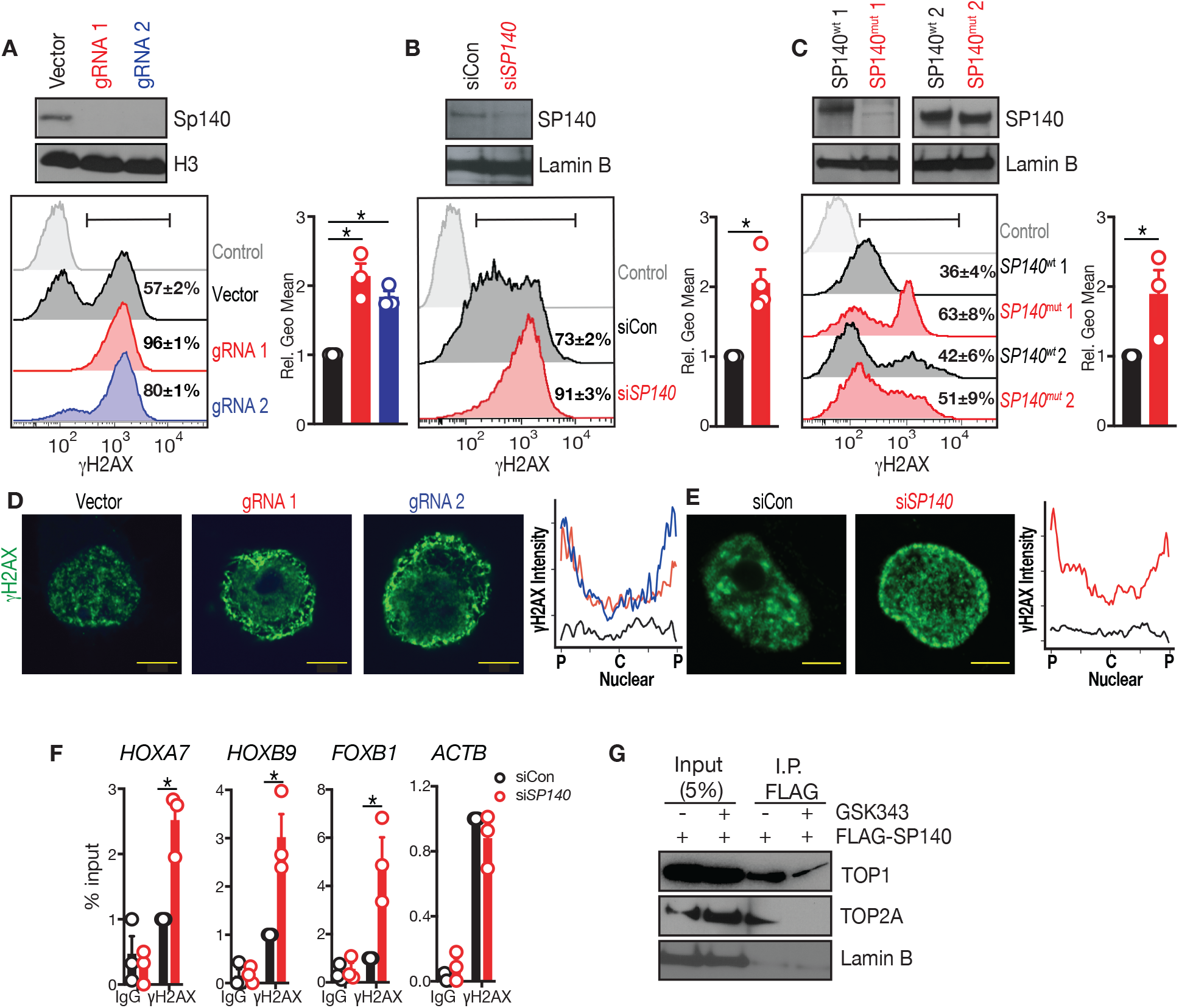
Loss-of-function SP140 or SP140 deletion increases DNA double strand breaks on heterochromatin. Quantification of DNA double stranded breaks as assessed by γH2AX (phospho S139) in **A,** Vector control or Sp140 CRISPR deleted Sp140 Cas9 transgenic immortalized mouse bone marrow macrophages (BMDMs) using two separate guide (g)RNAs **B,** control or *SP140* siRNA-mediated knockdown THP1 cells or **C,** race- and sex-matched lymphoblastoid B cell lines (LBL) bearing wild-type SP140 (SP140^wt^) or SP140 disease-risk mutations (SP140^mut^) by Flow Cytometry. Mean and s.e.m. γH2AX% positive cells are indicated in histogram gates, adjacent graphs are (γ)H2AX geometric mean relative to control. **D,** Cell localization of γH2AX in vector control or Sp140 CRISPR deleted Sp140 Cas9 transgenic immortalized mouse bone marrow macrophages (BMDMs) or **E,** control or *SP140* siRNA-mediated knockdown THP1 cells as assessed by confocal microscopy. Scale bars are 5μm. Line plots are the average quantification of γH2AX fluorescence intensity across the nuclear diameter (p=periphery, c=centre) of 25 cells using ImageJ. **F,** γH2AX Chromatin Immunoprecipitation quantitative PCR (ChIP qPCR) of *HOXA7, HOXB9, FOXB1* or *ACTB* in control (black) and SP140 knockdown (red) primary human macrophages, represented as % of input and normalized to siCon of each blood donor. IgG ChIP served as a negative control. **G,** Co-immunoprecipitation of FLAG SP140 and endogenous TOP1 and TOP2A in HEK293T cells treated with GSK343 (2.5μM).

### Gained TOP 1 and 2 occupancy at heterochromatin upon SP140 loss

TOP1 and TOP2A genome-wide occupancy in primary human macrophages, as assessed by Cleavage Under Targets and Tagmentation (CUT&Tag), revealed a preferential gain of both TOP1 and TOP2 at H3K27me3 rich regions upon knockdown of SP140 (**Figure 4A,B**) such as normally silenced loci *HOXA7* and *PAX5* (**Figure 4C**). Furthermore, Gene Ontology Biological Processes and Epigenomics Roadmap signatures of the loci with enhanced TOP1 and TOP2A occupancy upon SP140 knockdown reflected those regulating cell fate and identity and associated with markers of heterochromatin, H3K27me3 and H3K9me3 (**Figure 4D-G**). Collectively, this data reveals SP140 acts as gatekeeper of TOP1 and TOP2 activity that can occupy transcriptionally silent regions (Baranello et al., 2016) and in the case of TOP2, act to resolve facultative heterochromatin structure (Miller et al., 2017). Loss of SP140 leads to enhanced topoisomerase activity, aberrant chromatin accessibility (Mehta et al., 2017) and increased transcriptional activity and/or transcriptional potential at these normally silenced regions.

**Figure 4.**
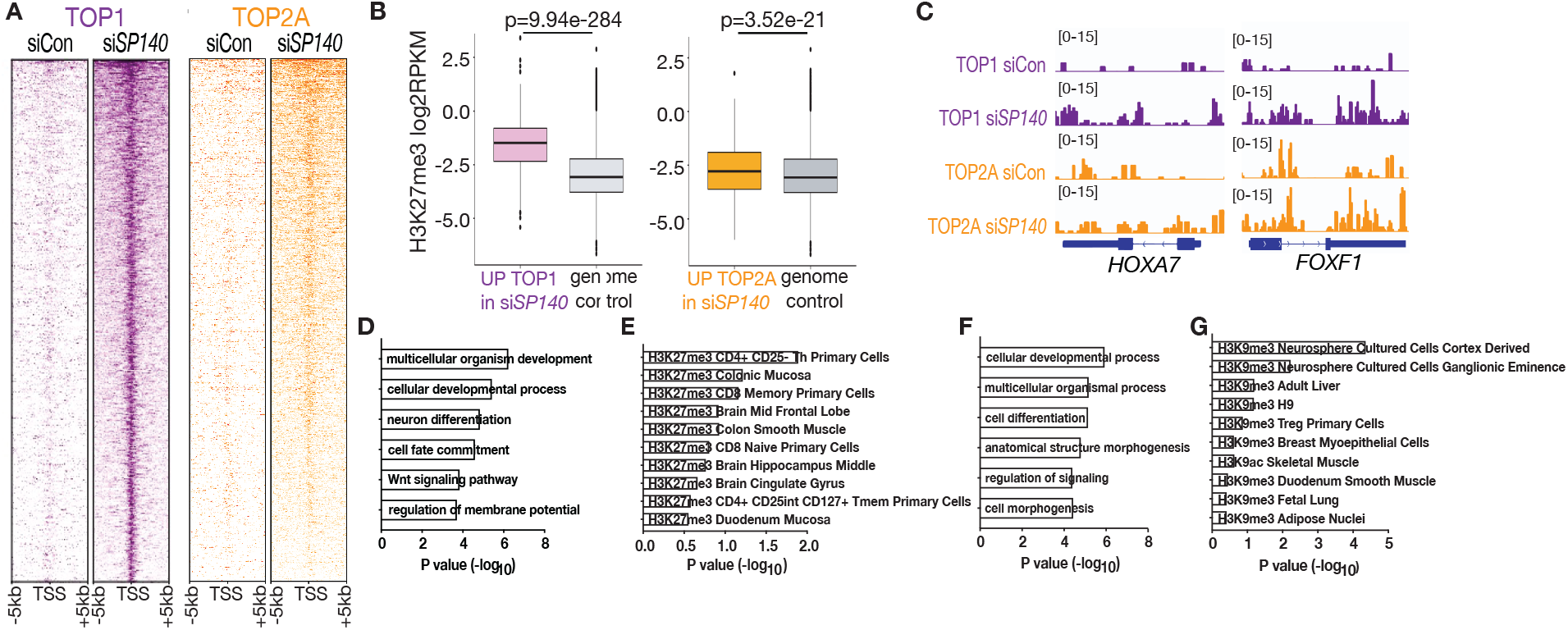
Gained occupancy of Topoisomerase 1 and 2 preferentially at heterochromatin regions upon SP140 loss. **A,** Heatmaps of TOP1 (purple) or TOP2 (orange) CUT&Tag peaks in transcriptional start site (TSS) proximal regions (TSS±5kb) in control or *SP140*-knockdown primary human macrophages rank-ordered by occupancy in control. **B,** H3K27me3 density (log_2_ scale), as determined by Multiplexed Indexed Chromatin Immunoprecipitation (MintChIP) at TSS-proximal regions (TSS±3kb) of genes that significantly increased TOP1 (purple) or TOP2 (orange) occupancy in SP140 knockdown human macrophages compared to genomic control (grey). RPKM, reads per kilobase, per million mapped reads. **C,** CUT&Tag genomic tracks of TOP1 (purple) and TOP2A (orange) densities at *HOXA7* and *FOXF1* in control and SP140 knockdown primary human macrophages. **D,** Gene Ontology Biological Processes or **E,** Epigenomics Roadmap Signature of loci that gained TOP1 occupancy in SP140 knockdown primary human macrophages (Fold Change>2, FDR<0.01) as assessed by CUT&Tag. **F,** Gene Ontology Biological Processes or **G,** Epigenomics Roadmap Signature of genes that gained TOP2A occupancy in SP140 knockdown primary human macrophages (Fold Change>2, FDR<0.01) as assessed by CUT&Tag. \**P*< 0.05. Data are mean of 3-4 biological replicates. Error bars are s.e.m. \**P*< 0.05; two-tailed, unpaired t test.

### TOP inhibitors rescue defective macrophage cytokine production and bacterial killing upon SP140 knockdown

The *ex vivo* reversal of topoisomerase activity with TPT provided proof-of-concept for use of topoisomerase inhibitors in SP140-deficient cells. We therefore next explored the use of TOP inhibition to rescue defective innate immune transcriptional programs driven by loss of SP140 (Mehta et al., 2017) and a key driver of disrupted immune-microbiome interactions in inflammatory bowel disease (Blander et al., 2017; Graham and Xavier, 2020). TOP1 inhibitor TPT is used clinically for treatment of small cell lung cancer and cervical cancer (Horita et al., 2015), while ETO is approved for treatment of refractory testicular tumour and small cell lung cancer (Loehrer, 1991). However, lower doses of TOP inhibitors can modulate microbe-inducible gene transcription and inflammation *in vivo* (Rialdi et al., 2016) and Aire-dependent immunological tolerance (Bansal et al., 2017). Treatment of SP140 knockdown primary human macrophages with TPT or ETO reduced inappropriate overexpression of *HOXA9* and *PAX5* at baseline (**Figure 5A-D**) and also rescued impaired transcription of cytokines *IL6* or *IL12B* in response to LPS (**Figure 5E,F**). In addition, administration of topoisomerase inhibitors rescued impaired antimicrobial activity of human SP140 knockdown macrophages toward Crohn’s disease associated adherent invasive E. *coli*, C. *rodentium* or S. *typhimurium* as assessed by gentamicin protection assays (**Fig 5G-I**).

**Figure 5.**
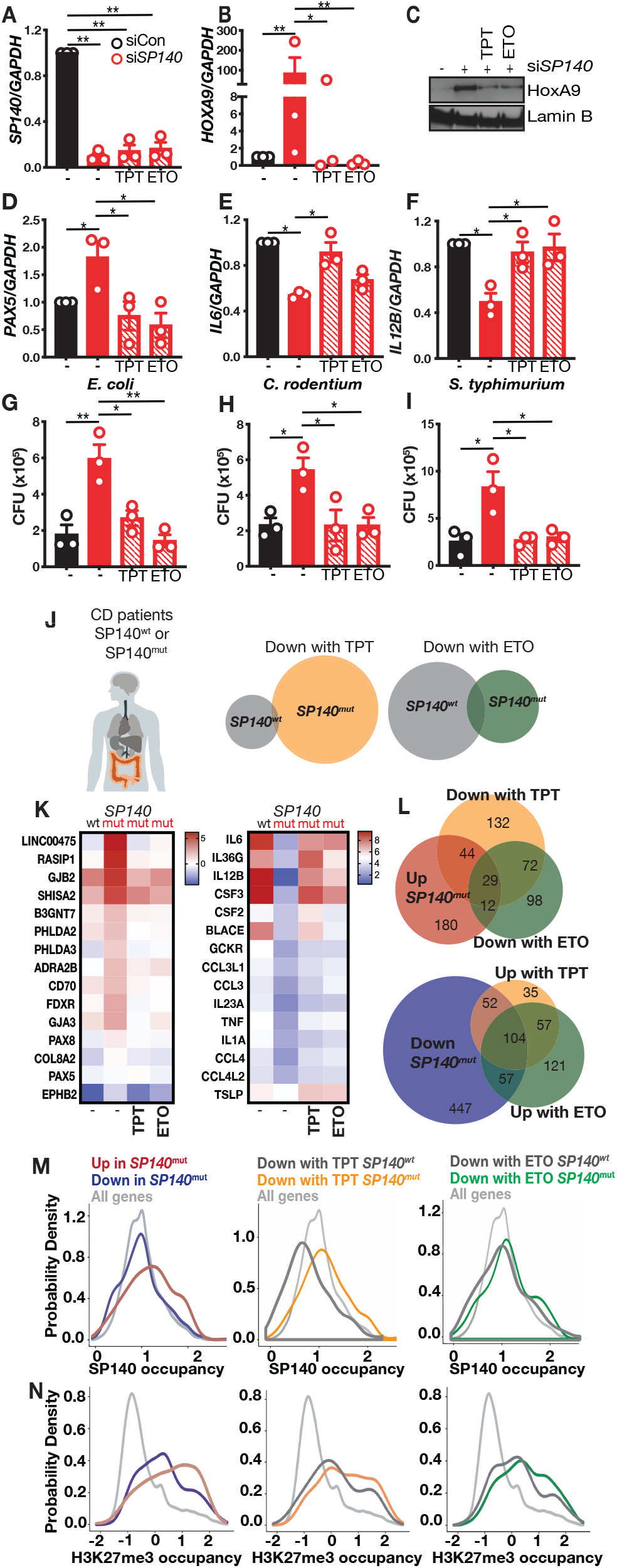
TOP inhibitors rescue defective innate immune transcriptional programs in SP140 loss-of-function Crohn’s disease patients. **A,** Levels of *SP140*, **B, C,** *HOXA9*, **D,** *PAX5*, **E,** *IL6* and **F,** *IL12B* in control (black) and SP140 knockdown (red) primary human macrophages from healthy donors in the absence or presence of topoisomerase I inhibitor, Topotecan (TPT, 100nM) or topoisomerase II inhibitor, Etoposide (ETO, 25 μM) as assessed by quantitative PCR or Western Blot. Data is normalized to DMSO siControl of each human blood donor. Lamin B is loading control. Data are mean of three biological replicates. Error bars are s.e.m. \**P*< 0.05, \*\**P*<0.01; one-way ANOVA with Tukey’s multiple comparisons test. G, Gentamicin protection assay on control or SP140 siRNA-mediated knockdown primary human macrophages treated with TPT (100 nM) or ETO (25 μM) and spin-infected with Crohn’s disease-associated *Escherichia coli* (E coli, 10 CFU/cell), **H**, Citrobacter rodentium (C rodentium, 10 CFU/cell), or **I**, *Salmonella enterica* serovar typhimurium (*S typhimurium,* 10 CFU/cell). Each dot is the average of a technical triplicate per individual blood donor. Error bars are s.e.m. \**P*< 0.05, \*\**P*<0.01; one-way ANOVA with Tukey’s multiple comparisons test. **J,** Fresh peripheral blood mononuclear cells (PBMCs) were obtained from CD patients carrying wild-type SP140 (SP140^wt^) or were homozygous for SP140 disease-associated mutations (SP140^mut^). Venn diagram of downregulated genes (fold change, FC>2) in CD patient SP140^wt^ (grey) or SP140^mut^ PBMCs by Topotecan (TPT, 100nM, orange) or Etoposide (ETO, 25μM, green) as determined by RNA-seq. K, Heatmap of expression changes (log_2_ FC) among top 15 differentially expressed genes in SP140^mut^ compared to SP140^wt^ that were rescued with TPT or ETO in LPS (100ng/mL, 4h)-stimulated PBMCs. L, Venn diagram of up- or down-regulated genes (log_2_FC >1) in CD patient SP140^mut^ PBMCs compared to SP140^wt^ which were reversed by log_2_FC>1 with TPT or ETO. M, Distribution (probability density function) of levels of SP140 enrichment (human macrophage ChIP-seq^12^) or **N,** H3K27me3 (average of ENCODE data ENCSR553XBX, ENCSR866UQO, ENCSR390SFH) in the TSS-proximal region for all genes (grey) compared with genes that were up-regulated (red) or down-regulated (blue) in SP140^mut^ PBMCs compared to SP140^wt^ PBMCs or TPT-(orange) or ETO-downregulated (green) genes in SP140^wt^ or SP140^mut^ PBMCs.

### TOP inhibitors rescue defective innate immune transcriptional programs in SP140 loss-of-function Crohn’s disease patients

We next exposed peripheral blood mononuclear cells (PBMCs) taken directly from Crohn’s disease (CD) patients bearing wild-type or CD-associated SP140 mutations to TPT or ETO. Fitting with enhanced TOP1 activity, more genes were responsive to TPT activity in SP140 loss-of-function cells than wild-type cells (**Figure 5G**), likely reflecting a new chromatin signature susceptible to TOP inhibition (Rialdi et al., 2016) upon loss of SP140. TPT or ETO delivery suppressed inappropriate upregulation of multiple genes in SP140 mutant PBMCs including *LINC000475, RASIP1, PHLDA2/3, PAX5/8* but importantly, also restored expression of inducible cytokines including *IL6, IL36G, IL12B, IL23A, TNF* and *IL1A* back to wild-type levels (**Figure 5H**). In fact, TPT or ETO rescued expression of ~50% or ~40% of aberrantly upregulated or downregulated genes in SP140 mutant CD patient PBMCs, respectively (**Figure 5I**). Importantly, the upregulated lineage-inappropriate genes in SP140 mutant PBMCs were those genes with a higher likelihood of being normally occupied by SP140 (**Figure 5J**) (Mehta et al., 2017). However, downregulated genes such as inducible cytokines were less likely to be occupied by SP140 suggesting an indirect effect of SP140 loss-of-function on these genes (**Figure 5J**). Furthermore, those genes whose irregular expression was rescued by TPT or ETO in CD SP140 mutant PBMCs were more likely to be normally occupied by SP140 (**Figure 5J**). Similarly, TPT or ETO target genes in SP140 mutant PBMCs were also more likely to reside in heterochromatin marked by H3K27me3 (**Figure 5K**). Consistently, unchecked topoisomerase activity and rescue of this activity in SP140 mutant PBMCs from CD patients with TOP inhibition occurs at loci normally occupied and repressed by SP140 within facultative heterochromatin.

### Precision rescue of exacerbated intestinal inflammation due to Sp140 loss with TOP inhibitors

Defective innate immunity and bacterial killing by resident tissue macrophages is a hallmark feature of Inflammatory Bowel Disease ((Graham and Xavier, 2020; Peloquin et al., 2016). We therefore examined the ability of TOP1 or TOP2 inhibition to rescue these macrophage defects for the treatment of inflammatory bowel disease associated specifically with loss of Sp140. We utilized Sp140 CRISPR-mediated knockout mice (Sp140^−/−^) and found that, similar to primary human macrophages, Sp140 deficient-bone marrow derived macrophages had impaired bacterial killing against E. *coli*, C. *rodentium* or S. *typhimurium* that was restored with addition of TPT, ETO or their combination (**Figure 6A-C**). We next assessed the ability of TOP inhibition to ameliorate DSS-colitis in Sp140 deficient mice *in vivo*. Sp140−/− mice had significantly exacerbated DSS-colitis compared to wild-type controls as measured by weight loss over time (**Figure 6D)** and histology (**Figure 6E**), consistent with previous data using shRNA-mediated Sp140 knockdown mice (Mehta et al., 2017). Indeed, intraperitoneal administration of single or combined TPT and ETO (1mg/kg each) delivered every other day significantly and specifically alleviated weight loss, rescued tissue architecture, edema and leukocyte infiltration and reverted colon length in Sp140^−/−^ mice (**Figure 6D-F**). In addition, markers of intestinal inflammation, including fecal lipocalin-2 and interleukin (IL)-6 and TNF in supernatants of colon explants of Sp140^−/−^ mice were all restored with topoisomerase inhibitor administration (**Figure 6G-I**). Notably, TOP1 or TOP2 inhibitors had no effect on the severity of DSS-colitis in wild-type mice (**Figure 6A-I**), further demonstrating an enhanced effectiveness of TOP inhibitors in the context of SP140 deficiency.

**Figure 6.**
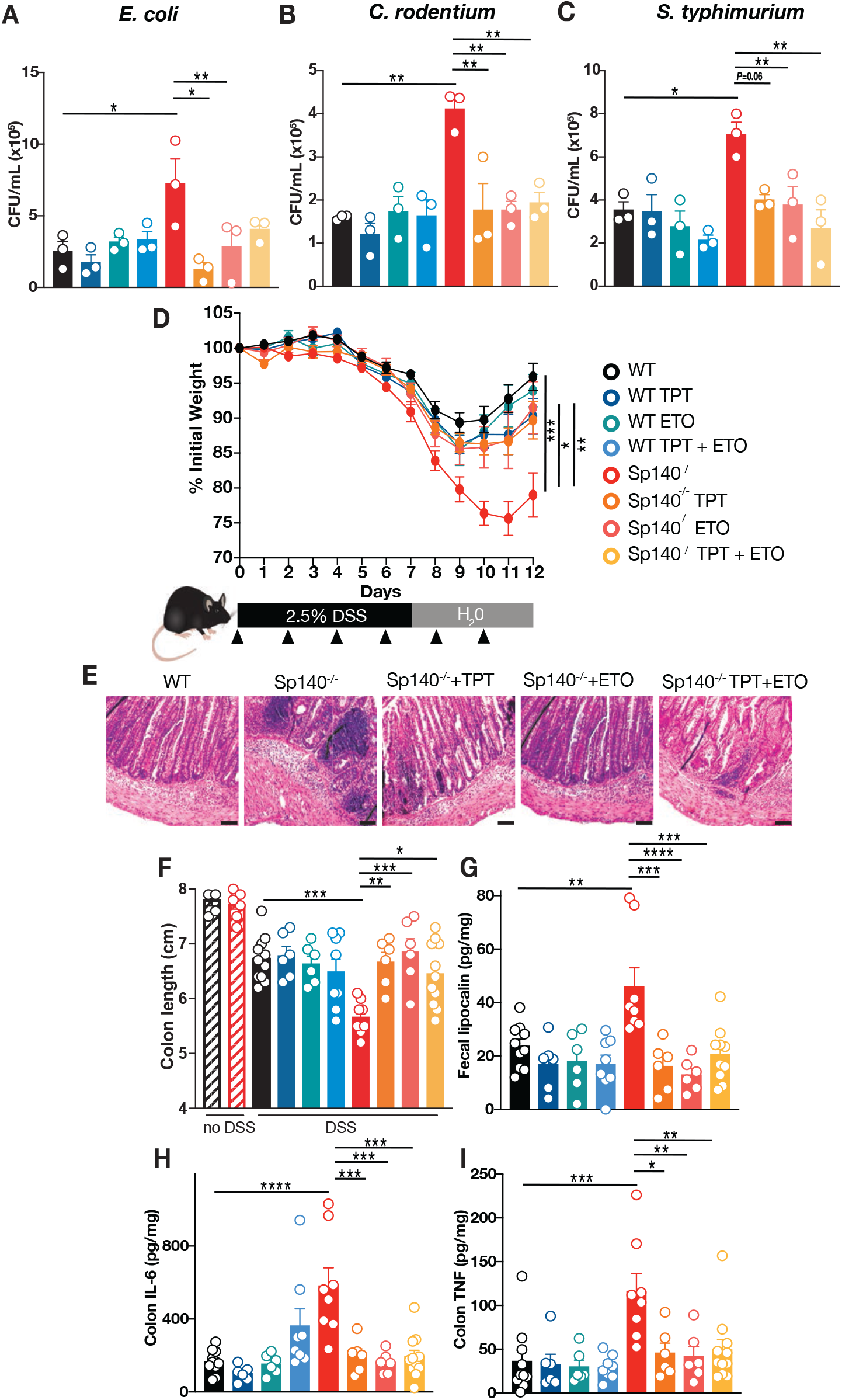
Precision rescue of DSS colitis in Sp140 knockout mice with TOP inhibition. **A,** Gentamicin protection assay using wild-type (WT, black) or SP140 CRISPR-deleted (Sp140^−/−^, red) bone marrow-derived macrophages treated with TPT (100 nM) or ETO (25 μM) and spin-infected with Crohn’s disease-associated *Escherichia coli* (E coli, 10 CFU/cell), **B**, Citrobacter rodentium (*C rodentium*, 10 CFU/cell), or **C**, *Salmonella enterica* serovar typhimurium (*S typhimurium,* 10 CFU/cell). Each dot is the average of a technical triplicate per individual mouse. Error bars are s.e.m. \**P*< 0.05, \*\**P*<0.01; one-way ANOVA with Tukey’s multiple comparisons test. Daily body weight measurements in WT and Sp140−/− mice after 2.5% DSS administration in drinking water for 7 days followed by water and intraperitoneal (i.p.) administration of DMSO control, TPT (1mg/kg), ETO (1mg/kg) or TPT+ETO (1mg/kg+1mg/kg) every other day as indicated with black arrow. **B**, Representative hematoxylin and eosin (H&E) staining of distal colon sections of WT and Sp140^−/−^ mice at day 12. Scale bar is 100μm. **C,** Colon length at baseline or day 12 of DSS. **D,** Fecal lipocalin-2 (Lcn-2) content at day 9 of DSS. **E,** Interleukin(IL)-6 or **F,** TNF levels in day 12 colonic explant supernatants cultured for 24 hours. \**P*< 0.05, \*\**P*<0.01, \*\*\**P*<0.001, \*\*\*\**P*<0.0001; one-way ANOVA with Tukey’s multiple comparisons test.

## Discussion

Mutations in chromatin reader SP140 are associated with at least three human immune-mediated diseases (International Multiple Sclerosis Genetics et al., 2013; Jostins et al., 2012; Sille et al., 2012), as well as *Mycobacterium tuberculosis* infection (Ji et al., 2021; Pan et al., 2005), directly implicating it in immunoregulation. However, insight into how SP140 functions in health or disease are limited. Here, we have identified a critical functional role for SP140 in the prevention of DNA accessibility and transcription by negative regulating topoisomerases at heterochromatin. This role of SP140 maintains gene silencing, immune cell identity and inducible innate immune transcriptional programs. Applying a combination of human genetics, proteomics, biochemistry, utilization of primary immune cells from Crohn’s disease individuals and *in vivo* animal studies, our study highlights the power of examining human-disease-associated mutations to advance mechanistic understanding of chromatin regulators in healthy and disease states. Furthermore, it has direct implications for precision-based therapies for Crohn’s disease patients bearing SP140 loss-of-function mutations, which to date have not been explored.

To gain molecular insight into SP140 biology, we characterized the protein complexes associated with SP140. At homeostasis, we found the predominant protein interactors of SP140 to be DNA unwinding and chromatin remodeling proteins including TOP1, TOP2A, TOP2B, DNA-PK, FACT components SUPT16H and SSRP1, as well as SMARCA5. We uncovered a prominent role of SP140 as a novel negative regulator of topoisomerases 1 and 2. Our previous studies revealed SP140 occupancy significantly correlated with a marker of facultative heterochromatin, H3K27me3, in primary human macrophages and loss of SP140 led to increased chromatin accessibility and expression of inappropriate lineage genes such as homeobox (*HOX)A9* that impair the functional response of differentiated macrophages (Mehta et al., 2017). Thus, we propose a model in which SP140, through its recognition of modified histones by its PHD and Bromodomain ‘reader’ modules, interacts with DNA unwinding and chromatin remodeling proteins such as topoisomerases to prevent chromatin remodeling and transcription at facultative heterochromatin. This function of SP140 maintains gene silencing and safeguards the differentiated immune cell state, and appropriate responses to environmental cues. Notably, this function of SP140, is in contrast to the well characterized and homologous chromatin reader, Aire which acts to promote transcription via topoisomerases (Abramson et al., 2010; Bansal et al., 2017). Moreover, SP140 is one of few endogenous proteins characterized to date that negatively regulate topoisomerases (Andersen et al., 2002; Kobayashi et al., 2009). Our findings further reinforce the importance of chromatin regulatory proteins in silencing lineage-specifying transcription factor genes, to maintain a functional cell state (Becker et al., 2016). Furthermore, we speculate that this SP140-dependent topoisomerase regulation to be a prominent requirement in immune cells, over other cell types in the human body, since they are uniquely mobile and highly responsive and often stably adaptive to environmental cues.

Importantly, this newly discovered mechanism of SP140 illuminates the indispensable role of chromatin-mediated gene silencing within malleable innate immune cells for lineage-defining and inducible transcriptional programs but also reveals a critical pathogenic mechanism in human immune-mediated disease. Crohn’s disease patients bearing SP140 loss-of-function variants lost essential SP140-TOP interactions resulting in unleashed TOP activity that derails normal innate immune cell-fate and inducible transcriptional programs. Importantly, chemical inhibition of TOP1 or TOP2 rescued the transcriptional defects in PBMCs taken directly from CD patients bearing SP140 loss-of-function mutations. Furthermore, precision rescue of Sp140^−/−^ mice with exacerbated DSS-colitis could be achieved with FDA-approved TOP inhibition *in vivo*. Future studies of TOP inhibition directly in SP140-mutant CD patients will be important for establishing the clinical impact of our work.

There is significant interest by the global research community in developing small molecule inhibitors of chromatin readers. However, a poorly understood feature of many chromatin ‘readers’ is how specificity of function is achieved through their interaction in multi-protein complexes and how this can be leveraged for therapeutic benefit when their loss-of-function drives disease. Moreover, while oncogenic outcomes are frequently reported (Filippakopoulos et al., 2010; Wan et al., 2020; Wen et al., 2014), human immune pathologies resulting from mutated chromatin regulators are less defined. Collectively, this work expands our knowledge of the types of human diseases that are caused by ‘misreading’ histone modifications. Furthermore, it reveals a mechanism-based strategy to rescue loss-of-function SP140 in human immune disease. Crohn’s disease remains incurable by surgical or therapeutic interventions so future studies may examine a precision-guided approach of FDA-approved TOP inhibitors for the treatment of CD, and potentially MS and CLL patients with loss of SP140, as well as examine potential roles of SP140 in metabolism and RNA processing.

## Limitations of the Study

Interactions between SP140 and chromatin remodeling and DNA unwinding proteins such as topoisomerases were lost in SP140 mutant immune cells but a gain of interactions between SP140 and multiple RNA processing related proteins was observed. These results indicate that loss of SP140 in immune disease does not just result in diminishment of the SP140-interactome, but rather a functional rewiring of the SP140 interactome is apparent. The importance of RNA processing and how it may influence RNA splicing and generation of alternative transcripts in SP140-driven disease remains an important avenue to be investigated further. Aire, similarly, complexes with RNA processing proteins and influences alternative splicing of its target genes (Abramson et al., 2010). Moreover, whether SP140 operates in a similar fashion in B cells and if unleashed TOP activity is a driver of CLL or MS upon SP140 loss-of-function warrants investigation. Future studies should also examine the role of other SP140-interactors identified in our Mass Spectrometry for regulation of transcription including chromatin remodeler SMARCA5 and the FACT complex. LPS-stimulated PBMCs from CD patients with wildtype or mutant SP140 were subjected to unbiased bulk-RNAseq and blinded bioinformatic analysis but sample size was limited to the MGH cohort. Additional experiments testing SP140 knockdown experiments were consistent with SP140 mutant data. Follow up work should expand to more CD patient numbers from other patient cohorts and to SP140 loss-of-function patients of other ethnicities and explored in other diseases associated with SP140 loss-of-function mutations.

## Acknowledgements

We thank the Taplin Mass Spectrometry Facility of Harvard Medical School, the Massachusetts General Hospital (MGH) Next Gen Sequencing core, the Broad Institute of Harvard and MIT Genomics Platform, the Harvard Stem Cell Institute CRM Flow Cytometry Core Facility and the Microscopy Core of the Program in Membrane Biology (PMB) at MGH. We also sincerely thank Stuti Mehta for initiating Mass Spectrometry experiments, Russell Vance (University of California, Berkeley) for Sp140−/− mice, Kate Fitzgerald (University of Massachusetts Medical School) for immortalized Cas9 transgenic bone marrow-derived mouse macrophages, Maya Kitaoka (Massachusetts General Hospital) for assistance with cloning, and the clinical coordinators and patients enrolled in the Prospective Registry in IBD study at Massachusetts General Hospital (PRISM). We also thank R.M. Anthony, R. Xavier, R.M. Anthony and R. Mostoslavsky for constructive comments. This study was supported by NIH R01DK119996 (K.L.J), Canadian Institutes of Health Research (CIHR) postdoctoral fellowship (H.A.) and K.L.J is a John Lawrence MGH Research Scholar, 2020-2025.

## Author contributions

H.A. contributed most experiments including proteomic analysis, co-immunoprecipitation experiments, mouse experiments, topoisomerase functional assays, ChIP, MintChIP and CUT&Tag experiments, developed methods, and generated resources, S.D. contributed co-immunoprecipitation experiments, CRISPR Cas9 mouse macrophages, primary human macrophage experiments and RNA-seq library preparation, I.F. and F.A contributed to endpoint measurements of mouse experiments, G.B., L.P.W, and R.S contributed RNAseq analysis, CUT&Tag analysis, MintChIP analysis and ChIP correlations, and K.L.J. conceived and supervised the study, obtained funding and wrote the final manuscript along with H.A.

## Competing interests

The authors declare no competing interests.

## STAR Methods

### LEAD CONTACT

Further information and requests for resources and reagents should be directed to and will be fulfilled by the Lead Contact, Kate L. Jeffrey

### MATERIALS AVAILABILITY

All unique/stable reagents generated in this study are available from the Lead Contact with a completed Materials Transfer Agreement.

### DATA AND CODE AVAILABILITY

RNA-seq, CUT&Tag, and MINT ChIP data have been deposited at GEO and are publicly available as of the date of publication. RNA-seq data have been deposited in the NCBI Gene Expression Omnibus (GEO) under accession GSE161031. TOP1 and TOP2A CUT&Tag data have been deposited in the NCBI Gene Expression Omnibus (GEO) under accession GSE174466. The H3K27me3 MINT ChIP data has been deposited in the NCBI Gene Expression Omnibus under accession GSE178632. There are no restrictions on data availability.

## EXPERIMENTAL MODEL AND SUBJECT DETAILS

### Human Subjects

Human Peripheral Blood Mononuclear Cells (PBMCs) were isolated from 20-30 mL of blood buffy coats from healthy human volunteers (Blood Components Lab, Massachusetts General Hospital) or from Crohn’s disease patients from the Prospective Registry in IBD study at Massachusetts General Hospital (PRISM) genotyped by CD-risk SP140 SNPs rs28445040 and rs6716753. Patient metadata is provided in **Extended Data Table 2**. Briefly, mononuclear cells were isolated by density gradient centrifugation of PBS-diluted buffy coat/blood (1:2) over Ficoll-Paque Plus (GE Healthcare). The PBMC layer was carefully removed and washed 3 times with PBS. In order to differentiate into macrophages, PBMCs were re-suspended in X-VIVO medium (Lonza) containing 1% penicillin/streptomycin (Gibco) and incubated at 37°C, 5% CO_2_ for 1h to adhere to the tissue culture dish. After 1h, adherent cells were washed 3 times with PBS and differentiated in complete X-VIVO medium containing 100 ng/mL human M-CSF (Peprotech) for 7 days at 37°C, 5% CO_2_. On day 4, cultures were supplemented with one volume of complete X-VIVO medium containing 100 ng/mL human M-CSF. 1 million PBMCs or mature macrophages were plated and pretreated with DMSO, topotecan (100nM) or etoposide (25μM) for 1 hour prior to 0.1mg/mL LPS treatment for 4 hours.

### Mice

All mice were housed in specific pathogen-free conditions according to the National Institutes of Health (NIH), and all animal experiments were conducted under protocols approved by the MGH Institutional Animal Care and Use Committee (IACUC), and in compliance with appropriate ethical regulations. For all experiments, age-matched mice Sp140 knockout or wildtype controls were used.

### Cells and treatments

HEK293T cells were maintained in DMEM media (Life Technologies) with 1% penicillin-streptomycin (Gibco) and 10% heat-inactivated Hyclone fetal bovine serum (FBS, GE Healthcare). THP-1 human monocytes were maintained in RPMI media (Life Technologies) with 1% penicillin-streptomycin (Gibco) and 10% heat-inactivated Hyclone fetal bovine serum (FBS, GE Healthcare).

## METHOD DETAILS

### Plasmids

Full-length cDNA encoding human SP140 was amplified from LPS-stimulated primary human peripheral blood derived macrophages. A plasmid expressing SP140 was made by cloning the purified SP140 cDNA into the p3xFLAG-CMV-10 vector (Addgene) using Gateway techniques (Invitrogen). SP140 mutants lacking DNA binding SAND domain (amino acids 580-661, SP140ΔSAND), or chromatin reading Plant homeobox domain (amino acids 690-736 PHD, SP140ΔPHD) or Bromodomain (amino acids 796-829, SP140ΔBromo) were generated by overlap-extension PCR using Phusion HF polymerase (Thermo Fisher Scientific). HA-Aire plasmid was obtained from Sun Hur (Boston Children’s Hospital, Harvard Medical School).

### siRNA-mediated knockdown in human macrophages and HEK293T

THP-1 human monocytes or mature primary human peripheral blood derived macrophages were transfected with control (D-001810-10-05, Dharmacon) or SP140 (L-016508-00-0005, Dharmacon) SMARTpool siRNA (100nM) using HiPerfect reagent (Qiagen) or RNAiMAX (Invitrogen) for 48 hours followed by 4 hour LPS treatment (0.1mg/mL). HEK293T cells were used for knockdown experiments with 100nM of Topoisomerase 1 siRNA (siGENOME SMARTpool Human TOP1, M-005278-00-0005, Dharmacon), Topoisomerase 2a siRNA (siGENOME SMARTpool Human TOP2A, M-004239-02-005, Dharmacon), DNA-PK siRNA (siGENOME SMARTpool Human PRKDC, M-005030-01-0005, Dharmacon) and Control (siGENOME Non-Targeting siRNA Pool #1, D-001206-13-05). All pooled siRNA sequences are available in **Extended Data Table 3**.

### Generation of CRISPR-mediated Sp140 null macrophages

Immortalized Cas9 overexpressing bone marrow-derived mouse macrophages were a kind gift from Dr. Kate Fitzgerald (University of Massachusetts Medical School). Mouse guide RNA sequences against SP140 were designed using Broad Institute’s sgRNA design algorithm (https://portals.broadinstitute.org/gpp/public/analysis-tools/sgrna-design). Guide 1 (TGTTGGGGAACATATGACAC) and Guide 2 (AAGGAAAAATTCAAACAAGG) sequences were cloned into lentiGuide-puro plasmid (Addgene, #52963), transformed in Stbl3 bacteria (Invitrogen, C7373-03), and subsequently purified with Qiagen Miniprep Kit. LentiGuide-Puro empty vector (control) or SP140 gRNA cloned plasmids were then co-transfected into HEK293T cells with the packaging plasmids pVSVg (AddGene 8454) and psPAX2 (AddGene 12260) for generation of lentivirus particles. Cas9 immortalized macrophages were transduced with lentivirus particles and 48 hours later treated with puromycin. Single cell clones were subsequently picked and grown.

### Quantitative PCR

RNA was extracted using the RNeasy Mini Kit (Qiagen) with on-column DNase digest (Qiagen) according to manufacturer’s instruction. 100ng-1μg RNA was used to synthesize cDNA by reverse transcription using the iScript cDNA Synthesis Kit (Bio-Rad). Quantitative PCR reactions were run with cDNA template in the presence of 0.625μM forward and reverse primer and 1x solution of iTaq Universal SYBR Green Supermix (Bio-Rad). Quantification of transcript was normalized to the indicated housekeeping gene. A complete list of primer sequences is provided in **Extended Data Table 4**.

### Mass Spectrometry

HEK293T cells were transfected with FLAG-Empty Vector (FLAG-EV) and FLAG-SP140 plasmids using Lipofectamine 2000 (Invitrogen) for 48 hours. Lysates were incubated with FLAG M2 (Sigma, F3165) antibody and immunoprecipitated protein was run on a 4-12% Tris-Bis. Gel and stained with Silver Stain (Thermo Fisher) and gel sections were cut and sent for in-gel digestion, micro-capillary LC/MS/MS, analysis, protein database searching and data analysis at the Taplin Mass Spectrometry Facility (Harvard Medical School). Mass Spec of lymphoblastoid cells (LBLs) carrying SP140 wildtype or SP140 loss-of-function variants (homozygous for rs7423615, Coriell Institute of Medical Research) was performed on endogenous SP140 immunoprecipitated from 1mg of nuclear lysates with SP140 antibody (Sigma).

### Nuclear fractionation

Cells were washed in PBS and gently lysed by resuspension in a freshly prepared and chilled sucrose lysis buffer (320 mM sucrose, 10 mM Tris pH 8.0, 3 mM CaCl2, 2mM magnesium acetate, 0.1 mM EDTA, 0.5% NP-40, 1% protease/phosphatase inhibitor cocktail). Cells were incubated in this buffer for 15 minutes on ice followed by centrifugation at 500*g* for 15 minutes at 4°C to pellet isolated nuclei. Supernatant containing the cytosolic fraction was removed and the remaining nuclear pellet was mixed in a nuclei sonication buffer (500 mM NaCl, 50 mM Tris pH 8.0, 10% glycerol, 1% NP-40, 1% protease/phosphatase inhibitor cocktail) followed by incubation on ice for 5 minutes. Next, the nuclei were lysed in this buffer by sonication for 10 seconds. Insoluble debris from this lysate was removed by centrifugation at 21000*g* for 20 minutes at 4°C and remaining supernatant was taken as nuclear fraction protein.

### Co-immunoprecipitation

HEK293T cells were seeded into 10-cm dishes and transfected with 25μg FLAG-Empty Vector (FLAG-EV) and FLAG-SP140 plasmids using Lipofectamine 2000 (Invitrogen). 48 hours after transfection, nuclear extracts were isolated from cells. Alternatively, nuclear extracts from 10 million THP1 monocyte or LBL cells were used for endogenous SP140 IPs. 500μg of nuclear lysate was incubated with antibody for pulldown overnight at 4°C. This was followed by addition of Protein G Dynabeads (Invitrogen) and incubation for 4 hours at 4°C with rotation. The bead slurry was then magnetized, the unbound fraction removed, and the beads subjected to a series of increasingly harsh salt-based washes. The beads were boiled at 90°C for 10 minutes in a reducing sample buffer to elute bound protein and the elution was used for probing of co-IP interactors by Western blotting. Eluted proteins were run on a 3-8% Tris Acetate Gel (Invitrogen) or 4-12% Tris Bis gel (Invitrogen), transferred onto PVDF membrane, blocked with 5% skim milk in TBS/0.1% Tween at room temperature for 1h, and then incubated with the indicated primary antibody in 3% BSA TBS/0.1% Tween overnight at 4°C. To assess DNA dependence of co-IP interactors, ethidium bromide was added at a concentration of 1mg/mL to 500μg of nuclear lysate and allowed to incubate for 30 minutes on ice prior to addition of antibody for pulldown as described previously. Salt-based wash solutions were also supplemented with 100μg/mL ethidium bromide. As before, beads were boiled and eluted protein was used for probing of interactors by Western blot.

### Topoisomerase activity assays

Full length human recombinant SP140 protein was commercially produced by BPS Bioscience using a HEK293T expression system. Recombinant Topoisomerase I and II were purchased from Topogen. Nuclear lysates were obtained from control or SP140 knockdown THP-1 cells or primary human peripheral blood-derived macrophages, HEK overexpressing FLAG SP140 or SP140 protein domain mutants, or lymphoblastoid cells (LBLs) carrying wildtype or SP140 genetic variants (homozygous for rs7423615) and Topoisomerase I activity was performed according to the manufacturer’s instructions (Topoisomerase I activity assay kit, TopoGen). Briefly, nuclear lysates were incubated with supercoiled plasmid (pHOT) DNA substrate, and reaction buffer (10mM Tris-HCl pH 7.9, 1mM EDTA, 0.15M NaCl, 0.1% BSA, 0.1mM Spermidine, 5% glycerol) for 30 minutes at 37°C. The reaction was stopped using a “Stop” Buffer (0.125% Bromophenol Blue, 25% glycerol, 5% Sarkosyl) and samples were treated with proteinase K (50μg/mL) and Ribonuclease A (Roche) for 30 minutes at 37°C to remove contaminating proteins and RNA. The reaction mixture, along with supercoiled and relaxed controls, were then run on a 1% agarose gel, stained with Ethidium Bromide for 20 minutes, de-stained in distilled water, and imaged for photodocumentation. Bands were quantified using Image Studio Lite (Licor Biosciences) and ratios were calculated with relaxed over supercoiled quantified data. Topoisomerase II activity assay was performed according to the manufacturer’s instructions (Topoisomerase II activity assay kit, TopoGen). Briefly, recombinant SP140 protein was incubated with recombinant TOP2A protein, catenated plasmid kinetoplast DNA (from insect C *fasciculate*) substrate, reaction buffer (0.5M Tris-HCl pH 8, 1.5M NaCl, 100mM MgCl2, 5mM Dithiothreitol, 300μg/mL BSA) for 30 minutes at 37°C. The reaction was stopped using a “Stop” Buffer (0.125% Bromophenol Blue, 25% glycerol, 5% Sarkosyl). The reaction mixture, along with catenated and decatenated circular and linear controls, were then run on a 1% Ethidium Bromide agarose gel, destained in distilled water, and imaged for photodocumentation. Bands were quantified using Image Studio Lite (Licor Biosciences) and ratios were calculated with decatenated relaxed and nicked DNA over catenated quantified data.

### Flow Cytometry

Macrophages were fixed as single cell suspensions in 4% formaldehyde. Fixed cells were then permeabilized in 0.25% Triton X-100 and blocked in 1% BSA/0.05% Tween-20 in PBS. Cells were then incubated with 1:500 γH2AX antibody (Abcam), followed by washing and collection in FACS buffer (1% FBS in PBS) for flow cytometry. Annexin V and propidium iodide for cell death analysis was performed according to manufacturer’s instructions (Alexa Fluor® 488 annexin V/Dead Cell Apoptosis Kit, Thermofisher). Cell cycle analysis was performed with ethanol fixed cells and stained in PBS with 50μg/mL Propidium Iodide and 100μg/mL RNase A. All flow cytometry data was collected on LSRII (BD) and analyzed with FlowJo (TreeStar).

### Immunofluorescence microscopy

Control or *SP140* siRNA-mediated knockdown THP1 or CRISPR-mediated Sp140 null mouse immortalized bone marrow-derived macrophages were fixed with 4% formaldehyde for 20 mins and subsequently permeabilized with 0.25% Triton X-100, blocked with 5% BSA/PBST and stained with 1:500 γH2AX antibody (Abcam) overnight at 4°C. The samples were subsequently washed the next day and incubated with secondary antibody at 1:1000 (AlexaFluor 488, Invitrogen) for 1 hour at room temperature, following which cells were washed and coverslips were mounted onto microscope slides with Prolong Diamond anti-fade mountant (Invitrogen). Multiple representative images were obtained on a confocal microscope (Zeiss LSM 800 Airyscan). Images were quantified using ImageJ by measuring fluorescence intensity across the diameter of a nucleus of 25 cells. Quantified data were binned to normalize different sized cells and plotted as a lineplot with Python showing average intensity from one periphery of the cell to the other periphery.

### Chromatin Immunoprecipitation (ChIP)

Ten million naïve or LPS-stimulated (0.1mg/mL for 4 hours) human peripheral blood-derived macrophages were cross-linked and processed using truChIP kit (Covaris Inc), as described in detail before(Mehta et al., 2017). Input fraction was saved (10%), and the remaining sheared chromatin was used for ChIP with γH2AX antibody (Abcam) or an immunoglobulin G isotype control (Abcam) in immunoprecipitation buffer (0.1% Triton X-100, 0.1 M Tris-HCl pH 8, 0.5 mM EDTA, and 0.15 M NaCl in 1Å~ Covaris D3 buffer) at 4°C, rotating overnight, followed by incubation with Dynabeads Protein G (Life Technologies) for 4 hours. The chromatin-bead-antibody complexes were then washed sequentially with three wash buffers of increasing salt concentrations and TE buffer. Chromatin was eluted using 1% SDS in Tris-EDTA. Cross-linking was reversed by incubation with ribonuclease A (Roche) for 1 hour at 37°C, followed by an overnight incubation at 65°C with Proteinase K (Roche). DNA was purified with the QIAquick PCR Purification Kit (Qiagen). DNA was used for ChIP qPCR using primers in Extended Table 4. ChIP data across different donors was normalized to γH2AX ChIP of control sample of each donor.

### RNA library prep, sequencing and analysis

RNA was extracted using the Qiagen RNeasy Mini Kit (with on-column DNase I digest) and samples were eluted in 40μL RNase-free water. Sample concentration and quality was measured using the HighSensitivity RNA TapeStation kit for the TapeStation Bioanalyzer (Agilent). After confirming the RNA was of sufficient quality using the RIN^e^ metric, 100ng of RNA was taken from each sample and made up to 50μL in RNase-free water. mRNA was enriched for and used to prepare libraries using the NEBnext Poly(A) mRNA magnetic isolation module (NEB E7490S) and NEBnext Ultra II RNA library prep kit (NEB E7770S). Briefly, poly(A)-tailed RNA was isolated by coupling with Oligo d(T) magnetic beads followed by a series of washes to remove unbound RNA. This coupling was repeated for a total of two selection steps. Finally, beads were resuspended in a mix of random primers and First Strand Synthesis Buffer and boiled for 10 minutes to fragment selected mRNA. First Strand Enzyme mix was added to the fragmented RNA-buffer elution and thermal cycled for preparation of cDNA. The second strand was synthesized in a similar manner, using the respective buffers and enzymes. Double-stranded cDNA was selected for using SPRI size selection magnetic beads, end prepped, and ligated with unique adapters for multiplexing. This ligation mixture was PCR cycled to amplify cDNA and libraries were purified using a series of SPRI size selection steps. Finally, library size and purity were confirmed using the High Sensitivity D1000 Tapestation kit on Tapestation Bioanalyzer. Multiplexed samples were submitted for paired-end sequencing on NovaSeq S2 (Genomics Platform, Broad Institute), resulting in 14-52 million mapped reads per sample. Reads were mapped using the STAR aligner (Dobin et al., 2013) and the hg19 assembly of the human genome. Read counts for individual transcripts were obtained using HTSeq (Anders et al., 2015) and the Ensembl gene annotation (GRCh37 release 75) (Yates et al., 2016). Differential expression analysis was performed using the EdgeR package (McCarthy et al., 2012), and differentially expressed genes were defined based on the criteria of >2-fold change in normalized expression value. For probability density distribution analysis of differential genes, ChIP signal enrichments at promoters were calculated by counting reads in the region defined by ±3 kb from the TSS. Read counts were normalized to reads per million (RPM), and the enrichment was calculated as the ratio of RPM in ChIP divided by the RPM in input chromatin (control) and was converted to log2 scale. The probability density distribution was computed and plotted with standard R package. SP140 ChIP-seq in primary human macrophages was previously generated in our lab (Mehta et al., 2017) and H3K27me3 analysis included ChIP-seq obtained in peripheral blood mononuclear cells (ENCODE accession: ENCSR553XBX, ENCSR866UQO, ENCSR390SFH. An average of these three studies was used).

### Multiplexed indexed T7 ChIP-seq (MintChIP)

To profile H3K27me3 levels in control primary human macrophages, we performed optimized version of Mint-ChIP3 (van Galen et al., 2016), as described before. Briefly, 100,000 cells were lysed and digested with 300 units of micrococcal nuclease (MNase, NEB) for 15 mins. T7 Adapter Ligation was subsequently performed and quenched. All samples were pooled and then equally split for primary antibody (H3K27me3 and H3 Antibody) overnight incubation at 4C. Protein G Dynabeads were added the next morning and incubated for 4 hours at 4°C. Following low salt RIPA, high salt RIPA, LiCl and TE washes, the beads were eluted and digested at 63°C for 1 hour. The beads were cleaned up using AMPure SPRI beads (Beckman Coulter). In vitro transcription of eluted DNA to RNA was performed using NEB HiScribe T7 kit for 2 hours and subsequently treated with DNase at 37°C for 15 mins. RNA was isolated using Silane beads (Thermofisher). RNA was reverse transcribed to cDNA and subsequently cleaned up with AMpure SPRI beds. The purified DNA was then used for Library PCR with Illumina barcoded primers to Enrich Adapter-Modified DNA Fragments. Samples were submitted for sequencing on HiSeq (Genomics Platform, Broad Institute). MINT ChIP reads were aligned to the hg19 genome using BWA version 0.7.15, technical replicates were pooled together. bigWig files were generated using the deepTools bamCoverage command, normalized using CPM, and with bin size of 50, and smooth length of 100. Normalized bigWig files were used to calculate intensities at promoters from the Ensembl gene annotation (GRCh37 release 75) using the deepTools multiBigwigSummary command, with a BED file obtained by extending the TSS by +/− 3000 bp.

### Cleavage Under Targets and Tagmentation (CUT&Tag)

CUT&Tag was performed according to Epicypher protocol. Briefly, nuclei were isolated from 100,000 primary human macrophages and bound to activated magnetic concanavalin A beads (Epicypher). The bound nuclei were incubated with primary antibody overnight at 4°C. The next day, the beads were washed and subsequently incubated with secondary antibody for 0.5 hour at room temperature. This was followed by washing and incubation with the protein A/G conjugated Tn5 (CUTANA pAG-Tn5, Epicypher) enzyme for one hour at room temperature. The enzyme bound beads were then washed, and then tagmentation reaction was performed for 1 hour at 37°C. The reaction was stopped with TAPS buffer containing EDTA, released with SDS buffer, and quenched with 0.67% Triton X-100 solution. The released DNA was cleaned up using Monarch DNA PCR Clean Kit (NEB) and eluted in TE buffer. Library was constructed using universal i5 and barcoded i7 primers and non-hot start CUTANA® High Fidelity 2x PCR Master Mix (Epicypher). DNA was cleaned and size selected using 1.3x AMPure beads and visualized on Agilent Bioanalyzer system. Samples were submitted for sequencing on the NovaSeq_SP_100 (Genomics Platform, Broad Institute). Paired-end sequencing reads were aligned to hg19 human reference genome using bwa version 0.7.17 (Li and Durbin, 2009). diffBind R package (Ross-Innes et al., 2012) was used for the analysis of differential regions between knockdown and control samples at promoter regions (+/− 3kb proximity of transcription start sites). Differential regions were determined based on the cutoffs of > 2-fold change of read density and false discovery rate (FDR) < 0.01. Heatmaps and metaplots of CUT&Tag signal densities were generated using deepTools (Ramirez et al., 2014).

### Gentamicin Protection Assay

Primary human macrophages or bone marrow derived macrophages (1× 10^6^) plated in 12-well plates were infected with live adherent invasive E. *coli*, C. *rodentium*, and S. *typhimurium* at a 1:10 with spin infection for 10 mins and incubation for 20 mins. Cells were washed three times with 1x PBS and provided medium containing 100 μg/mL gentamicin for 1 hour. Cells were washed again with 1x PBS and lysed with 0.1% Triton x-100 in 1x PBS following 1, 2 or 3 hours for E. *coli*, C. r*odentium* and S. *typhimurium*, respectively. Serially diluted cell lysates were plated on LB agar plates. Colony forming units (CFUs) were determined after 24 hours of incubation at 37°C with 5% CO_2_.

### Dextran Sodium Sulfate (DSS) colitis

All mice were housed in specific pathogen-free conditions according to the National Institutes of Health (NIH), and all animal experiments were conducted under protocols approved by the MGH Institutional Animal Care and Use Committee (IACUC), and in compliance with appropriate ethical regulations. For all experiments, age-matched mice Sp140 knockout or wildtype controls were randomized and allocated to experimental group, with 6-12 mice per group. No statistical method was used to determine sample size. Mice were administered 2.5% Dextran Sodium Sulfate Salt (DSS, MW = 36,000-50,000 Da; MP Biomedicals) in drinking water *ad libitum* for 7 days (freshly prepared every other day), followed by regular drinking water for 5 days. Mice were administered DMSO (control), topotecan (Sigma) or etoposide (Sigma) by intraperitoneal injection (i.p. 1mg/kg) starting on day 1 and administered every other day until mice were sacrificed on day 12. Colon length was measured and Day 9 fecal samples were homogenized in PBS, spun down and supernatants were collected and assayed for Lipocalin-2 by ELISA, according to manufacturer’s instructions (R&D). For cytokine measurements, day 12 colon tissue (excised approximately 1 cm) was washed in PBS and then placed in a 24 well plate containing 1mL of RPMI medium with 1% penicillin/streptomycin and incubated at 37°C with 5% CO2 for 24 hours. Supernatants were collected and centrifuged for 10 min at 4°C and IL-6 and TNF levels were assessed by ELISA, according to manufacturer’s instructions (R&D).

## QUANTIFICATION AND STATISTICAL ANALYSIS

Results are shown as mean ± s.e.m. Visual examination of the data distribution as well as normality testing demonstrated that all variables appeared to be normally distributed. Comparisons and statistical tests were performed as indicated in each figure legend. For comparisons of two groups, two-tailed unpaired t tests were used, except where indicated. For comparison of multiple groups, one-way ANOVA was used. Statistical analyses were performed in the GraphPad Prism 8 software. The *P* values denoted throughout the manuscript highlight biologically relevant comparisons. A *P* value of less than 0.05 was considered significant, denoted as \**P*<0.05, \*\**P*<0.01, \*\*\**P*<0.001, and \*\*\*\**P*<0.0001 for all analyses.

---

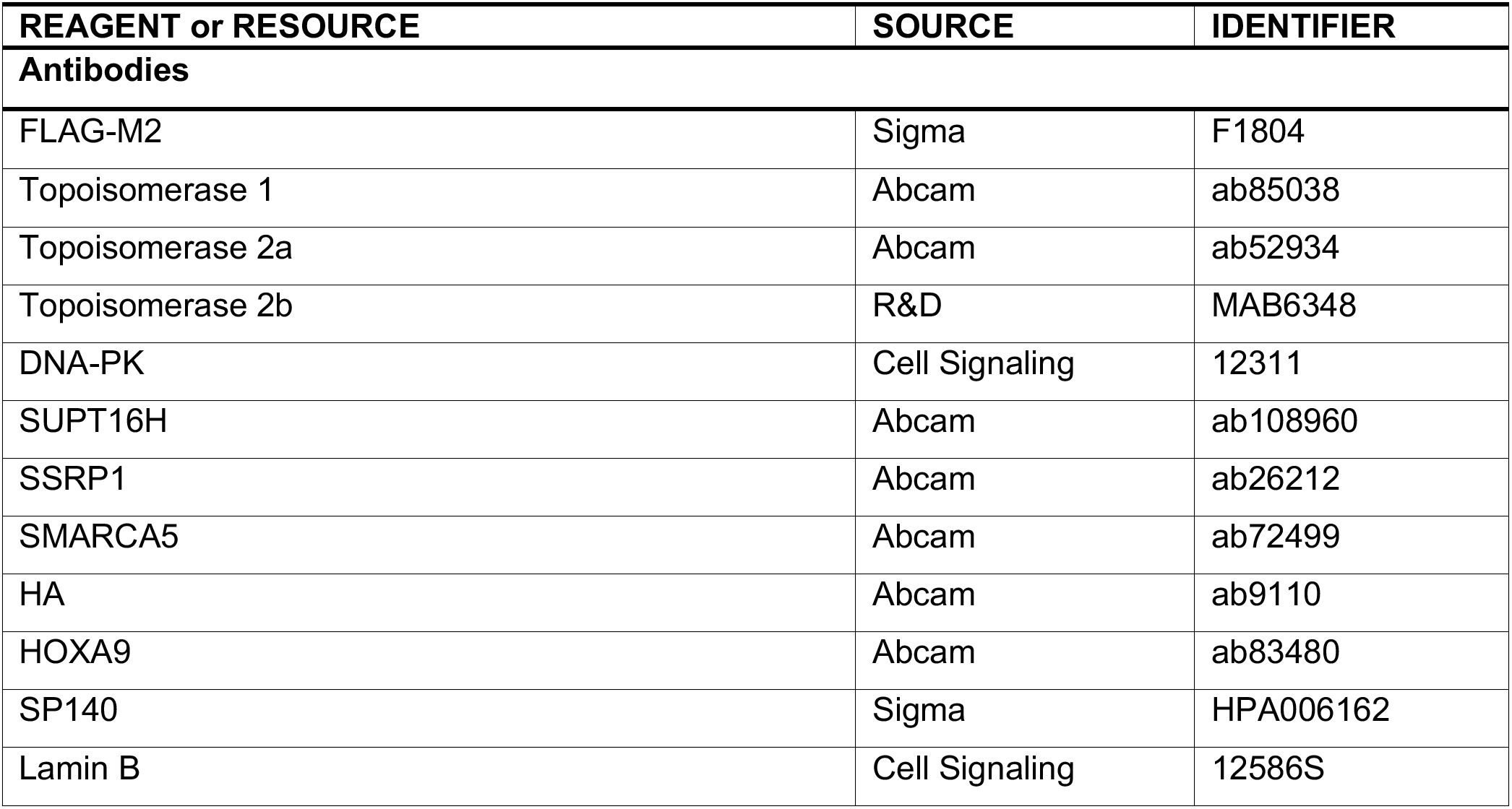
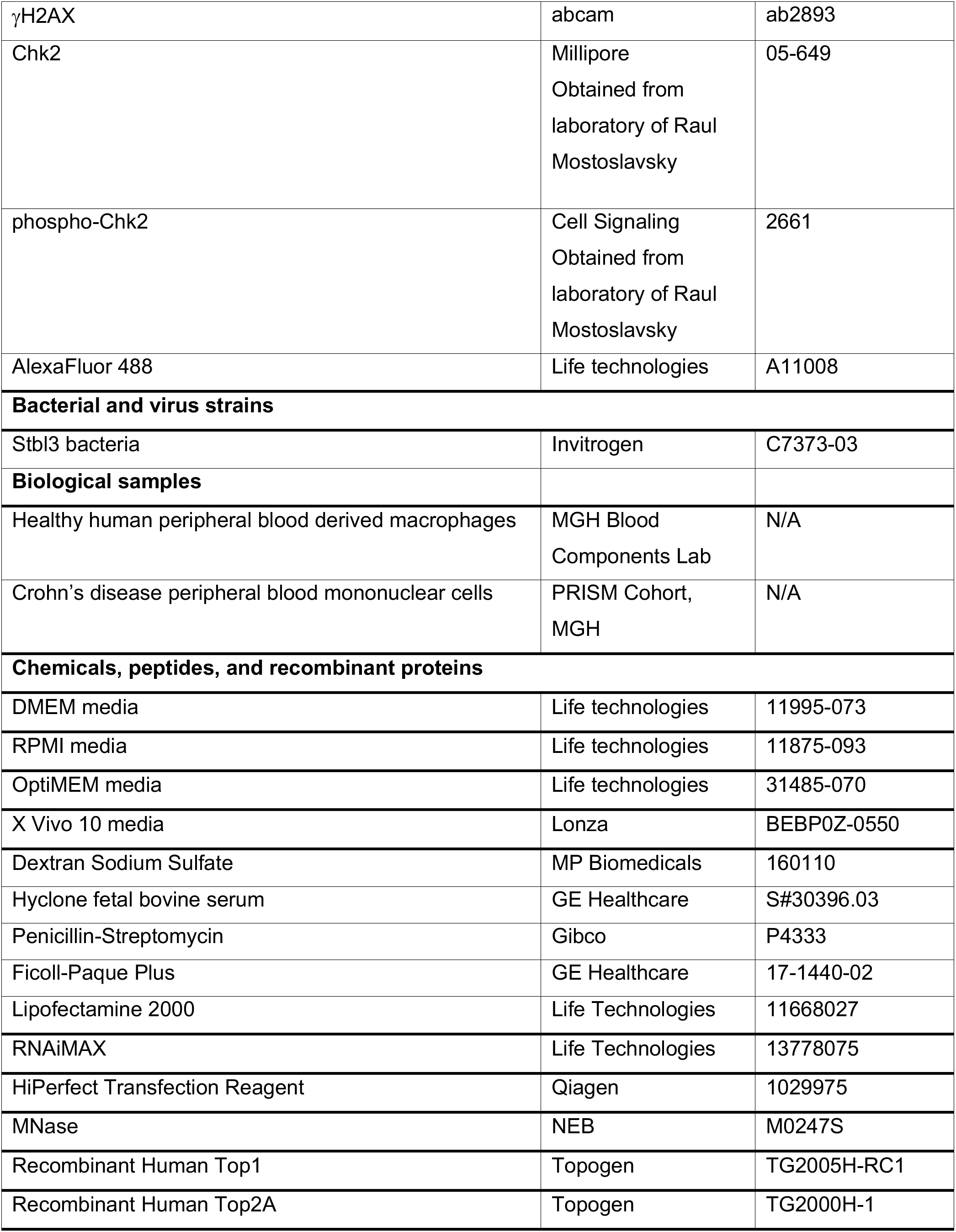
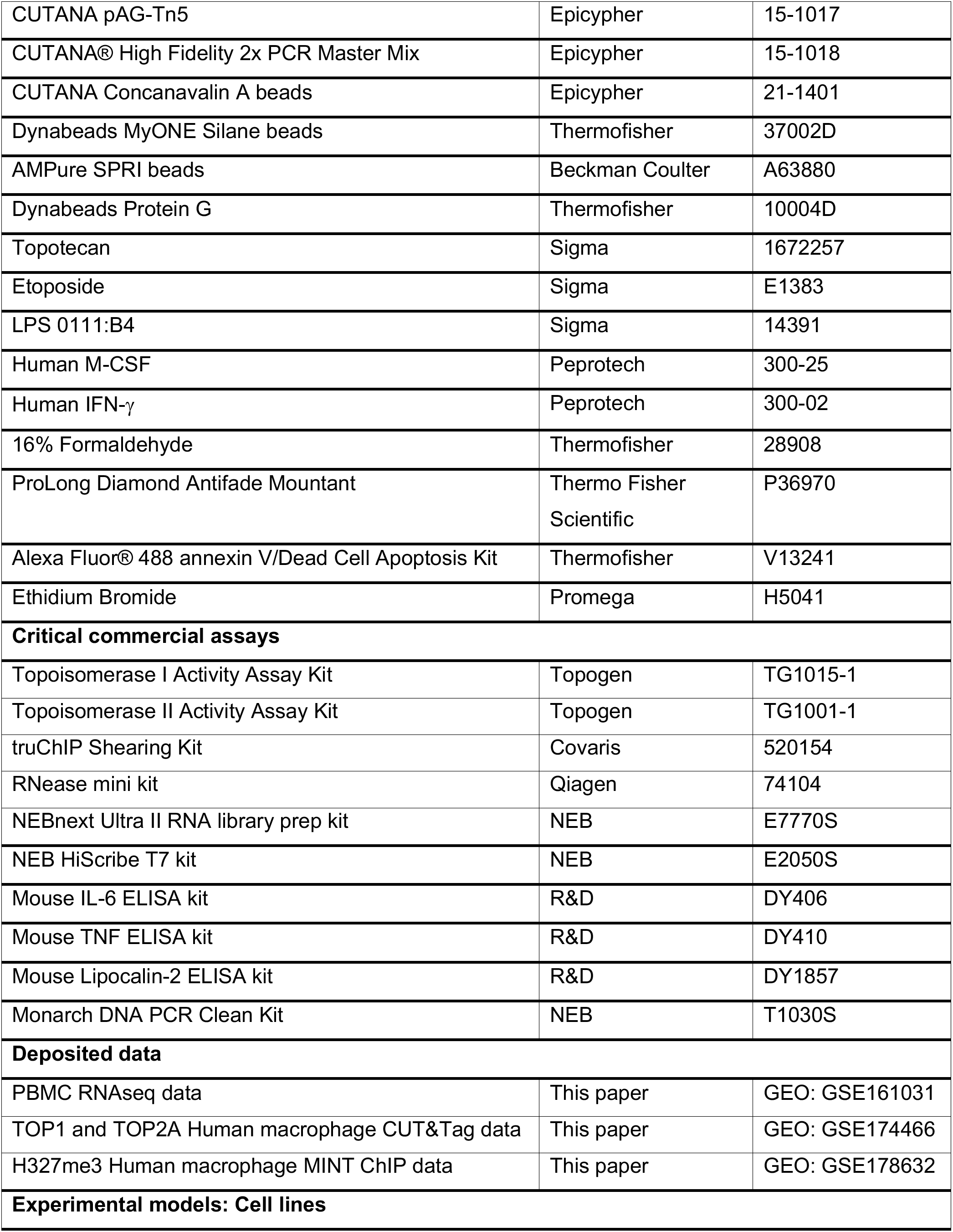
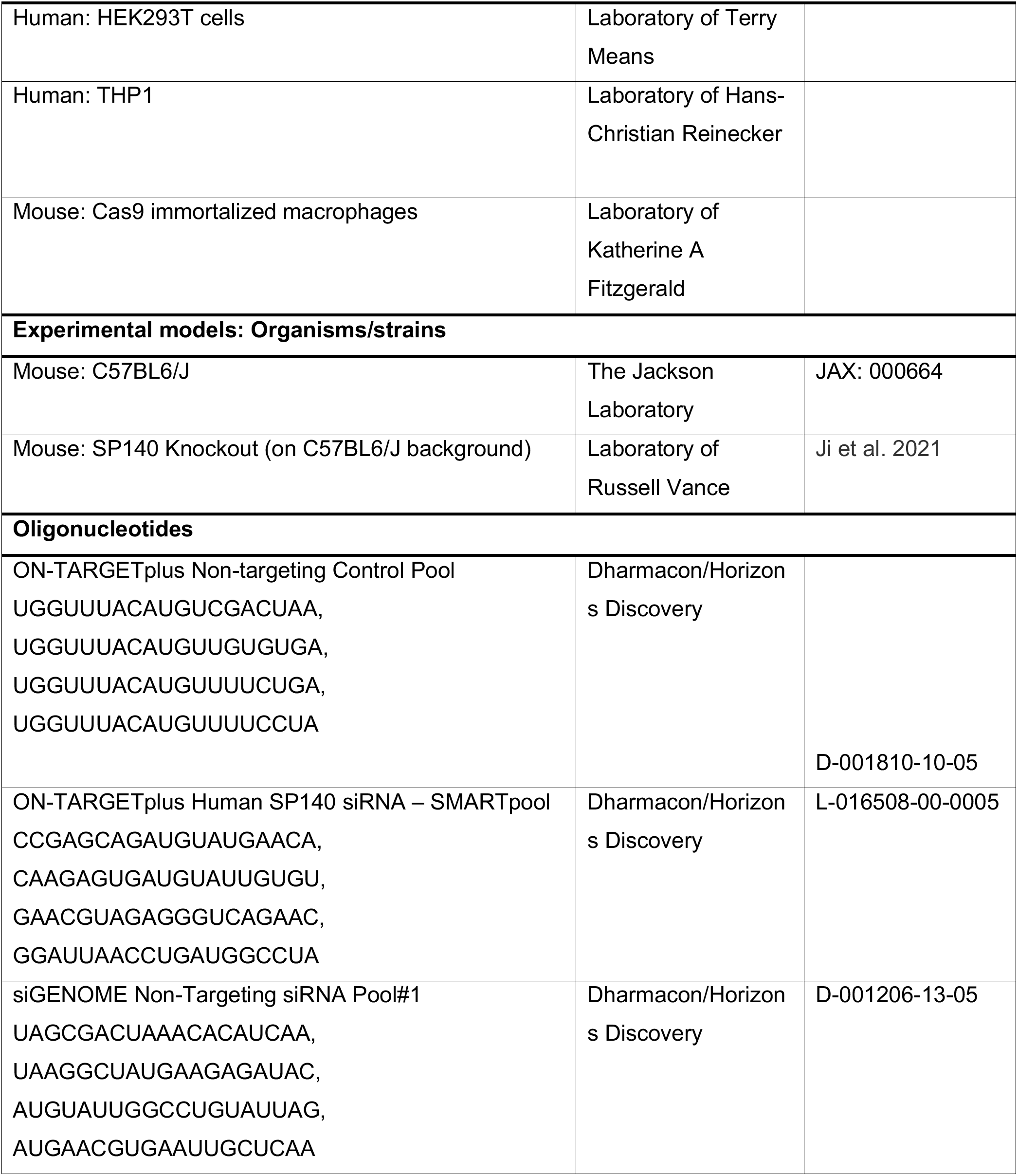
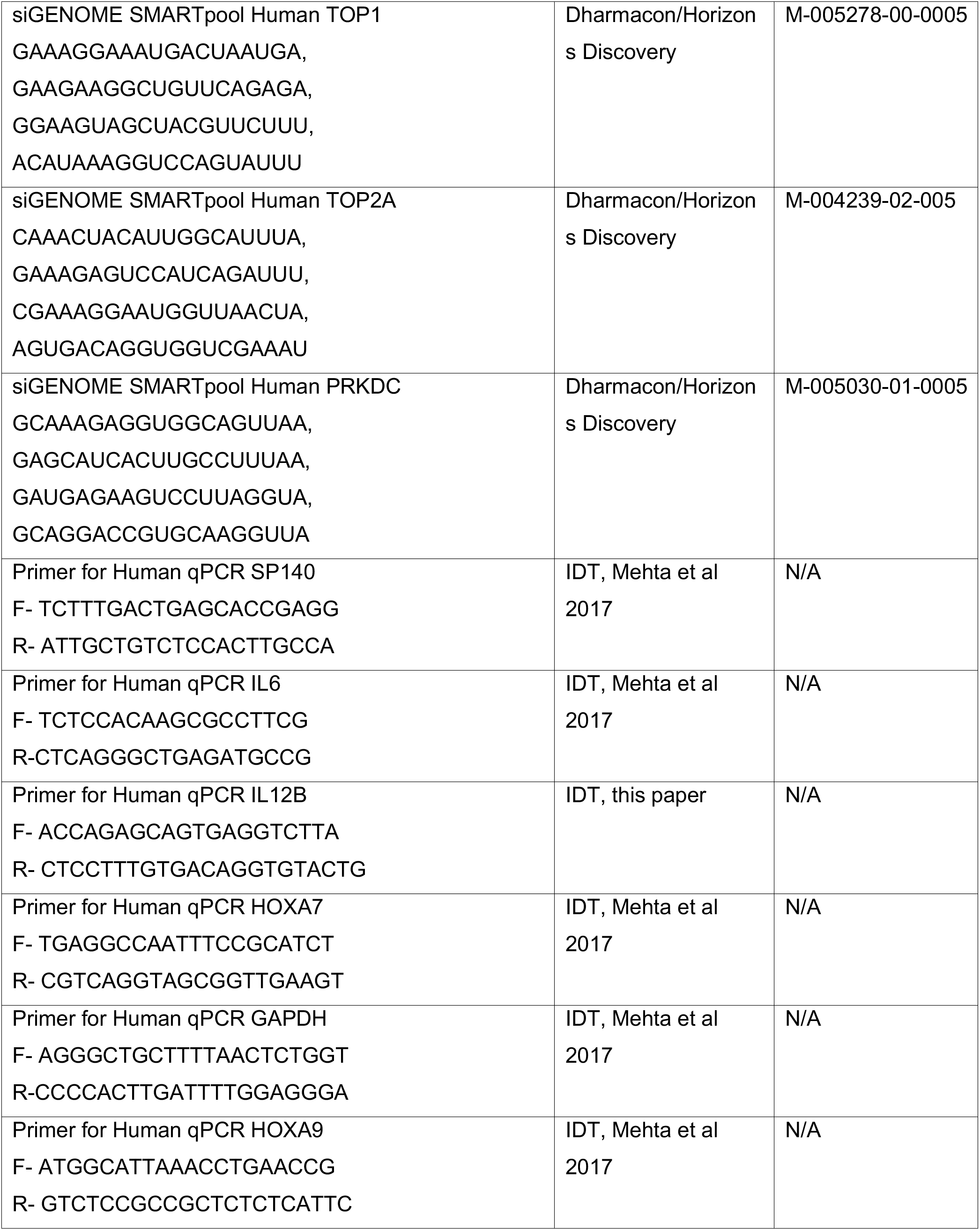
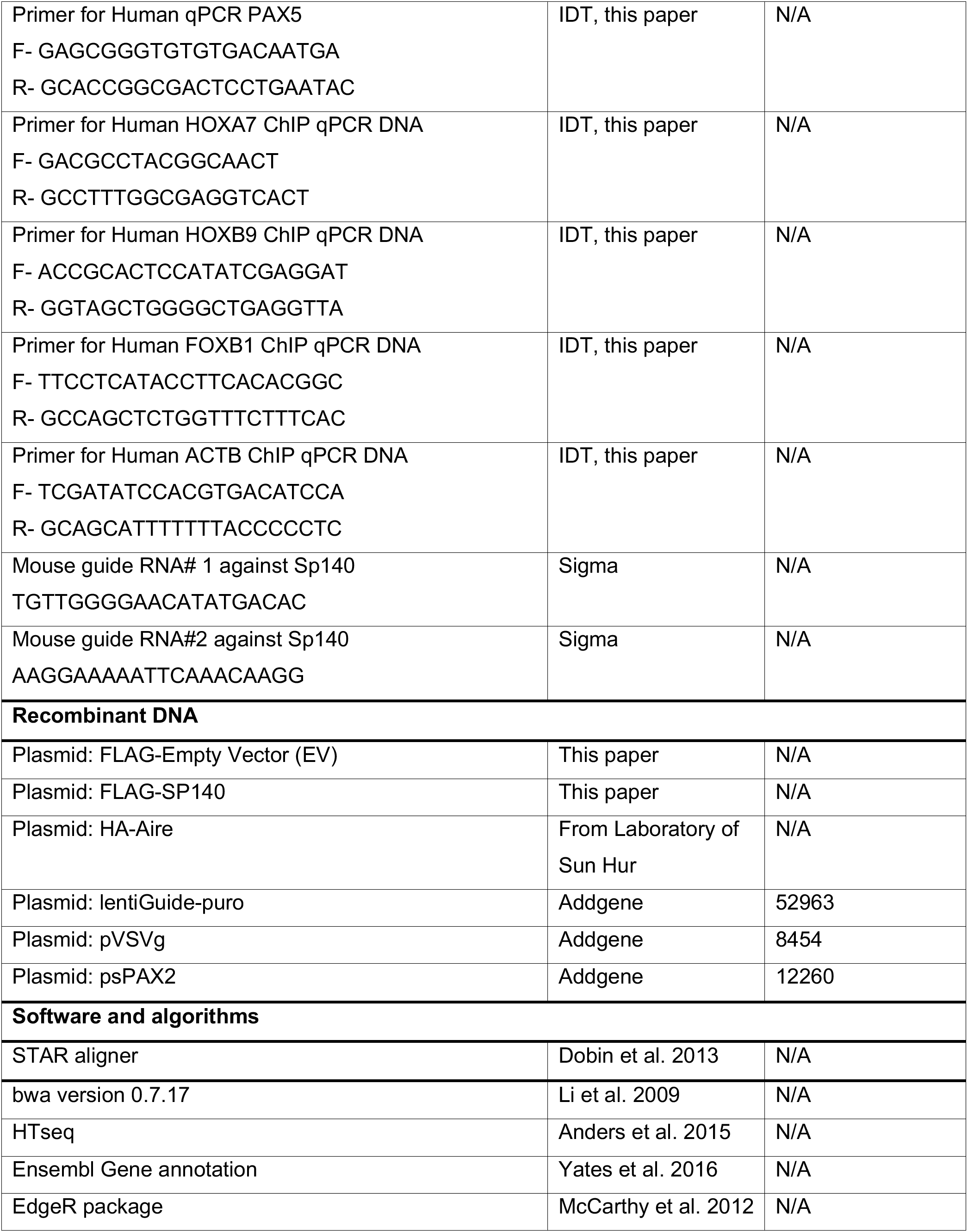
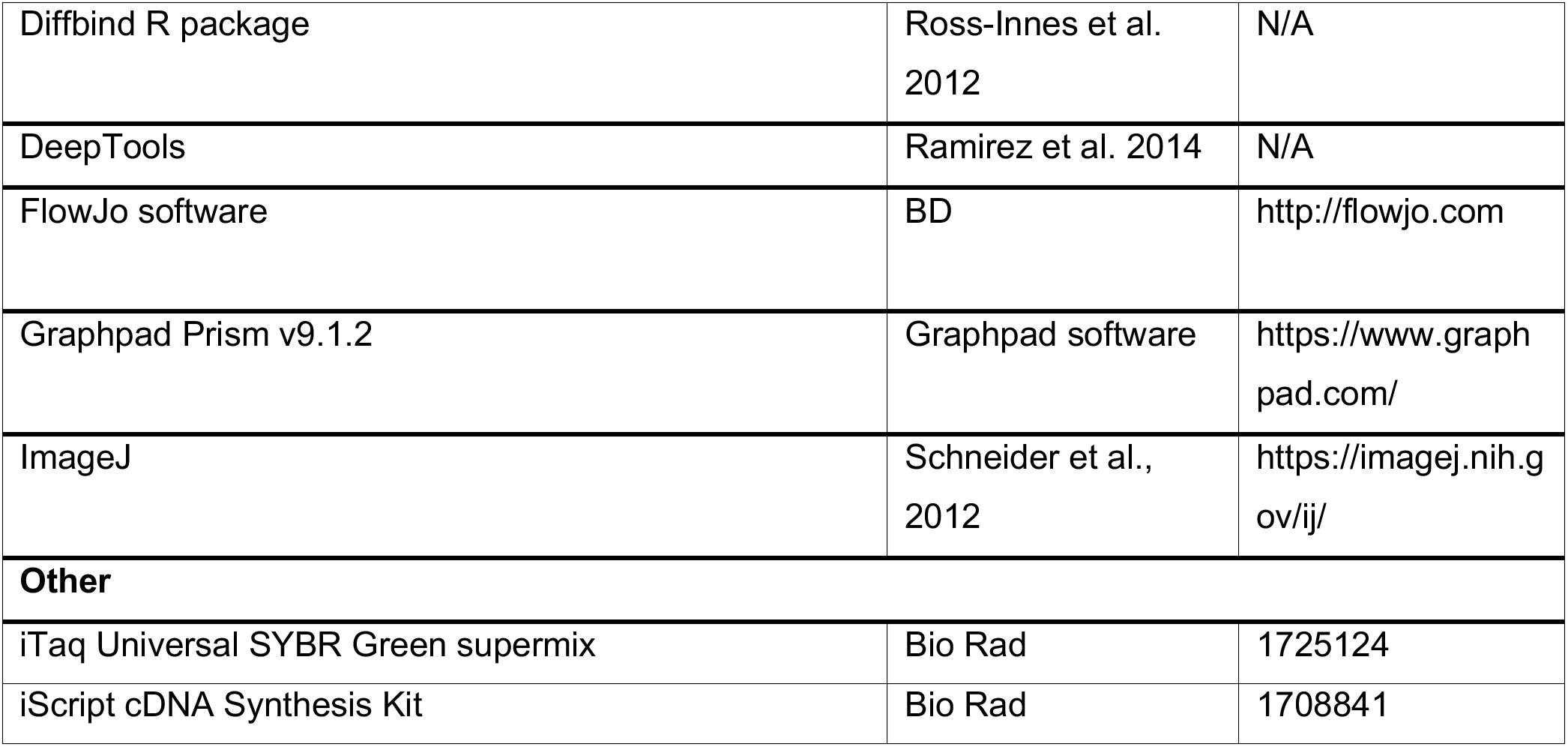
KEY RESOURCES TABLE.

## Notes

### Competing Interest Statement

The authors have declared no competing interest.

### Summary of Updates

This pdf was updated with higher resolution figures

