## Supplementary Material for "Identification of topoisomerase as a precision-medicine target in chromatin reader SP140-driven Crohn’s disease"

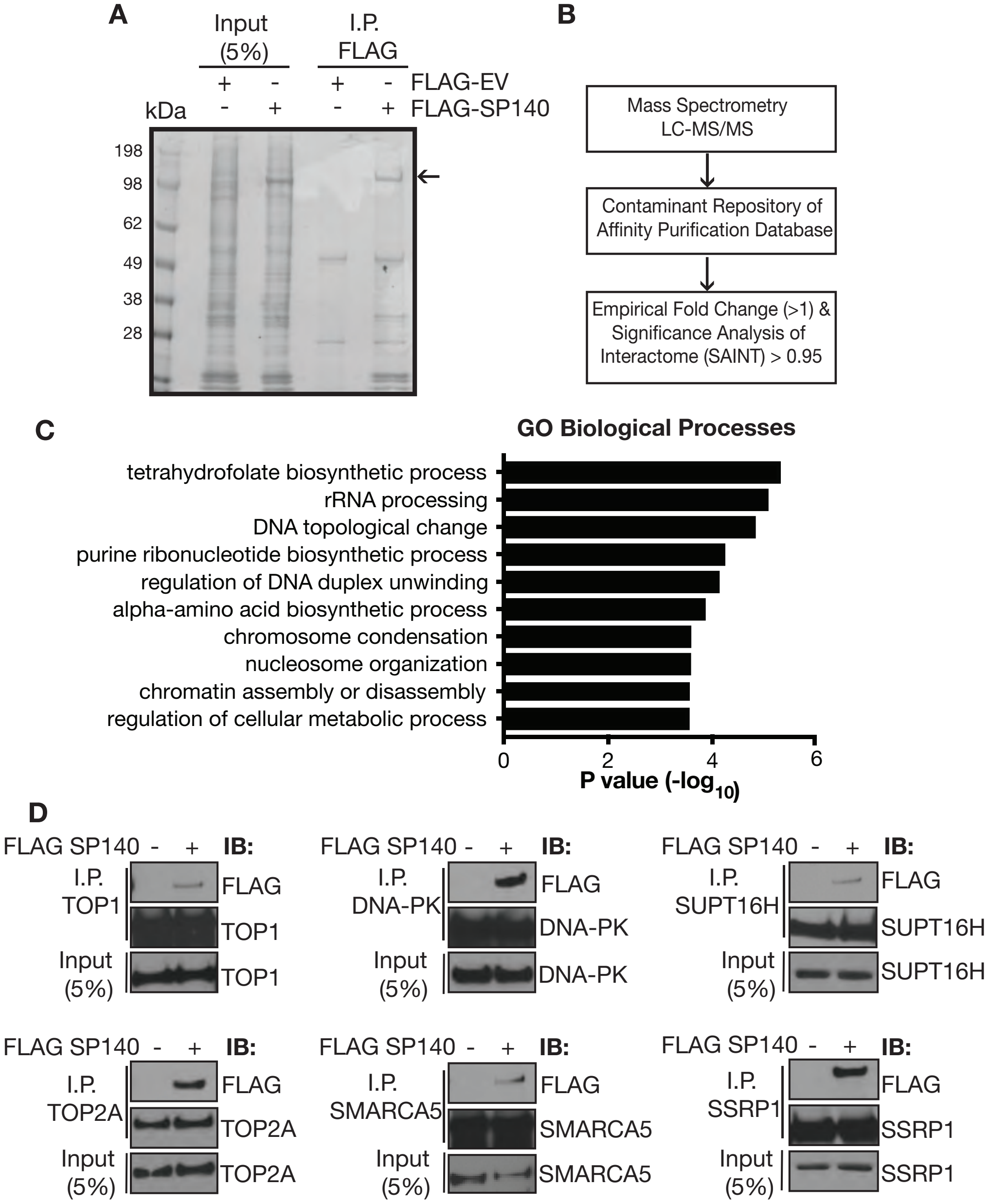


**Figure S1. Characterization of the SP140 interactome. Related to Figure 1. A,** FLAG immunoprecipitation (IP) of HEK293T lysates expressing FLAG-Empty vector (EV) or FLAG-SP140 constructs. Protein complexes were eluted, separated by SDS-PAGE and stained with Silver Stain. Arrow indicates full length SP140 protein. Excised bands were analyzed by Mass Spectrometry (MS). **B,** Flow chart depicting analysis strategy of MS hits**.** Significance analysis of interactome (SAINT) and fold change (FC) over EV FLAG IP were calculated using contaminant repository of affinity matrix (CRAPome) analysis. **C,** Significant SP140 interacting proteins (with SAINT score $\geq$ 0.99 and FC >2) were entered into panther database tools (pantherdb.org) for assessing overrepresented Gene Ontology (GO) Biological Processes. **D,** Reciprocal IPs of nuclear HEK293T lysates overexpressing FLAG EV and FLAG SP140 with indicated endogenous proteins and probed with FLAG antibody.

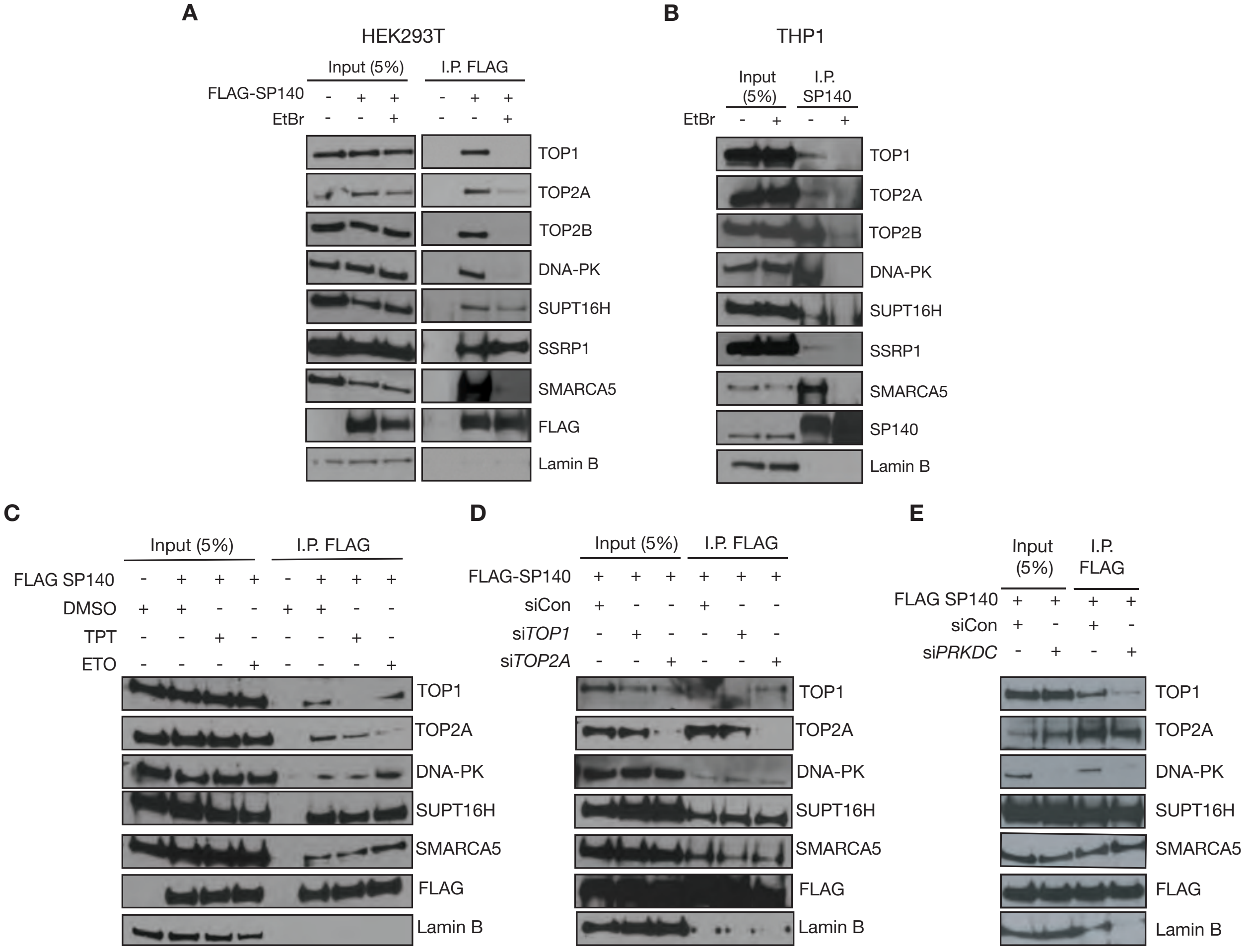


**Figure S2. DNA requirement for SP140 protein interactions. Related to Figure 1. A,** Immunoprecipitation (IP) of FLAG-EV or FLAG-SP140 in HEK293T nuclear lysates in the presence of ethidium bromide (EtBr, 1mg/mL) and immunoblot of indicated endogenous proteins. **B,** IP of endogenous SP140 in THP1 nuclear lysates in the presence of ethidium bromide (EtBr, 1mg/mL) and immunoblot of indicated endogenous proteins. **C,** IP of FLAG EV or FLAG SP140 in HEK293T cells with indicated endogenous proteins in the presence of TOP1 inhibitor topotecan (TPT, 100nM), TOP2 inhibitor etoposide (ETO, 25μM) or DMSO control. **D,** IP of FLAG EV or FLAG SP140 in HEK293T cells and immunoblot of indicated endogenous proteins in the presence of control, *Top1*, or *Top2a* siRNA (100nM). **E,** IP of FLAG EV or FLAG SP140 in HEK293T cells and immunoblot of indicated endogenous proteins in the presence of control or *PRKDC* siRNA (100nM). Data are representative of two independent experiments.

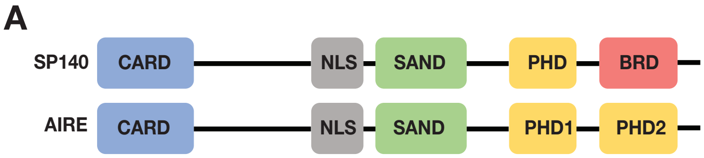


**B**

**Chromatin Remodeling/ DNA unwinding**

| SP140 | AIRE |
|---|---|
| TOP1 | YES |
| TOP2A | YES |
| TOP2B | YES |
| DNA-PK | YES |
| SUPT16H | YES |
| SSRP1 | NO |
| SMARCA5 | NO |

**Transcriptional Elongation**

| AIRE | SP140 |
|---|---|
| PARP1 | NO |
| POLR2B | NO |
| pTEFb | NO |
| CDK7 | NO |
| BRD4 | NO |
| CycT1 | NO |
| HEXIM1 | NO |

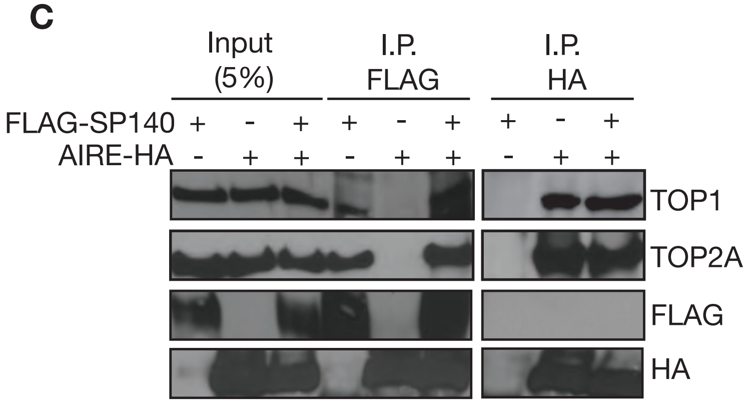


**Figure S3. SP140 and Aire interact with distinct pools of topoisomerases. Related to Figure 1. A,** Visual representation of SP140 and AIRE protein domains. **B,** List of shared and AIRE-exclusive interacting proteins in Chromatin Remodeling/DNA unwinding or Transcriptional Elongation functional categories. **C,** FLAG or HA Immunoprecipitation (IP) of FLAG-SP140, Aire-HA or FLAG-SP140 + Aire-HA expressing HEK293T cells and probed for TOP1 and TOP2A. Data are representative of two independent experiments.

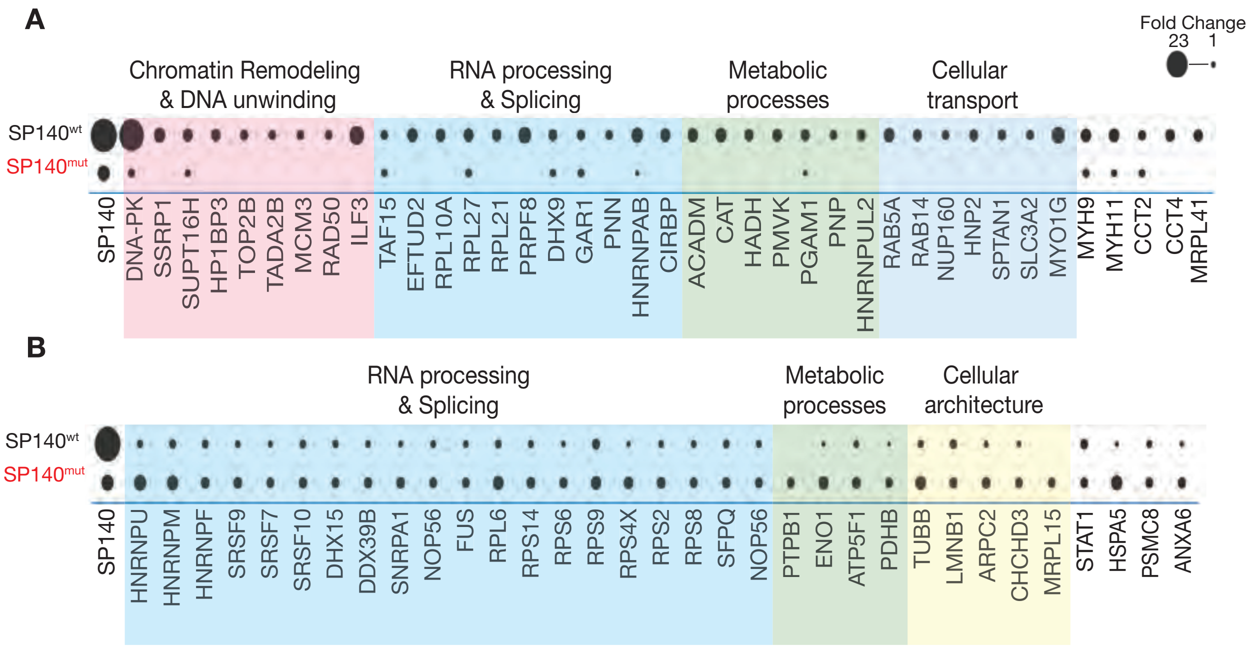


**Figure S4. Rewiring of the SP140 proteome in immune cells bearing CD-risk SP140 mutations. Related to Figure 1. A,** Dot plot displaying empirical fold change of endogenous SP140 IP mass spectrometry (MS) peptide hits over IgG IP control of proteins up-regulated (Fold Change >1.5) or **B,** down-regulated (Fold Change >1.5) in lymphoblastoid cell lines (LBL) from individuals bearing Crohn's disease (CD)-risk SP140 mutations ($SP140^{mut}$) compared to control wild-type SP140 ($SP140^{wt}$). Data are presented as average of two biological replicates of each genotype.

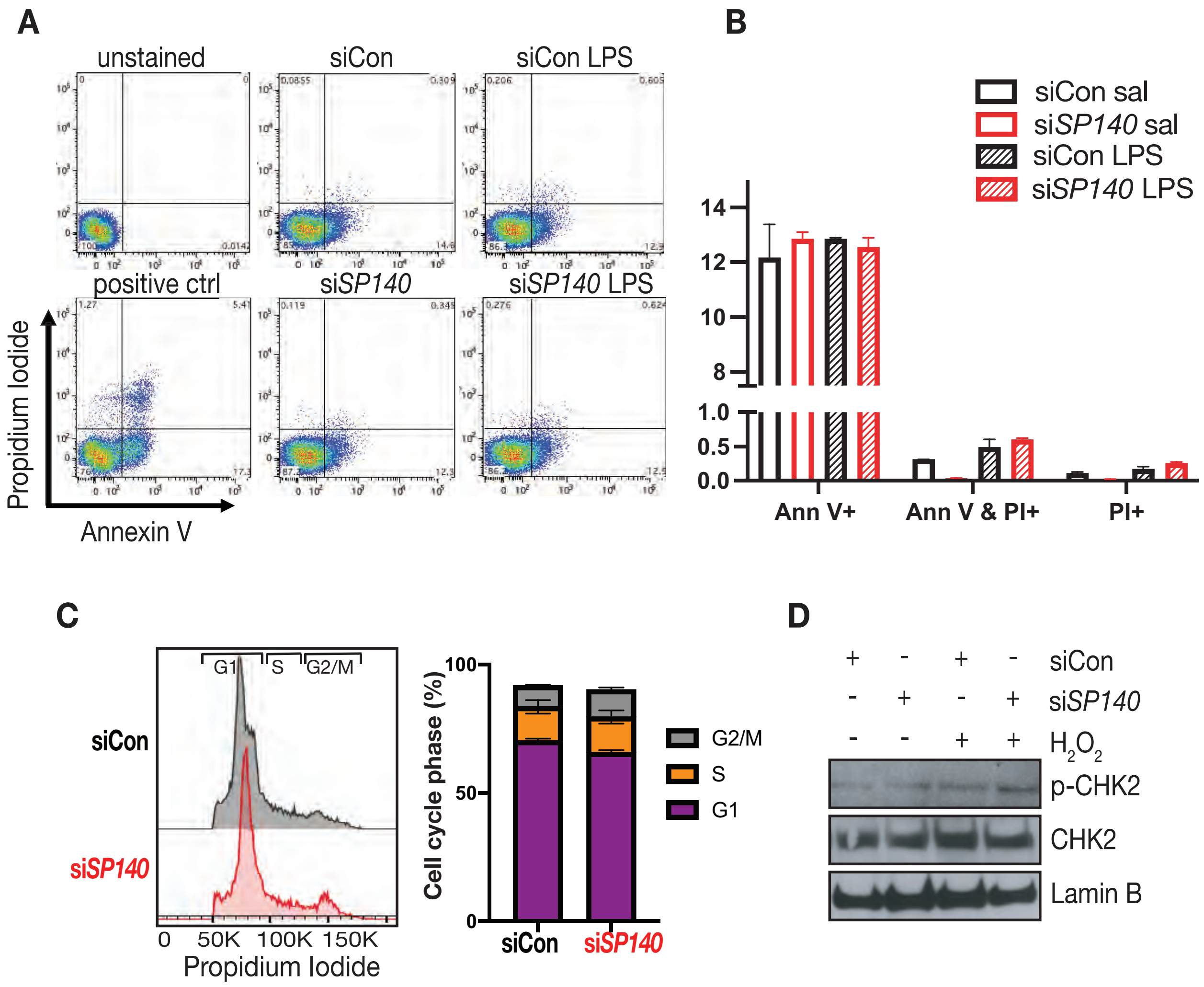


**Figure S5. Cell death or cell cycle are unaffected upon SP140 depletion in human monocytes. Related to Figure 3. A,** Flow Cytometry analysis of Propidium iodide (PI) and Annexin V staining in control or *SP140* siRNA-mediated knockdown THP1 cells with or without LPS treatment (4h, 0.1mg/mL). Positive control are cells treated with $H_2O_2$ (4h, 25μM). **B,** Quantification of percent uptake PI and Annexin V**. C,** Cell cycle quantification by DNA content analysis using PI staining in control and SP140 siRNA knockdown THP1 cells. **D,** Representative Western blot showing phosphorylated (Thr68) and total Checkpoint kinase 2 (CHK2) levels in control and SP140 siRNA knockdown THP-1 cells at baseline or with $H_2O_2$ (4h, 25μM) treatment. Data are mean of three independent experiments. Error bars are s.e.m.

**Extended Data Table 1 | Mass Spectrometry hits of SP140 in HEK293 with Empirical Fold Change (FC) and SAINT probability (SP) values from CRAPome analysis**

| PROTID | GENE | SP140_FC_A | SP140_FC_B | SP140_SP | IP_SP140_REPLICATE 2 | IP_SP140_REPLICATE 1 | IP_CONTROL_REPLICATE 1 |
|---|---|---|---|---|---|---|---|
| Q13342 | SP140 | 35.23 | 33.87 | 1 | 106 | 75 | 0 |
| P11388 | TOP2A | 7.69 | 7.29 | 1 | 22 | 13 | 0 |
| P11586 | MTHFD1 | 6.71 | 5.83 | 1 | 16 | 17 | 0 |
| Q9Y5B9 | SUPT16H | 5.91 | 5.76 | 1 | 15 | 11 | 0 |
| P78527 | PRKDC | 11.83 | 5.57 | 1 | 60 | 62 | 0 |
| P01631 | P01631 | 5.48 | 5.43 | 1 | 9 | 16 | 0 |
| P40939 | HADHA | 5.72 | 5.42 | 1 | 7 | 20 | 0 |
| P49411 | TUFM | 6.38 | 5.37 | 1 | 17 | 14 | 0 |
| P07355 | ANXA2 | 5.36 | 5.36 | 1 | 10 | 14 | 0 |
| P53621 | COPA | 6.37 | 5.27 | 1 | 19 | 11 | 0 |
| P54886 | ALDH18A1 | 6.48 | 5.25 | 1 | 16 | 17 | 0 |
| P04843 | RPN1 | 5.86 | 5 | 1 | 15 | 13 | 0 |
| O75367 | H2AFY | 4.62 | 4.61 | 1 | 8 | 12 | 0 |
| Q9UJS0 | SLC25A13 | 4.62 | 4.61 | 1 | 8 | 12 | 0 |
| P22314 | UBA1 | 4.59 | 4.48 | 1 | 11 | 8 | 0 |
| P49327 | FASN | 8.7 | 4.22 | 1 | 36 | 36 | 0 |
| Q9Y2L1 | DIS3 | 4.17 | 4.14 | 1 | 9 | 8 | 0 |
| Q9UKM9 | RALY | 4.17 | 4.14 | 1 | 9 | 8 | 0 |
| P31939 | ATIC | 3.8 | 3.77 | 1 | 8 | 7 | 0 |
| Q14697 | GANAB | 4.33 | 3.77 | 1 | 12 | 6 | 0 |
| Q15424 | SAFB | 3.63 | 3.5 | 1 | 4 | 11 | 0 |
| Q7KZF4 | SND1 | 4.55 | 3.45 | 1 | 12 | 10 | 0 |
| P84077 | ARF1 | 3.48 | 3.37 | 1 | 8 | 5 | 0 |
| O60264 | SMARCA5 | 3.34 | 3.33 | 1 | 5 | 8 | 0 |
| P31942 | HNRNPH3 | 3.34 | 3.33 | 1 | 5 | 8 | 0 |
| Q9H9B4 | SFXN1 | 3.22 | 3.21 | 1 | 6 | 6 | 0 |
| Q13595 | TRA2A | 3.27 | 3.21 | 1 | 7 | 5 | 0 |
| Q12931 | TRAP1 | 5.18 | 3.15 | 1 | 18 | 16 | 0 |
| O00567 | NOP56 | 4.51 | 3.13 | 1 | 11 | 13 | 0 |
| Q9H583 | HEATR1 | 3.06 | 3.03 | 1 | 6 | 5 | 0 |
| Q5BKZ1 | ZNF326 | 3.02 | 3.02 | 1 | 5 | 6 | 0 |
| P45880 | VDAC2 | 3.09 | 2.97 | 1 | 3 | 9 | 0 |
| P61221 | ABCE1 | 2.97 | 2.96 | 1 | 4 | 7 | 0 |
| O14980 | XPO1 | 3.2 | 2.9 | 1 | 4 | 9 | 0 |
| O76094 | SRP72 | 2.93 | 2.85 | 1 | 3 | 8 | 0 |
| P15311 | EZR | 2.85 | 2.85 | 1 | 5 | 5 | 0 |
| P49736 | MCM2 | 2.81 | 2.81 | 1 | 4 | 6 | 0 |
| Q02880 | TOP2B | 2.81 | 2.81 | 1 | 4 | 6 | 0 |
| P63244 | RACK1 | 3.24 | 2.8 | 1 | 7 | 6 | 0 |
| Q9BYG3 | NIFK | 2.94 | 2.78 | 1 | 7 | 3 | 0 |
| P46087 | NOP2 | 4.22 | 2.77 | 1 | 11 | 12 | 0 |
| P00403 | MT-CO2 | 2.69 | 2.66 | 1 | 5 | 4 | 0 |
| P06737 | PYGL | 2.69 | 2.66 | 1 | 5 | 4 | 0 |
| Q9Y678 | COPG1 | 2.65 | 2.65 | 1 | 4 | 5 | 0 |
| P05090 | APOD | 2.65 | 2.65 | 1 | 4 | 5 | 0 |
| P05023 | ATP1A1 | 4.04 | 2.61 | 1 | 11 | 11 | 0 |
| P51149 | RAB7A | 2.6 | 2.58 | 1 | 3 | 6 | 0 |
| O00154 | ACOT7 | 2.48 | 2.48 | 1 | 4 | 4 | 0 |
| A5YKK6 | CNOT1 | 2.44 | 2.43 | 1 | 3 | 5 | 0 |
| Q9BVP2 | GNL3 | 3 | 2.34 | 1 | 5 | 8 | 0 |
| Q9UKV3 | ACIN1 | 5.29 | 2.29 | 1 | 28 | 17 | 1 |
| P33993 | MCM7 | 3.68 | 2.11 | 1 | 5 | 17 | 1 |
| Q7L2E3 | DHX30 | 3.99 | 1.93 | 1 | 17 | 11 | 0 |
| P51659 | HSD17B4 | 3.87 | 3.16 | 0.99 | 11 | 5 | 0 |
| Q86VP6 | CAND1 | 3.26 | 3.1 | 0.99 | 3 | 10 | 0 |
| Q8TEM1 | NUP210 | 2.77 | 2.72 | 0.99 | 3 | 7 | 0 |
| P11387 | TOP1 | 3.6 | 2.68 | 0.99 | 8 | 9 | 0 |
| Q92598 | HSPH1 | 2.69 | 2.66 | 0.99 | 5 | 4 | 0 |
| Q9NSD9 | FARSB | 2.69 | 2.66 | 0.99 | 5 | 4 | 0 |
| Q5SSJ5 | HP1BP3 | 2.65 | 2.65 | 0.99 | 4 | 5 | 0 |
| Q96RP9 | GFM1 | 2.73 | 2.63 | 0.99 | 6 | 3 | 0 |

| | | | | | | | |
|---|---|---|---|---|---|---|---|
| P22102 | GART | 2.89 | 2.5 | 0.99 | 6 | 5 | 0 |
| Q15269 | PWP2 | 2.48 | 2.48 | 0.99 | 4 | 4 | 0 |
| Q96GQ7 | DDX27 | 2.53 | 2.47 | 0.99 | 5 | 3 | 0 |
| P21796 | VDAC1 | 2.53 | 2.47 | 0.99 | 5 | 3 | 0 |
| Q13838 | DDX39B | 2.74 | 2.34 | 0.99 | 6 | 4 | 0 |
| Q96HS1 | PGAM5 | 2.83 | 2.34 | 0.99 | 6 | 5 | 0 |
| P08243 | ASNS | 2.32 | 2.29 | 0.99 | 4 | 3 | 0 |
| Q6P2Q9 | PRPF8 | 4.22 | 2.28 | 0.99 | 31 | 38 | 0 |
| Q2NL82 | TSR1 | 2.11 | 2.11 | 0.99 | 3 | 3 | 0 |
| P26038 | MSN | 2.11 | 2.11 | 0.99 | 3 | 3 | 0 |
| O75746 | SLC25A12 | 2.11 | 2.11 | 0.99 | 3 | 3 | 0 |
| P14618 | PKM | 4.08 | 2.1 | 0.99 | 15 | 13 | 0 |
| O15371 | EIF3D | 1.83 | 1 | 0.99 | 4 | 5 | 0 |
| P68400 | CSNK2A1 | 2.73 | 2.63 | 0.98 | 6 | 3 | 0 |
| Q8IY81 | FTSJ3 | 2.61 | 2.39 | 0.98 | 5 | 4 | 0 |
| P27695 | APEX1 | 2.23 | 2.2 | 0.98 | 2 | 5 | 0 |
| O14983 | ATP2A1 | 2.54 | 2.2 | 0.98 | 5 | 4 | 0 |
| P40937 | RFC5 | 2.23 | 2.2 | 0.98 | 2 | 5 | 0 |
| P18754 | RCC1 | 2.23 | 2.2 | 0.98 | 2 | 5 | 0 |
| P31930 | UQCRC1 | 2.11 | 2.11 | 0.98 | 3 | 3 | 0 |
| Q8WVM0 | TFB1M | 2.11 | 2.11 | 0.98 | 3 | 3 | 0 |
| Q15517 | CDSN | 2.11 | 2.11 | 0.98 | 3 | 3 | 0 |
| P28370 | SMARCA1 | 2.07 | 2.05 | 0.98 | 2 | 4 | 0 |
| Q7Z7K6 | CENPV | 2.07 | 2.05 | 0.98 | 2 | 4 | 0 |
| Q08J23 | NSUN2 | 2.07 | 2.05 | 0.98 | 2 | 4 | 0 |
| Q92620 | DHX38 | 1.91 | 1.9 | 0.98 | 2 | 3 | 0 |
| Q14739 | LBR | 1.91 | 1.9 | 0.98 | 2 | 3 | 0 |
| P53985 | SLC16A1 | 2.15 | 1.77 | 0.98 | 4 | 3 | 0 |
| P40926 | MDH2 | 2.4 | 2.33 | 0.97 | 2 | 6 | 0 |
| Q14690 | PDCD11 | 2.4 | 2.33 | 0.97 | 2 | 6 | 0 |
| O75439 | PMPCB | 2.36 | 2.25 | 0.97 | 5 | 2 | 0 |
| Q14151 | SAFB2 | 2.16 | 2.1 | 0.97 | 4 | 2 | 0 |
| P53004 | BLVRA | 2.16 | 2.1 | 0.97 | 4 | 2 | 0 |
| Q16836 | HADH | 2.07 | 2.05 | 0.97 | 2 | 4 | 0 |
| O60832 | DKC1 | 2.21 | 2.04 | 0.97 | 3 | 4 | 0 |
| P61026 | RAB10 | 1.95 | 1.93 | 0.97 | 3 | 2 | 0 |
| Q99798 | ACO2 | 1.95 | 1.93 | 0.97 | 3 | 2 | 0 |
| Q9H0S4 | DDX47 | 1.91 | 1.9 | 0.97 | 2 | 3 | 0 |
| Q9BSJ8 | ESYT1 | 1.91 | 1.9 | 0.97 | 2 | 3 | 0 |
| Q9BXJ9 | NAA15 | 1.91 | 1.9 | 0.97 | 2 | 3 | 0 |
| Q99873 | PRMT1 | 2.58 | 1.9 | 0.97 | 3 | 8 | 0 |
| Q8WXF1 | PSPC1 | 2.1 | 1.66 | 0.97 | 4 | 3 | 0 |
| Q9UJZ1 | STOML2 | 3.38 | 3 | 0.96 | 2 | 12 | 0 |
| Q9BQ39 | DDX50 | 3.04 | 2.38 | 0.96 | 8 | 4 | 0 |
| Q8NI36 | WDR36 | 2.25 | 2.06 | 0.96 | 4 | 3 | 0 |
| Q9H0D6 | XRN2 | 2.25 | 2.06 | 0.96 | 4 | 3 | 0 |
| Q7Z2W4 | ZC3HAV1 | 1.95 | 1.93 | 0.96 | 3 | 2 | 0 |
| P55039 | DRG2 | 1.95 | 1.93 | 0.96 | 3 | 2 | 0 |
| P08754 | GNAI3 | 1.95 | 1.93 | 0.96 | 3 | 2 | 0 |
| P63165 | SUMO1 | 1.91 | 1.9 | 0.96 | 2 | 3 | 0 |
| Q13330 | MTA1 | 1.74 | 1.74 | 0.96 | 2 | 2 | 0 |
| O00178 | GTPBP1 | 1.74 | 1.74 | 0.96 | 2 | 2 | 0 |
| P13804 | ETFA | 1.74 | 1.74 | 0.96 | 2 | 2 | 0 |
| Q8WXE9 | STON2 | 1.74 | 1.74 | 0.96 | 2 | 2 | 0 |
| P49792 | RANBP2 | 1.93 | 0.99 | 0.96 | 4 | 7 | 0 |
| Q96PK6 | RBM14 | 1.53 | 0.66 | 0.96 | 6 | 4 | 0 |
| O75694 | NUP155 | 2.16 | 2.1 | 0.95 | 4 | 2 | 0 |
| P11216 | PYGB | 2.07 | 2.05 | 0.95 | 2 | 4 | 0 |
| P02786 | TFRC | 1.91 | 1.9 | 0.95 | 2 | 3 | 0 |
| Q9BV20 | MRI1 | 1.74 | 1.74 | 0.95 | 2 | 2 | 0 |
| Q15436 | SEC23A | 1.74 | 1.74 | 0.95 | 2 | 2 | 0 |
| Q9NR31 | SAR1A | 1.74 | 1.74 | 0.95 | 2 | 2 | 0 |
| P62820 | RAB1A | 2.1 | 1.53 | 0.95 | 3 | 5 | 0 |
| P82650 | MRPS22 | 1.84 | 1.4 | 0.95 | 2 | 4 | 0 |
| Q6UB35 | MTHFD1L | 2.24 | 1.33 | 0.95 | 5 | 6 | 0 |
| Q9BZE4 | GTPBP4 | 2.65 | 2.35 | 0.94 | 6 | 3 | 0 |
| Q8N1F7 | NUP93 | 2.38 | 2 | 0.94 | 5 | 3 | 0 |
| O43491 | EPB41L2 | 1.95 | 1.93 | 0.94 | 3 | 2 | 0 |

| | | | | | | | |
|---|---|---|---|---|---|---|---|
| P55786 | NPEPPS | 1.74 | 1.74 | 0.94 | 2 | 2 | 0 |
| Q92643 | PIGK | 1.74 | 1.74 | 0.94 | 2 | 2 | 0 |
| Q8IWA4 | MFN1 | 1.74 | 1.74 | 0.94 | 2 | 2 | 0 |
| Q9Y4L1 | HYOU1 | 1.74 | 1.74 | 0.94 | 2 | 2 | 0 |
| Q8TCJ2 | STT3B | 1.74 | 1.74 | 0.94 | 2 | 2 | 0 |
| O14617 | AP3D1 | 2.18 | 1.86 | 0.93 | 4 | 3 | 0 |
| Q15031 | LARS2 | 1.74 | 1.74 | 0.93 | 2 | 2 | 0 |
| Q9NVP1 | DDX18 | 2.03 | 1.35 | 0.93 | 4 | 4 | 0 |
| Q01780 | EXOSC10 | 2.64 | 2.3 | 0.92 | 2 | 8 | 0 |
| P57740 | NUP107 | 1.74 | 1.74 | 0.92 | 2 | 2 | 0 |
| P55060 | CSE1L | 1.85 | 1.71 | 0.92 | 2 | 3 | 0 |
| O43684 | BUB3 | 2.43 | 1.98 | 0.91 | 6 | 2 | 0 |
| P15121 | AKR1B1 | 1.7 | 1.67 | 0.91 | 1 | 3 | 0 |
| O60783 | MRPS14 | 1.53 | 1.53 | 0.91 | 1 | 2 | 0 |
| Q9P2E9 | RRBP1 | 1.99 | 1.32 | 0.91 | 3 | 5 | 0 |
| Q14152 | EIF3A | 6.22 | 3.52 | 0.9 | 26 | 6 | 0 |
| P14625 | HSP90B1 | 3.57 | 2.02 | 0.9 | 9 | 13 | 0 |
| Q96N66 | MBOAT7 | 1.7 | 1.67 | 0.9 | 1 | 3 | 0 |
| P23284 | PPIB | 1.7 | 1.67 | 0.9 | 1 | 3 | 0 |
| P00367 | GLUD1 | 1.69 | 1.56 | 0.9 | 2 | 2 | 0 |
| P05198 | EIF2S1 | 2.49 | 1.46 | 0.9 | 5 | 9 | 0 |
| Q9BXP5 | SRRT | 1.74 | 1.08 | 0.9 | 2 | 5 | 0 |
| P13639 | EEF2 | 3.79 | 1.96 | 0.89 | 10 | 20 | 0 |
| O00487 | PSMD14 | 2.27 | 1.84 | 0.89 | 4 | 4 | 0 |
| P62834 | RAP1A | 1.7 | 1.67 | 0.89 | 1 | 3 | 0 |
| P17612 | PRKACA | 1.7 | 1.67 | 0.89 | 1 | 3 | 0 |
| Q12849 | GRSF1 | 1.53 | 1.53 | 0.89 | 1 | 2 | 0 |
| Q8NE71 | ABCF1 | 2.55 | 2.06 | 0.88 | 2 | 8 | 0 |
| P07737 | PFN1 | 2.18 | 1.68 | 0.88 | 3 | 5 | 0 |
| P57088 | TMEM33 | 1.58 | 1.56 | 0.88 | 2 | 1 | 0 |
| P36776 | LONP1 | 1.79 | 1.54 | 0.88 | 2 | 3 | 0 |
| O96008 | TOMM40 | 2.02 | 1.93 | 0.87 | 1 | 5 | 0 |
| Q9Y276 | BCS1L | 2.02 | 1.93 | 0.87 | 1 | 5 | 0 |
| P12273 | PIP | 1.99 | 1.88 | 0.87 | 4 | 1 | 0 |
| P53618 | COPB1 | 2.22 | 1.83 | 0.87 | 5 | 2 | 0 |
| P22087 | FBL | 2.59 | 1.7 | 0.87 | 6 | 6 | 0 |
| P11172 | UMPS | 1.53 | 1.53 | 0.87 | 1 | 2 | 0 |
| Q15717 | ELAVL1 | 2.72 | 1.53 | 0.87 | 8 | 7 | 0 |
| Q8N752 | CSNK1A1L | 1.79 | 1.73 | 0.86 | 3 | 1 | 0 |
| P17812 | CTPS1 | 1.79 | 1.73 | 0.86 | 3 | 1 | 0 |
| Q14694 | USP10 | 1.7 | 1.67 | 0.86 | 1 | 3 | 0 |
| P07741 | APRT | 1.58 | 1.56 | 0.86 | 2 | 1 | 0 |
| P61160 | ACTR2 | 1.53 | 1.53 | 0.86 | 1 | 2 | 0 |
| O00116 | AGPS | 1.53 | 1.53 | 0.86 | 1 | 2 | 0 |
| Q6PI48 | DARS2 | 1.53 | 1.53 | 0.86 | 1 | 2 | 0 |
| Q53GS9 | USP39 | 1.75 | 1.43 | 0.86 | 2 | 3 | 0 |
| Q9H0A0 | NAT10 | 3.7 | 3.19 | 0.85 | 2 | 14 | 0 |
| P82933 | MRPS9 | 1.79 | 1.73 | 0.85 | 3 | 1 | 0 |
| P49915 | GMPS | 1.79 | 1.73 | 0.85 | 3 | 1 | 0 |
| Q8NI27 | THOC2 | 1.84 | 1.59 | 0.85 | 3 | 2 | 0 |
| P82921 | MRPS21 | 1.58 | 1.56 | 0.85 | 2 | 1 | 0 |
| Q96EY1 | DNAJA3 | 1.79 | 1.54 | 0.85 | 2 | 3 | 0 |
| P04004 | VTN | 1.53 | 1.53 | 0.85 | 1 | 2 | 0 |
| Q14166 | TTLL12 | 1.53 | 1.53 | 0.85 | 1 | 2 | 0 |
| Q14683 | SMC1A | 1.75 | 1.43 | 0.85 | 2 | 3 | 0 |
| Q9HCE1 | MOV10 | 1.61 | 1.13 | 0.85 | 2 | 3 | 0 |
| O15160 | POLR1C | 1.99 | 1.88 | 0.84 | 4 | 1 | 0 |
| P52209 | PGD | 1.79 | 1.73 | 0.84 | 3 | 1 | 0 |
| P51114 | FXR1 | 1.79 | 1.73 | 0.84 | 3 | 1 | 0 |
| Q96GD4 | AURKB | 1.58 | 1.56 | 0.84 | 2 | 1 | 0 |
| A1L0T0 | ILVBL | 1.53 | 1.53 | 0.84 | 1 | 2 | 0 |
| Q15365 | PCBP1 | 2.15 | 1.25 | 0.84 | 6 | 4 | 0 |
| P33991 | MCM4 | 1.35 | 0.69 | 0.84 | 2 | 4 | 0 |
| P41250 | GARS | 1.79 | 1.73 | 0.83 | 3 | 1 | 0 |
| O15226 | NKRF | 1.79 | 1.73 | 0.83 | 3 | 1 | 0 |
| Q9BXW7 | CECR5 | 1.79 | 1.73 | 0.83 | 3 | 1 | 0 |
| Q8WVM8 | SCFD1 | 1.79 | 1.73 | 0.83 | 3 | 1 | 0 |
| P02748 | C9 | 1.58 | 1.56 | 0.83 | 2 | 1 | 0 |

| | | | | | | | |
|---|---|---|---|---|---|---|---|
| P51116 | FXR2 | 1.6 | 1.38 | 0.83 | 1 | 3 | 0 |
| Q9NVI7 | ATAD3A | 1.8 | 1.29 | 0.82 | 3 | 3 | 0 |
| Q9NYK5 | MRPL39 | 2.52 | 2.26 | 0.81 | 1 | 8 | 0 |
| P34932 | HSPA4 | 2.35 | 2.16 | 0.81 | 1 | 7 | 0 |
| P11217 | PYGM | 2.02 | 1.93 | 0.81 | 1 | 5 | 0 |
| O43837 | IDH3B | 1.99 | 1.88 | 0.81 | 4 | 1 | 0 |
| P57678 | GEMIN4 | 1.8 | 1.62 | 0.81 | 1 | 4 | 0 |
| Q9NR30 | DDX21 | 3.86 | 1.51 | 0.81 | 23 | 19 | 0 |
| Q9BXS5 | AP1M1 | 1.45 | 1.26 | 0.81 | 1 | 2 | 0 |
| P62995 | TRA2B | 2.51 | 1.24 | 0.81 | 10 | 4 | 0 |
| Q96EY7 | PTCD3 | 1.58 | 1.56 | 0.8 | 2 | 1 | 0 |
| Q9UHI6 | DDX20 | 1.53 | 1.53 | 0.8 | 1 | 2 | 0 |
| Q1KMD3 | HNRNPUL2 | 1.74 | 1.45 | 0.8 | 1 | 4 | 0 |
| O95347 | SMC2 | 1.75 | 1.43 | 0.8 | 2 | 3 | 0 |
| B7ZW38 | HNRNPCL3 | 2.39 | 1.02 | 0.8 | 6 | 15 | 0 |
| Q8IX18 | DHX40 | 1.58 | 1.56 | 0.79 | 2 | 1 | 0 |
| P04844 | RPN2 | 1.68 | 1.29 | 0.79 | 2 | 3 | 0 |
| Q9P035 | HACD3 | 1.49 | 1.28 | 0.79 | 2 | 1 | 0 |
| Q14980 | NUMA1 | 1.74 | 1.74 | 0.78 | 2 | 2 | 0 |
| Q08945 | SSRP1 | 2.05 | 1.64 | 0.78 | 1 | 6 | 0 |
| P49588 | AARS | 1.58 | 1.56 | 0.77 | 2 | 1 | 0 |
| P00558 | PGK1 | 1.21 | 0.73 | 0.77 | 2 | 1 | 1 |
| Q96T37 | RBM15 | 1.58 | 1.56 | 0.76 | 2 | 1 | 0 |
| P62136 | PPP1CA | 1.75 | 1.47 | 0.76 | 1 | 4 | 0 |
| P04406 | GAPDH | 2.55 | 1.32 | 0.76 | 6 | 10 | 0 |
| Q15366 | PCBP2 | 1.77 | 1.23 | 0.76 | 3 | 3 | 0 |
| P11177 | PDHB | 1.96 | 0.95 | 0.76 | 6 | 6 | 0 |
| O94973 | AP2A2 | 1.58 | 1.56 | 0.75 | 2 | 1 | 0 |
| P62873 | GNB1 | 1.6 | 1.31 | 0.75 | 2 | 2 | 0 |
| Q5JWF2 | GNAS | 1.58 | 1.56 | 0.74 | 2 | 1 | 0 |
| Q14566 | MCM6 | 1.67 | 1.27 | 0.74 | 1 | 4 | 0 |
| P06733 | ENO1 | 2.81 | 0.91 | 0.72 | 7 | 32 | 4 |
| Q9P2J5 | LARS | 3.55 | 1.57 | 0.71 | 18 | 19 | 0 |
| P55884 | EIF3B | 2.05 | 1.4 | 0.71 | 4 | 4 | 0 |
| P07305 | H1F0 | 1.72 | 0.96 | 0.71 | 3 | 5 | 0 |
| O76021 | RSL1D1 | 2.2 | 0.95 | 0.71 | 6 | 12 | 0 |
| Q86UE4 | MTDH | 1.92 | 1.12 | 0.69 | 6 | 1 | 0 |
| P52292 | KPNA2 | 1.34 | 0.83 | 0.69 | 2 | 2 | 0 |
| P60842 | EIF4A1 | 3.79 | 1.88 | 0.68 | 4 | 21 | 0 |
| P12277 | CKB | 2 | 1.52 | 0.68 | 1 | 6 | 0 |
| Q9Y295 | DRG1 | 2 | 1.35 | 0.68 | 2 | 6 | 0 |
| Q96KR1 | ZFR | 1.48 | 1.25 | 0.68 | 2 | 1 | 0 |
| Q9NTJ3 | SMC4 | 1.53 | 1.4 | 0.67 | 2 | 1 | 0 |
| Q8WTT2 | NOC3L | 1.48 | 1.26 | 0.67 | 2 | 1 | 0 |
| P39656 | DDOST | 1.84 | 0.97 | 0.67 | 1 | 8 | 1 |
| P38159 | RBMX | 2.93 | 1.37 | 0.66 | 12 | 9 | 0 |
| Q14684 | RRP1B | 1.57 | 0.83 | 0.66 | 1 | 6 | 1 |
| Q9UQE7 | SMC3 | 1.49 | 1.28 | 0.65 | 2 | 1 | 0 |
| P26640 | VARS | 2.31 | 1.24 | 0.65 | 3 | 10 | 0 |
| Q92499 | DDX1 | 1.41 | 0.66 | 0.65 | 5 | 1 | 0 |
| Q7L576 | CYFIP1 | 1.58 | 1.56 | 0.64 | 2 | 1 | 0 |
| P20020 | ATP2B1 | 1.58 | 1.56 | 0.64 | 2 | 1 | 0 |
| O75643 | SNRNP200 | 3.35 | 1.59 | 0.62 | 22 | 21 | 0 |
| Q99460 | PSMD1 | 1.63 | 1.28 | 0.62 | 3 | 1 | 0 |
| P17980 | PSMC3 | 1.83 | 1.26 | 0.62 | 4 | 2 | 0 |
| P07195 | LDHB | 1.84 | 1.22 | 0.62 | 3 | 4 | 0 |
| P43243 | MATR3 | 2.46 | 1.19 | 0.62 | 6 | 11 | 0 |
| O15523 | DDX3Y | 2.26 | 1.08 | 0.62 | 5 | 17 | 0 |
| P04075 | ALDOA | 1.72 | 1.17 | 0.61 | 2 | 4 | 0 |
| Q92621 | NUP205 | 3.33 | 2.73 | 0.6 | 1 | 13 | 0 |
| O75165 | DNAJC13 | 1.7 | 1.67 | 0.59 | 1 | 3 | 0 |
| Q9H307 | PNN | 1.07 | 0.29 | 0.58 | 21 | 8 | 17 |
| P49368 | CCT3 | 2.87 | 1.41 | 0.57 | 4 | 15 | 0 |
| Q9NX58 | LYAR | 1.55 | 1.24 | 0.57 | 1 | 3 | 0 |
| Q14676 | MDC1 | 1.99 | 1.88 | 0.56 | 4 | 1 | 0 |
| Q9UQ80 | PA2G4 | 0.99 | 0.32 | 0.56 | 6 | 12 | 10 |
| P51532 | SMARCA4 | 1.58 | 1.56 | 0.55 | 2 | 1 | 0 |
| P27708 | CAD | 2.24 | 1.14 | 0.55 | 1 | 11 | 1 |

| | | | | | | | |
|---|---|---|---|---|---|---|---|
| P35232 | PHB | 3.3 | 1.19 | 0.54 | 2 | 25 | 1 |
| O14654 | IRS4 | 1.14 | 0.55 | 0.54 | 1 | 4 | 0 |
| P51610 | HCFC1 | 1.58 | 1.56 | 0.53 | 2 | 1 | 0 |
| Q99613 | EIF3C | 1.58 | 0.67 | 0.53 | 7 | 10 | 0 |
| P29803 | PDHA2 | 1.25 | 0.79 | 0.52 | 2 | 1 | 0 |
| Q14974 | KPNB1 | 1.35 | 0.65 | 0.52 | 3 | 4 | 0 |
| Q92616 | GCN1 | 2.71 | 1.65 | 0.5 | 0 | 12 | 0 |
| P00338 | LDHA | 1.92 | 1.54 | 0.5 | 0 | 6 | 0 |
| P55084 | HADHB | 2.03 | 1.5 | 0.5 | 0 | 7 | 0 |
| Q9UBX3 | SLC25A10 | 2.14 | 1.81 | 0.49 | 0 | 7 | 0 |
| P19623 | SRM | 1.65 | 1.52 | 0.49 | 0 | 4 | 0 |
| Q53GQ0 | HSD17B12 | 1.65 | 1.52 | 0.49 | 0 | 4 | 0 |
| Q00325 | SLC25A3 | 1.86 | 1.4 | 0.49 | 0 | 6 | 0 |
| P46776 | RPL27A | 1.64 | 1.15 | 0.49 | 3 | 2 | 0 |
| Q99459 | CDC5L | 1.63 | 1.14 | 0.49 | 0 | 5 | 0 |
| O43242 | PSMD3 | 1.83 | 1.11 | 0.49 | 0 | 7 | 0 |
| P42285 | SKIV2L2 | 1.4 | 1.06 | 0.49 | 2 | 1 | 0 |
| P62304 | SNRPE | 1.19 | 0.74 | 0.49 | 1 | 2 | 0 |
| Q02978 | SLC25A11 | 1.29 | 0.53 | 0.49 | 0 | 7 | 0 |
| P36542 | ATP5C1 | 2.31 | 1.9 | 0.48 | 0 | 8 | 0 |
| P51570 | GALK1 | 2.14 | 1.81 | 0.48 | 0 | 7 | 0 |
| Q9H7B2 | RPF2 | 2.04 | 1.75 | 0.48 | 5 | 0 | 0 |
| P35250 | RFC2 | 2.04 | 1.75 | 0.48 | 5 | 0 | 0 |
| P10909 | CLU | 1.98 | 1.72 | 0.48 | 0 | 6 | 0 |
| Q06210 | GFPT1 | 1.98 | 1.72 | 0.48 | 0 | 6 | 0 |
| Q13148 | TARDBP | 1.98 | 1.72 | 0.48 | 0 | 6 | 0 |
| P12004 | PCNA | 1.83 | 1.63 | 0.48 | 4 | 0 | 0 |
| P04899 | GNAI2 | 1.82 | 1.62 | 0.48 | 0 | 5 | 0 |
| P22695 | UQCRC2 | 1.82 | 1.62 | 0.48 | 0 | 5 | 0 |
| Q9P015 | MRPL15 | 1.65 | 1.52 | 0.48 | 0 | 4 | 0 |
| Q9P0M6 | H2AFY2 | 1.65 | 1.52 | 0.48 | 0 | 4 | 0 |
| P82675 | MRPS5 | 1.62 | 1.5 | 0.48 | 3 | 0 | 0 |
| Q86W42 | THOC6 | 1.62 | 1.5 | 0.48 | 3 | 0 | 0 |
| Q8WUM4 | PDCD6IP | 1.98 | 1.72 | 0.47 | 0 | 6 | 0 |
| Q13085 | ACACA | 1.58 | 1.56 | 0.47 | 2 | 1 | 0 |
| O75436 | VPS26A | 1.65 | 1.52 | 0.47 | 0 | 4 | 0 |
| P63000 | RAC1 | 1.65 | 1.52 | 0.47 | 0 | 4 | 0 |
| Q9Y6C9 | MTCH2 | 1.65 | 1.52 | 0.47 | 0 | 4 | 0 |
| Q13867 | BLMH | 1.62 | 1.5 | 0.47 | 3 | 0 | 0 |
| Q9HB71 | CACYBP | 1.49 | 1.41 | 0.47 | 0 | 3 | 0 |
| P04083 | ANXA1 | 1.49 | 1.41 | 0.47 | 0 | 3 | 0 |
| P00387 | CYB5R3 | 1.49 | 1.41 | 0.47 | 0 | 3 | 0 |
| O95299 | NDUFA10 | 1.49 | 1.41 | 0.47 | 0 | 3 | 0 |
| Q8NAV1 | PRPF38A | 1.49 | 1.41 | 0.47 | 0 | 3 | 0 |
| O75477 | ERLIN1 | 1.49 | 1.41 | 0.47 | 0 | 3 | 0 |
| P04181 | OAT | 1.44 | 1.26 | 0.47 | 0 | 3 | 0 |
| Q16540 | MRPL23 | 1.44 | 1.26 | 0.47 | 0 | 3 | 0 |
| P06493 | CDK1 | 1.44 | 1.26 | 0.47 | 0 | 3 | 0 |
| O75955 | FLOT1 | 1.65 | 1.52 | 0.46 | 0 | 4 | 0 |
| Q9Y5J1 | UTP18 | 1.62 | 1.5 | 0.46 | 3 | 0 | 0 |
| Q16795 | NDUFA9 | 1.49 | 1.41 | 0.46 | 0 | 3 | 0 |
| P24752 | ACAT1 | 1.49 | 1.41 | 0.46 | 0 | 3 | 0 |
| Q9Y2P8 | RCL1 | 1.49 | 1.41 | 0.46 | 0 | 3 | 0 |
| P00374 | DHFR | 1.41 | 1.35 | 0.46 | 2 | 0 | 0 |
| P24666 | ACP1 | 1.33 | 1.29 | 0.46 | 0 | 2 | 0 |
| Q14839 | CHD4 | 1.66 | 1 | 0.46 | 4 | 2 | 0 |
| O60841 | EIF5B | 1.42 | 0.94 | 0.46 | 0 | 4 | 0 |
| P62826 | RAN | 1.41 | 0.81 | 0.46 | 3 | 3 | 0 |
| P08133 | ANXA6 | 1.62 | 1.5 | 0.45 | 3 | 0 | 0 |
| O43615 | TIMM44 | 1.49 | 1.41 | 0.45 | 0 | 3 | 0 |
| P61421 | ATP6V0D1 | 1.49 | 1.41 | 0.45 | 0 | 3 | 0 |
| P62495 | ETF1 | 1.49 | 1.41 | 0.45 | 0 | 3 | 0 |
| Q8IXM3 | MRPL41 | 1.41 | 1.35 | 0.45 | 2 | 0 | 0 |
| O95433 | AHSA1 | 1.41 | 1.35 | 0.45 | 2 | 0 | 0 |
| O94905 | ERLIN2 | 1.41 | 1.35 | 0.45 | 2 | 0 | 0 |
| Q96QD9 | FYTTD1 | 1.41 | 1.35 | 0.45 | 2 | 0 | 0 |
| Q9NX46 | ADPRHL2 | 1.41 | 1.35 | 0.45 | 2 | 0 | 0 |
| O75223 | GGCT | 1.33 | 1.29 | 0.45 | 0 | 2 | 0 |

| | | | | | | | |
|---|---|---|---|---|---|---|---|
| Q00483 | NDUFA4 | 1.33 | 1.29 | 0.45 | 0 | 2 | 0 |
| Q96CW1 | AP2M1 | 1.34 | 1.12 | 0.45 | 2 | 0 | 0 |
| P27824 | CANX | 1.57 | 1.03 | 0.45 | 3 | 2 | 0 |
| Q92900 | UPF1 | 1.52 | 0.93 | 0.45 | 4 | 0 | 0 |
| Q16695 | HIST3H3 | 1.62 | 0.75 | 0.45 | 9 | 6 | 2 |
| P33992 | MCM5 | 1.62 | 1.5 | 0.44 | 3 | 0 | 0 |
| P19784 | CSNK2A2 | 1.41 | 1.35 | 0.44 | 2 | 0 | 0 |
| Q9NWU5 | MRPL22 | 1.41 | 1.35 | 0.44 | 2 | 0 | 0 |
| Q9NQG5 | RPRD1B | 1.41 | 1.35 | 0.44 | 2 | 0 | 0 |
| Q9BYC9 | MRPL20 | 1.41 | 1.35 | 0.44 | 2 | 0 | 0 |
| Q9Y277 | VDAC3 | 1.33 | 1.29 | 0.44 | 0 | 2 | 0 |
| P29992 | GNA11 | 1.29 | 1.15 | 0.44 | 0 | 2 | 0 |
| P12532 | CKMT1A;CKMT1B | 1.34 | 1.12 | 0.44 | 2 | 0 | 0 |
| P13667 | PDIA4 | 1.25 | 1.06 | 0.44 | 0 | 2 | 0 |
| P61163 | ACTR1A | 1.36 | 1.03 | 0.44 | 0 | 3 | 0 |
| Q13423 | NNT | 1.98 | 1.72 | 0.43 | 0 | 6 | 0 |
| P37837 | TALDO1 | 1.41 | 1.35 | 0.43 | 2 | 0 | 0 |
| Q8TCT9 | HM13 | 1.41 | 1.35 | 0.43 | 2 | 0 | 0 |
| P01591 | JCHAIN | 1.41 | 1.35 | 0.43 | 2 | 0 | 0 |
| Q9Y5R4 | HEMK1 | 1.33 | 1.29 | 0.43 | 0 | 2 | 0 |
| O00442 | RTCA | 1.33 | 1.29 | 0.43 | 0 | 2 | 0 |
| Q9H9L3 | ISG20L2 | 1.33 | 1.29 | 0.43 | 0 | 2 | 0 |
| P67775 | PPP2CA | 1.33 | 1.29 | 0.43 | 0 | 2 | 0 |
| Q5T653 | MRPL2 | 1.33 | 1.29 | 0.43 | 0 | 2 | 0 |
| Q96J01 | THOC3 | 1.33 | 1.29 | 0.43 | 0 | 2 | 0 |
| Q03701 | CEBPZ | 1.37 | 1.21 | 0.43 | 2 | 0 | 0 |
| Q9UN86 | G3BP2 | 1.27 | 0.95 | 0.43 | 2 | 0 | 0 |
| P48147 | PREP | 1.17 | 0.86 | 0.43 | 0 | 2 | 0 |
| O43143 | DHX15 | 1.99 | 0.83 | 0.43 | 11 | 13 | 0 |
| P43246 | MSH2 | 1.65 | 1.52 | 0.42 | 0 | 4 | 0 |
| P19367 | HK1 | 1.62 | 1.5 | 0.42 | 3 | 0 | 0 |
| Q6L8Q7 | PDE12 | 1.49 | 1.41 | 0.42 | 0 | 3 | 0 |
| P61619 | SEC61A1 | 1.41 | 1.35 | 0.42 | 2 | 0 | 0 |
| P27694 | RPA1 | 1.41 | 1.35 | 0.42 | 2 | 0 | 0 |
| P48651 | PTDSS1 | 1.33 | 1.29 | 0.42 | 0 | 2 | 0 |
| P61962 | DCAF7 | 1.33 | 1.29 | 0.42 | 0 | 2 | 0 |
| O14929 | HAT1 | 1.33 | 1.29 | 0.42 | 0 | 2 | 0 |
| P63241 | EIF5A | 1.29 | 1.15 | 0.42 | 0 | 2 | 0 |
| Q92552 | MRPS27 | 0.88 | 0.39 | 0.42 | 0 | 3 | 0 |
| A0FGR8 | ESYT2 | 1.62 | 1.5 | 0.41 | 3 | 0 | 0 |
| Q15645 | TRIP13 | 1.41 | 1.35 | 0.41 | 2 | 0 | 0 |
| Q8NFF5 | FLAD1 | 1.41 | 1.35 | 0.41 | 2 | 0 | 0 |
| P39748 | FEN1 | 1.33 | 1.29 | 0.41 | 0 | 2 | 0 |
| Q9Y2Z4 | YARS2 | 1.33 | 1.29 | 0.41 | 0 | 2 | 0 |
| O43813 | LANCL1 | 1.33 | 1.29 | 0.41 | 0 | 2 | 0 |
| Q1ED39 | KNOP1 | 1.33 | 1.29 | 0.41 | 0 | 2 | 0 |
| Q8TDD1 | DDX54 | 1.29 | 1.15 | 0.41 | 0 | 2 | 0 |
| Q99714 | HSD17B10 | 1.4 | 1.14 | 0.41 | 0 | 3 | 0 |
| P35611 | ADD1 | 1.25 | 1.04 | 0.41 | 0 | 2 | 0 |
| Q99623 | PHB2 | 2.4 | 1.01 | 0.41 | 11 | 10 | 0 |
| Q16630 | CPSF6 | 1.48 | 0.88 | 0.41 | 0 | 5 | 0 |
| Q99848 | EBNA1BP2 | 1.16 | 0.58 | 0.41 | 3 | 1 | 0 |
| Q16822 | PCK2 | 1.41 | 1.35 | 0.4 | 2 | 0 | 0 |
| O75390 | CS | 1.41 | 1.35 | 0.4 | 2 | 0 | 0 |
| Q8TDN6 | BRIX1 | 1.33 | 1.29 | 0.4 | 0 | 2 | 0 |
| Q96I24 | FUBP3 | 1.33 | 1.29 | 0.4 | 0 | 2 | 0 |
| P19525 | EIF2AK2 | 1.33 | 1.29 | 0.4 | 0 | 2 | 0 |
| O14979 | HNRNPDL | 1.86 | 1.24 | 0.4 | 3 | 4 | 0 |
| P62140 | PPP1CB | 1.33 | 1.1 | 0.4 | 2 | 0 | 0 |
| P63104 | YWHAZ | 1.44 | 1.01 | 0.4 | 1 | 3 | 0 |
| P41219 | PRPH | 1.23 | 0.87 | 0.4 | 2 | 0 | 0 |
| Q9BUQ8 | DDX23 | 1.01 | 0.55 | 0.4 | 2 | 0 | 0 |
| Q02539 | HIST1H1A | 1.25 | 0.44 | 0.4 | 13 | 9 | 0 |
| P46940 | IQGAP1 | 1.98 | 1.72 | 0.39 | 0 | 6 | 0 |
| Q8WUQ7 | CACTIN | 1.41 | 1.35 | 0.39 | 2 | 0 | 0 |
| O60678 | PRMT3 | 1.33 | 1.29 | 0.39 | 0 | 2 | 0 |
| Q9P2R7 | SUCLA2 | 1.33 | 1.29 | 0.39 | 0 | 2 | 0 |
| Q9H9A6 | LRRC40 | 1.33 | 1.29 | 0.39 | 0 | 2 | 0 |

| | | | | | | | |
|---|---|---|---|---|---|---|---|
| Q12873 | CHD3 | 1.4 | 1.14 | 0.39 | 0 | 3 | 0 |
| Q9NZ01 | TECR | 1.19 | 0.79 | 0.39 | 2 | 0 | 0 |
| Q12769 | NUP160 | 1.01 | 0.6 | 0.39 | 0 | 2 | 1 |
| P50454 | SERPINH1 | 0.94 | 0.51 | 0.39 | 0 | 2 | 1 |
| P08195 | SLC3A2 | 1.41 | 1.35 | 0.38 | 2 | 0 | 0 |
| Q96EK5 | KIF1BP | 1.33 | 1.29 | 0.38 | 0 | 2 | 0 |
| Q4G0J3 | LARP7 | 1.33 | 1.29 | 0.38 | 0 | 2 | 0 |
| A8MWD9 | SNRPGP15 | 1.29 | 0.99 | 0.38 | 2 | 0 | 0 |
| O75027 | ABCB7 | 1.41 | 1.35 | 0.37 | 2 | 0 | 0 |
| P28288 | ABCD3 | 1.33 | 1.29 | 0.37 | 0 | 2 | 0 |
| P78362 | SRPK2 | 1.15 | 0.81 | 0.37 | 0 | 2 | 0 |
| Q9Y265 | RUVBL1 | 1.1 | 0.53 | 0.37 | 0 | 4 | 0 |
| Q9Y5L0 | TNPO3 | 1.41 | 1.35 | 0.36 | 2 | 0 | 0 |
| P42224 | STAT1 | 1.33 | 1.29 | 0.36 | 0 | 2 | 0 |
| P54136 | RARS | 1.96 | 1.03 | 0.36 | 7 | 4 | 0 |
| P07910 | HNRNPC | 1.66 | 0.68 | 0.36 | 10 | 23 | 4 |
| Q13155 | AIMP2 | 1.21 | 0.68 | 0.36 | 0 | 4 | 0 |
| Q14498 | RBM39 | 1.44 | 0.67 | 0.36 | 4 | 4 | 0 |
| P25205 | MCM3 | 0.96 | 0.48 | 0.36 | 1 | 2 | 0 |
| Q9NXF1 | TEX10 | 1.49 | 1.41 | 0.35 | 0 | 3 | 0 |
| Q15020 | SART3 | 1.41 | 1.35 | 0.35 | 2 | 0 | 0 |
| Q9H8H0 | NOL11 | 1.41 | 1.35 | 0.35 | 2 | 0 | 0 |
| P18074 | ERCC2 | 1.33 | 1.29 | 0.35 | 0 | 2 | 0 |
| P26639 | TARS | 1.33 | 1.29 | 0.35 | 0 | 2 | 0 |
| Q5T9A4 | ATAD3B | 1.13 | 0.79 | 0.35 | 0 | 2 | 0 |
| P61513 | RPL37A | 1.13 | 0.65 | 0.35 | 0 | 3 | 0 |
| Q9H2U1 | DHX36 | 1.41 | 1.35 | 0.34 | 2 | 0 | 0 |
| Q96P70 | IPO9 | 1.41 | 1.35 | 0.34 | 2 | 0 | 0 |
| O60568 | PLOD3 | 1.33 | 1.29 | 0.34 | 0 | 2 | 0 |
| P62263 | RPS14 | 1.78 | 1.04 | 0.34 | 7 | 8 | 0 |
| O43795 | MYO1B | 1.14 | 0.81 | 0.34 | 0 | 2 | 0 |
| P35268 | RPL22 | 1.27 | 0.78 | 0.34 | 1 | 3 | 0 |
| O43175 | PHGDH | 1.18 | 0.89 | 0.33 | 0 | 2 | 0 |
| Q92522 | H1FX | 1.6 | 0.68 | 0.33 | 8 | 0 | 0 |
| P46013 | MKI67 | 2.47 | 1.99 | 0.32 | 0 | 9 | 0 |
| Q8IXT5 | RBM12B | 1.41 | 1.35 | 0.32 | 2 | 0 | 0 |
| Q9Y4W2 | LAS1L | 1.33 | 1.29 | 0.32 | 0 | 2 | 0 |
| P63173 | RPL38 | 1.31 | 0.81 | 0.32 | 1 | 3 | 0 |
| O75400 | PRPF40A | 1.33 | 1.29 | 0.31 | 0 | 2 | 0 |
| P17987 | TCP1 | 1.94 | 0.97 | 0.31 | 2 | 9 | 0 |
| Q01844 | EWSR1 | 1.24 | 0.82 | 0.31 | 1 | 2 | 0 |
| Q8N5C6 | SRBD1 | 1.33 | 1.29 | 0.3 | 0 | 2 | 0 |
| Q9H6R4 | NOL6 | 1.33 | 1.29 | 0.3 | 0 | 2 | 0 |
| P55265 | ADAR | 1.33 | 1.29 | 0.3 | 0 | 2 | 0 |
| P40227 | CCT6A | 1.2 | 0.64 | 0.3 | 0 | 4 | 0 |
| Q8N3C0 | ASCC3 | 1.49 | 1.41 | 0.28 | 0 | 3 | 0 |
| Q99570 | PIK3R4 | 1.41 | 1.35 | 0.28 | 2 | 0 | 0 |
| Q8WUM0 | NUP133 | 1.33 | 1.29 | 0.28 | 0 | 2 | 0 |
| Q9Y2A7 | NCKAP1 | 1.33 | 1.29 | 0.28 | 0 | 2 | 0 |
| Q02878 | RPL6 | 1.69 | 0.8 | 0.28 | 20 | 12 | 0 |
| Q93009 | USP7 | 1.33 | 1.29 | 0.27 | 0 | 2 | 0 |
| P20042 | EIF2S2 | 1.21 | 0.76 | 0.27 | 0 | 3 | 0 |
| P26358 | DNMT1 | 1.41 | 1.35 | 0.26 | 2 | 0 | 0 |
| Q9NU22 | MDN1 | 1.25 | 1.04 | 0.26 | 0 | 2 | 0 |
| Q3ZCQ8 | TIMM50 | 1.2 | 0.65 | 0.26 | 2 | 3 | 0 |
| Q9UQ35 | SRRM2 | 0.59 | 0.16 | 0.26 | 41 | 25 | 68 |
| P52701 | MSH6 | 1.33 | 1.29 | 0.25 | 0 | 2 | 0 |
| Q96AG4 | LRRC59 | 1.18 | 0.89 | 0.25 | 0 | 2 | 0 |
| P08240 | SRPRA | 1.13 | 0.78 | 0.25 | 0 | 2 | 0 |
| P01869 | P01869 | 0.54 | 0.15 | 0.25 | 37 | 28 | 77 |
| O15067 | PFAS | 1.33 | 1.29 | 0.24 | 0 | 2 | 0 |
| P20700 | LMNB1 | 1.59 | 1.06 | 0.23 | 3 | 2 | 0 |
| P27816 | MAP4 | 1.18 | 0.88 | 0.23 | 0 | 2 | 0 |
| Q96DI7 | SNRNP40 | 1.22 | 0.85 | 0.23 | 2 | 0 | 0 |
| P23246 | SFPQ | 1.91 | 0.84 | 0.23 | 9 | 8 | 0 |
| O00303 | EIF3F | 1.17 | 0.61 | 0.22 | 0 | 4 | 0 |
| P47897 | QARS | 1.81 | 1.01 | 0.21 | 4 | 9 | 0 |
| Q00610 | CLTC | 2 | 0.92 | 0.21 | 26 | 17 | 0 |

| | | | | | | | |
|---|---|---|---|---|---|---|---|
| Q9BUJ2 | HNRNPUL1 | 1.9 | 1.09 | 0.2 | 3 | 6 | 0 |
| Q58FF8 | HSP90AB2P | 1.56 | 0.81 | 0.2 | 5 | 11 | 0 |
| Q13547 | HDAC1 | 1.22 | 0.78 | 0.2 | 1 | 2 | 0 |
| Q9Y6M1 | IGF2BP2 | 0.98 | 0.54 | 0.2 | 1 | 3 | 0 |
| Q14966 | ZNF638 | 1.33 | 1.29 | 0.19 | 0 | 2 | 0 |
| Q93008 | USP9X | 1.41 | 1.35 | 0.18 | 2 | 0 | 0 |
| P50851 | LRBA | 1.41 | 1.35 | 0.18 | 2 | 0 | 0 |
| Q14669 | TRIP12 | 1.33 | 1.29 | 0.18 | 0 | 2 | 0 |
| E9PAV3 | NACA | 1.32 | 0.97 | 0.18 | 1 | 2 | 0 |
| P62195 | PSMC5 | 1.13 | 0.65 | 0.17 | 0 | 3 | 0 |
| P62861 | FAU | 0.91 | 0.44 | 0.17 | 2 | 0 | 0 |
| P08670 | VIM | 1.54 | 0.78 | 0.16 | 5 | 10 | 0 |
| P09874 | PARP1 | 1.96 | 1.16 | 0.15 | 22 | 15 | 1 |
| P62081 | RPS7 | 2.19 | 1.11 | 0.15 | 7 | 8 | 0 |
| Q58FF7 | HSP90AB3P | 1.48 | 0.58 | 0.15 | 4 | 12 | 0 |
| Q15008 | PSMD6 | 1.15 | 0.68 | 0.14 | 0 | 3 | 0 |
| P48643 | CCT5 | 1.1 | 0.41 | 0.14 | 0 | 8 | 0 |
| P26368 | U2AF2 | 0.78 | 0.36 | 0.14 | 0 | 2 | 2 |
| P13010 | XRCC5 | 2.27 | 1.18 | 0.13 | 19 | 16 | 0 |
| Q99832 | CCT7 | 1.41 | 0.58 | 0.12 | 10 | 9 | 3 |
| P18077 | RPL35A | 1.1 | 0.64 | 0.11 | 0 | 4 | 0 |
| P78371 | CCT2 | 1.07 | 0.53 | 0.11 | 2 | 3 | 0 |
| Q06265 | EXOSC9 | 0.85 | 0.39 | 0.11 | 2 | 0 | 0 |
| P62829 | RPL23 | 1.39 | 0.82 | 0.1 | 3 | 4 | 0 |
| Q9ULV4 | CORO1C | 1.05 | 0.65 | 0.1 | 0 | 2 | 0 |
| P62753 | RPS6 | 1.25 | 0.59 | 0.1 | 0 | 8 | 0 |
| P05455 | SSB | 1.05 | 0.5 | 0.1 | 3 | 3 | 0 |
| P62888 | RPL30 | 0.89 | 0.43 | 0.1 | 2 | 0 | 0 |
| P62277 | RPS13 | 1.47 | 0.86 | 0.09 | 3 | 6 | 0 |
| P08865 | RPSA | 1.03 | 0.58 | 0.09 | 2 | 1 | 0 |
| O43390 | HNRNPR | 1.22 | 0.57 | 0.09 | 11 | 9 | 0 |
| P46781 | RPS9 | 1.66 | 0.67 | 0.08 | 9 | 7 | 0 |
| O00232 | PSMD12 | 1.12 | 0.64 | 0.08 | 0 | 3 | 0 |
| Q15758 | SLC1A5 | 0.98 | 0.5 | 0.08 | 1 | 2 | 0 |
| Q66PJ3 | ARL6IP4 | 0.81 | 0.39 | 0.08 | 0 | 2 | 0 |
| Q9Y230 | RUVBL2 | 0.78 | 0.35 | 0.08 | 0 | 3 | 0 |
| P62280 | RPS11 | 0.67 | 0.23 | 0.08 | 1 | 6 | 0 |
| P42766 | RPL35 | 1.13 | 0.62 | 0.07 | 3 | 0 | 0 |
| P62899 | RPL31 | 1.23 | 0.68 | 0.06 | 4 | 5 | 0 |
| P46778 | RPL21 | 1.01 | 0.61 | 0.06 | 2 | 0 | 0 |
| P23526 | AHCY | 0.52 | 0.16 | 0.06 | 7 | 7 | 19 |
| Q9NZI8 | IGF2BP1 | 1.19 | 0.63 | 0.05 | 9 | 9 | 1 |
| O75369 | FLNB | 1.11 | 0.61 | 0.05 | 2 | 1 | 0 |
| Q71UI9 | H2AFV | 0.77 | 0.39 | 0.05 | 5 | 0 | 0 |
| P62857 | RPS28 | 0.82 | 0.45 | 0.04 | 2 | 1 | 0 |
| Q9BQG0 | MYBBP1A | 1.06 | 0.42 | 0.04 | 4 | 7 | 0 |
| P50991 | CCT4 | 1.37 | 0.71 | 0.03 | 2 | 4 | 0 |
| P84103 | SRSF3 | 1.16 | 0.65 | 0.03 | 2 | 4 | 0 |
| P10515 | DLAT | 1.13 | 0.59 | 0.03 | 2 | 3 | 0 |
| P62847 | RPS24 | 1.23 | 0.57 | 0.03 | 3 | 5 | 0 |
| P31689 | DNAJA1 | 1.06 | 0.56 | 0.03 | 0 | 3 | 0 |
| Q02543 | RPL18A | 0.98 | 0.56 | 0.03 | 0 | 2 | 0 |
| P08559 | PDHA1 | 0.98 | 0.56 | 0.03 | 2 | 2 | 0 |
| P32119 | PRDX2 | 1.09 | 0.55 | 0.03 | 5 | 2 | 0 |
| P62805 | HIST1H4A;HIST1H4B;HI. | 0.83 | 0.47 | 0.03 | 10 | 22 | 4 |
| P18124 | RPL7 | 0.84 | 0.32 | 0.03 | 5 | 14 | 6 |
| P62854 | RPS26 | 1.18 | 0.7 | 0.02 | 2 | 1 | 0 |
| Q92841 | DDX17 | 1.26 | 0.68 | 0.02 | 14 | 11 | 3 |
| P36578 | RPL4 | 1.2 | 0.52 | 0.02 | 15 | 5 | 0 |
| O75821 | EIF3G | 0.72 | 0.32 | 0.02 | 0 | 2 | 0 |
| O00159 | MYO1C | 0.75 | 0.27 | 0.02 | 7 | 3 | 0 |
| P11940 | PABPC1 | 1.74 | 1.1 | 0.01 | 17 | 18 | 0 |
| P46779 | RPL28 | 1.12 | 0.7 | 0.01 | 2 | 4 | 0 |
| P83731 | RPL24 | 1.13 | 0.63 | 0.01 | 4 | 3 | 0 |
| P47914 | RPL29 | 0.98 | 0.59 | 0.01 | 2 | 2 | 0 |
| Q96QV6 | HIST1H2AA | 1 | 0.54 | 0.01 | 6 | 8 | 2 |
| P42704 | LRPPRC | 1.44 | 0.52 | 0.01 | 9 | 4 | 0 |
| P52597 | HNRNPF | 0.83 | 0.52 | 0.01 | 1 | 5 | 0 |

| | | | | | | | |
|---|---|---|---|---|---|---|---|
| P12235 | SLC25A4 | 0.99 | 0.49 | 0.01 | 4 | 5 | 0 |
| Q9UNX3 | RPL26L1 | 1.04 | 0.48 | 0.01 | 1 | 5 | 0 |
| P62701 | RPS4X | 1 | 0.46 | 0.01 | 13 | 12 | 3 |
| P62241 | RPS8 | 0.84 | 0.45 | 0.01 | 7 | 2 | 0 |
| Q13347 | EIF3I | 1.15 | 0.44 | 0.01 | 5 | 9 | 0 |
| O00425 | IGF2BP3 | 0.73 | 0.42 | 0.01 | 0 | 2 | 0 |
| Q9Y262 | EIF3L | 0.91 | 0.39 | 0.01 | 3 | 3 | 0 |
| P60866 | RPS20 | 0.78 | 0.37 | 0.01 | 2 | 2 | 0 |
| Q14568 | HSP90AA2P | 0.72 | 0.36 | 0.01 | 0 | 4 | 0 |
| P08708 | RPS17 | 0.55 | 0.29 | 0.01 | 0 | 3 | 0 |
| P23528 | CFL1 | 0.66 | 0.29 | 0.01 | 2 | 5 | 0 |
| P35998 | PSMC2 | 0.67 | 0.26 | 0.01 | 0 | 3 | 0 |
| Q13206 | DDX10 | 1.37 | 1.37 | 0 | 1 | 1 | 0 |
| O94776 | MTA2 | 1.37 | 1.37 | 0 | 1 | 1 | 0 |
| Q8TBK6 | ZCCHC10 | 1.37 | 1.37 | 0 | 1 | 1 | 0 |
| O60762 | DPM1 | 1.37 | 1.37 | 0 | 1 | 1 | 0 |
| Q15181 | PPA1 | 1.37 | 1.37 | 0 | 1 | 1 | 0 |
| Q8TCS8 | PNPT1 | 1.37 | 1.37 | 0 | 1 | 1 | 0 |
| Q969P6 | TOP1MT | 1.37 | 1.37 | 0 | 1 | 1 | 0 |
| Q9UNX4 | WDR3 | 1.37 | 1.37 | 0 | 1 | 1 | 0 |
| Q9Y241 | HIGD1A | 1.37 | 1.37 | 0 | 1 | 1 | 0 |
| P01876 | IGHA1 | 1.37 | 1.37 | 0 | 1 | 1 | 0 |
| Q8NEJ9 | NGDN | 1.37 | 1.37 | 0 | 1 | 1 | 0 |
| O95478 | NSA2 | 1.37 | 1.37 | 0 | 1 | 1 | 0 |
| Q71RC2 | LARP4 | 1.37 | 1.37 | 0 | 1 | 1 | 0 |
| P35080 | PFN2 | 1.37 | 1.37 | 0 | 1 | 1 | 0 |
| Q86TJ2 | TADA2B | 1.37 | 1.37 | 0 | 1 | 1 | 0 |
| Q5C9Z4 | NOM1 | 1.37 | 1.37 | 0 | 1 | 1 | 0 |
| P61923 | COPZ1 | 1.37 | 1.37 | 0 | 1 | 1 | 0 |
| Q9BZE1 | MRPL37 | 1.37 | 1.37 | 0 | 1 | 1 | 0 |
| P61106 | RAB14 | 1.37 | 1.37 | 0 | 1 | 1 | 0 |
| O75964 | ATP5L | 1.37 | 1.37 | 0 | 1 | 1 | 0 |
| Q7L0Y3 | TRMT10C | 1.37 | 1.37 | 0 | 1 | 1 | 0 |
| P67812 | SEC11A | 1.37 | 1.37 | 0 | 1 | 1 | 0 |
| P40616 | ARL1 | 1.37 | 1.37 | 0 | 1 | 1 | 0 |
| P53007 | SLC25A1 | 1.37 | 1.37 | 0 | 1 | 1 | 0 |
| Q16650 | TBR1 | 1.37 | 1.37 | 0 | 1 | 1 | 0 |
| Q9BZF1 | OSBPL8 | 1.37 | 1.37 | 0 | 1 | 1 | 0 |
| Q9BSC4 | NOL10 | 1.37 | 1.37 | 0 | 1 | 1 | 0 |
| Q5T6V5 | C9orf64 | 1.37 | 1.37 | 0 | 1 | 1 | 0 |
| P56134 | ATP5J2 | 1.33 | 1.23 | 0 | 1 | 1 | 0 |
| Q12789 | GTF3C1 | 1.33 | 1.23 | 0 | 1 | 1 | 0 |
| Q15155 | NOMO1 | 1.33 | 1.23 | 0 | 1 | 1 | 0 |
| A6NEC2 | NPEPPSL1 | 1.21 | 1.19 | 0 | 1 | 0 | 0 |
| O15078 | CEP290 | 1.21 | 1.19 | 0 | 1 | 0 | 0 |
| P24928 | POLR2A | 1.21 | 1.19 | 0 | 1 | 0 | 0 |
| O75190 | DNAJB6 | 1.21 | 1.19 | 0 | 1 | 0 | 0 |
| Q9HB07 | C12orf10 | 1.21 | 1.19 | 0 | 1 | 0 | 0 |
| P78344 | EIF4G2 | 1.21 | 1.19 | 0 | 1 | 0 | 0 |
| Q9UIG0 | BAZ1B | 1.21 | 1.19 | 0 | 1 | 0 | 0 |
| Q04323 | UBXN1 | 1.21 | 1.19 | 0 | 1 | 0 | 0 |
| Q9GZR7 | DDX24 | 1.21 | 1.19 | 0 | 1 | 0 | 0 |
| P19086 | GNAZ | 1.21 | 1.19 | 0 | 1 | 0 | 0 |
| P61019 | RAB2A | 1.21 | 1.19 | 0 | 1 | 0 | 0 |
| Q5VV42 | CDKAL1 | 1.21 | 1.19 | 0 | 1 | 0 | 0 |
| Q8NC60 | NOA1 | 1.21 | 1.19 | 0 | 1 | 0 | 0 |
| Q6TDU7 | CASC1 | 1.21 | 1.19 | 0 | 1 | 0 | 0 |
| P50570 | DNM2 | 1.21 | 1.19 | 0 | 1 | 0 | 0 |
| Q9NPF4 | OSGEP | 1.21 | 1.19 | 0 | 1 | 0 | 0 |
| Q9BVS5 | TRMT61B | 1.21 | 1.19 | 0 | 1 | 0 | 0 |
| Q9UGP8 | SEC63 | 1.21 | 1.19 | 0 | 1 | 0 | 0 |
| Q96G21 | IMP4 | 1.21 | 1.19 | 0 | 1 | 0 | 0 |
| Q6FI81 | CIAPIN1 | 1.21 | 1.19 | 0 | 1 | 0 | 0 |
| Q9Y623 | MYH4 | 1.21 | 1.19 | 0 | 1 | 0 | 0 |
| Q6PGP7 | TTC37 | 1.21 | 1.19 | 0 | 1 | 0 | 0 |
| P46020 | PHKA1 | 1.21 | 1.19 | 0 | 1 | 0 | 0 |
| Q2M389 | KIAA1033 | 1.21 | 1.19 | 0 | 1 | 0 | 0 |
| Q13427 | PPIG | 1.21 | 1.19 | 0 | 1 | 0 | 0 |

| | | | | | | | |
|---|---|---|---|---|---|---|---|
| Q9UBM7 | DHCR7 | 1.21 | 1.19 | 0 | 1 | 0 | 0 |
| Q9NT62 | ATG3 | 1.21 | 1.19 | 0 | 1 | 0 | 0 |
| Q9Y3Z3 | SAMHD1 | 1.21 | 1.19 | 0 | 1 | 0 | 0 |
| P61225 | RAP2B | 1.21 | 1.19 | 0 | 1 | 0 | 0 |
| P52565 | ARHGDIA | 1.21 | 1.19 | 0 | 1 | 0 | 0 |
| P49959 | MRE11A | 1.21 | 1.19 | 0 | 1 | 0 | 0 |
| O43592 | XPOT | 1.21 | 1.19 | 0 | 1 | 0 | 0 |
| Q8WUK0 | PTPMT1 | 1.21 | 1.19 | 0 | 1 | 0 | 0 |
| Q7Z2W9 | MRPL21 | 1.21 | 1.19 | 0 | 1 | 0 | 0 |
| P35251 | RFC1 | 1.21 | 1.19 | 0 | 1 | 0 | 0 |
| Q66K14 | TBC1D9B | 1.21 | 1.19 | 0 | 1 | 0 | 0 |
| O14646 | CHD1 | 1.21 | 1.19 | 0 | 1 | 0 | 0 |
| Q9BZJ0 | CRNKL1 | 1.21 | 1.19 | 0 | 1 | 0 | 0 |
| Q6ZXV5 | TMTC3 | 1.21 | 1.19 | 0 | 1 | 0 | 0 |
| Q9BSJ2 | TUBGCP2 | 1.21 | 1.19 | 0 | 1 | 0 | 0 |
| Q6TFL3 | CCDC171 | 1.21 | 1.19 | 0 | 1 | 0 | 0 |
| Q86UK0 | ABCA12 | 1.21 | 1.19 | 0 | 1 | 0 | 0 |
| Q14137 | BOP1 | 1.21 | 1.19 | 0 | 1 | 0 | 0 |
| O95757 | HSPA4L | 1.21 | 1.19 | 0 | 1 | 0 | 0 |
| Q7Z6Z7 | HUWE1 | 1.21 | 1.19 | 0 | 1 | 0 | 0 |
| Q6P158 | DHX57 | 1.21 | 1.19 | 0 | 1 | 0 | 0 |
| Q8IWA0 | WDR75 | 1.21 | 1.19 | 0 | 1 | 0 | 0 |
| Q9H0U4 | RAB1B | 1.21 | 1.19 | 0 | 1 | 0 | 0 |
| P48729 | CSNK1A1 | 1.21 | 1.19 | 0 | 1 | 0 | 0 |
| Q92797 | SYMPK | 1.21 | 1.19 | 0 | 1 | 0 | 0 |
| Q7KZI7 | MARK2 | 1.21 | 1.19 | 0 | 1 | 0 | 0 |
| Q9NQZ2 | UTP3 | 1.21 | 1.19 | 0 | 1 | 0 | 0 |
| Q8N766 | EMC1 | 1.21 | 1.19 | 0 | 1 | 0 | 0 |
| Q96PE3 | INPP4A | 1.21 | 1.19 | 0 | 1 | 0 | 0 |
| Q6P3W7 | SCYL2 | 1.21 | 1.19 | 0 | 1 | 0 | 0 |
| P56556 | NDUFA6 | 1.21 | 1.19 | 0 | 1 | 0 | 0 |
| Q9NP92 | MRPS30 | 1.21 | 1.19 | 0 | 1 | 0 | 0 |
| Q8N1B4 | VPS52 | 1.21 | 1.19 | 0 | 1 | 0 | 0 |
| Q03252 | LMNB2 | 1.21 | 1.19 | 0 | 1 | 0 | 0 |
| Q9H2V7 | SPNS1 | 1.21 | 1.19 | 0 | 1 | 0 | 0 |
| Q9Y4W6 | AFG3L2 | 1.21 | 1.19 | 0 | 1 | 0 | 0 |
| P17661 | DES | 1.21 | 1.19 | 0 | 1 | 0 | 0 |
| P43304 | GPD2 | 1.21 | 1.19 | 0 | 1 | 0 | 0 |
| P0C221 | CCDC175 | 1.21 | 1.19 | 0 | 1 | 0 | 0 |
| Q9BVC6 | TMEM109 | 1.21 | 1.19 | 0 | 1 | 0 | 0 |
| P51531 | SMARCA2 | 1.21 | 1.19 | 0 | 1 | 0 | 0 |
| P10114 | RAP2A | 1.21 | 1.19 | 0 | 1 | 0 | 0 |
| Q8TEX9 | IPO4 | 1.21 | 1.19 | 0 | 1 | 0 | 0 |
| P09110 | ACAA1 | 1.21 | 1.19 | 0 | 1 | 0 | 0 |
| Q14139 | UBE4A | 1.21 | 1.19 | 0 | 1 | 0 | 0 |
| Q9BW92 | TARS2 | 1.21 | 1.19 | 0 | 1 | 0 | 0 |
| P25685 | DNAJB1 | 1.21 | 1.19 | 0 | 1 | 0 | 0 |
| Q6ZRS2 | SRCAP | 1.21 | 1.19 | 0 | 1 | 0 | 0 |
| Q8WVX9 | FAR1 | 1.21 | 1.19 | 0 | 1 | 0 | 0 |
| Q9UIF9 | BAZ2A | 1.21 | 1.19 | 0 | 1 | 0 | 0 |
| P40925 | MDH1 | 1.21 | 1.19 | 0 | 1 | 0 | 0 |
| Q9UBU9 | NXF1 | 1.21 | 1.19 | 0 | 1 | 0 | 0 |
| Q86XP3 | DDX42 | 1.21 | 1.19 | 0 | 1 | 0 | 0 |
| Q16891 | IMMT | 1.21 | 1.19 | 0 | 1 | 0 | 0 |
| Q92945 | KHSRP | 1.21 | 1.19 | 0 | 1 | 0 | 0 |
| Q13619 | CUL4A | 1.21 | 1.19 | 0 | 1 | 0 | 0 |
| Q13617 | CUL2 | 1.21 | 1.19 | 0 | 1 | 0 | 0 |
| Q8N8A6 | DDX51 | 1.21 | 1.19 | 0 | 1 | 0 | 0 |
| Q02241 | KIF23 | 1.21 | 1.19 | 0 | 1 | 0 | 0 |
| Q9NPE3 | NOP10 | 1.21 | 1.19 | 0 | 1 | 0 | 0 |
| Q7Z4Q2 | HEATR3 | 1.21 | 1.19 | 0 | 1 | 0 | 0 |
| Q86X67 | NUDT13 | 1.21 | 1.19 | 0 | 1 | 0 | 0 |
| Q9ULT8 | HECTD1 | 1.21 | 1.19 | 0 | 1 | 0 | 0 |
| Q8IXI1 | RHOT2 | 1.21 | 1.19 | 0 | 1 | 0 | 0 |
| P33981 | TTK | 1.21 | 1.19 | 0 | 1 | 0 | 0 |
| Q9ULK4 | MED23 | 1.21 | 1.19 | 0 | 1 | 0 | 0 |
| P49760 | CLK2 | 1.21 | 1.19 | 0 | 1 | 0 | 0 |
| Q07864 | POLE | 1.21 | 1.19 | 0 | 1 | 0 | 0 |

| | | | | | | | |
|---|---|---|---|---|---|---|---|
| P22694 | PRKACB | 1.21 | 1.19 | 0 | 1 | 0 | 0 |
| P84085 | ARF5 | 1.16 | 1.15 | 0 | 0 | 1 | 0 |
| Q5VYK3 | ECM29 | 1.16 | 1.15 | 0 | 0 | 1 | 0 |
| Q9Y2R5 | MRPS17 | 1.16 | 1.15 | 0 | 0 | 1 | 0 |
| Q9Y2R4 | DDX52 | 1.16 | 1.15 | 0 | 0 | 1 | 0 |
| P30049 | ATP5D | 1.16 | 1.15 | 0 | 0 | 1 | 0 |
| Q9C0C9 | UBE2O | 1.16 | 1.15 | 0 | 0 | 1 | 0 |
| Q9Y5P6 | GMPPB | 1.16 | 1.15 | 0 | 0 | 1 | 0 |
| P18085 | ARF4 | 1.16 | 1.15 | 0 | 0 | 1 | 0 |
| Q96CS3 | FAF2 | 1.16 | 1.15 | 0 | 0 | 1 | 0 |
| Q9H5V9 | CXorf56 | 1.16 | 1.15 | 0 | 0 | 1 | 0 |
| Q9NX24 | NHP2 | 1.16 | 1.15 | 0 | 0 | 1 | 0 |
| Q9Y450 | HBS1L | 1.16 | 1.15 | 0 | 0 | 1 | 0 |
| Q8NFH5 | NUP35 | 1.16 | 1.15 | 0 | 0 | 1 | 0 |
| P09001 | MRPL3 | 1.16 | 1.15 | 0 | 0 | 1 | 0 |
| O95071 | UBR5 | 1.16 | 1.15 | 0 | 0 | 1 | 0 |
| Q14807 | KIF22 | 1.16 | 1.15 | 0 | 0 | 1 | 0 |
| Q14202 | ZMYM3 | 1.16 | 1.15 | 0 | 0 | 1 | 0 |
| P09960 | LTA4H | 1.16 | 1.15 | 0 | 0 | 1 | 0 |
| Q9UNN5 | FAF1 | 1.16 | 1.15 | 0 | 0 | 1 | 0 |
| P30084 | ECHS1 | 1.16 | 1.15 | 0 | 0 | 1 | 0 |
| Q9BSV6 | TSEN34 | 1.16 | 1.15 | 0 | 0 | 1 | 0 |
| P04040 | CAT | 1.16 | 1.15 | 0 | 0 | 1 | 0 |
| Q29RF7 | PDS5A | 1.16 | 1.15 | 0 | 0 | 1 | 0 |
| Q9Y2G8 | DNAJC16 | 1.16 | 1.15 | 0 | 0 | 1 | 0 |
| Q9Y3Y2 | CHTOP | 1.16 | 1.15 | 0 | 0 | 1 | 0 |
| P99999 | CYCS | 1.16 | 1.15 | 0 | 0 | 1 | 0 |
| O95168 | NDUFB4 | 1.16 | 1.15 | 0 | 0 | 1 | 0 |
| Q16352 | INA | 1.16 | 1.15 | 0 | 0 | 1 | 0 |
| O60502 | MGEA5 | 1.16 | 1.15 | 0 | 0 | 1 | 0 |
| Q7Z6B0 | CCDC91 | 1.16 | 1.15 | 0 | 0 | 1 | 0 |
| Q6P6C2 | ALKBH5 | 1.16 | 1.15 | 0 | 0 | 1 | 0 |
| Q8WUY1 | THEM6 | 1.16 | 1.15 | 0 | 0 | 1 | 0 |
| Q9NQS7 | INCENP | 1.16 | 1.15 | 0 | 0 | 1 | 0 |
| Q9NPJ3 | ACOT13 | 1.16 | 1.15 | 0 | 0 | 1 | 0 |
| Q6YN16 | HSDL2 | 1.16 | 1.15 | 0 | 0 | 1 | 0 |
| P41240 | CSK | 1.16 | 1.15 | 0 | 0 | 1 | 0 |
| Q5M9Q1 | NKAPL | 1.16 | 1.15 | 0 | 0 | 1 | 0 |
| Q9Y536 | PPIAL4A | 1.16 | 1.15 | 0 | 0 | 1 | 0 |
| P00846 | MT-ATP6 | 1.16 | 1.15 | 0 | 0 | 1 | 0 |
| Q9H9P8 | L2HGDH | 1.16 | 1.15 | 0 | 0 | 1 | 0 |
| Q6PK04 | CCDC137 | 1.16 | 1.15 | 0 | 0 | 1 | 0 |
| Q9HCD5 | NCOA5 | 1.16 | 1.15 | 0 | 0 | 1 | 0 |
| Q9Y6D5 | ARFGEF2 | 1.16 | 1.15 | 0 | 0 | 1 | 0 |
| Q9NRN7 | AASDHPPT | 1.16 | 1.15 | 0 | 0 | 1 | 0 |
| O75396 | SEC22B | 1.16 | 1.15 | 0 | 0 | 1 | 0 |
| Q96AC1 | FERMT2 | 1.16 | 1.15 | 0 | 0 | 1 | 0 |
| Q13576 | IQGAP2 | 1.16 | 1.15 | 0 | 0 | 1 | 0 |
| Q9H993 | ARMT1 | 1.16 | 1.15 | 0 | 0 | 1 | 0 |
| P17858 | PFKL | 1.16 | 1.15 | 0 | 0 | 1 | 0 |
| Q13131 | PRKAA1 | 1.16 | 1.15 | 0 | 0 | 1 | 0 |
| P52735 | VAV2 | 1.16 | 1.15 | 0 | 0 | 1 | 0 |
| Q9GZK3 | OR2B2 | 1.16 | 1.15 | 0 | 0 | 1 | 0 |
| Q96TA2 | YME1L1 | 1.16 | 1.15 | 0 | 0 | 1 | 0 |
| P59998 | ARPC4 | 1.16 | 1.15 | 0 | 0 | 1 | 0 |
| Q02218 | OGDH | 1.16 | 1.15 | 0 | 0 | 1 | 0 |
| Q8TD19 | NEK9 | 1.16 | 1.15 | 0 | 0 | 1 | 0 |
| Q16512 | PKN1 | 1.16 | 1.15 | 0 | 0 | 1 | 0 |
| Q16513 | PKN2 | 1.16 | 1.15 | 0 | 0 | 1 | 0 |
| Q10570 | CPSF1 | 1.16 | 1.15 | 0 | 0 | 1 | 0 |
| Q3SXM5 | HSDL1 | 1.16 | 1.15 | 0 | 0 | 1 | 0 |
| Q9H0U3 | MAGT1 | 1.16 | 1.15 | 0 | 0 | 1 | 0 |
| P60953 | CDC42 | 1.16 | 1.15 | 0 | 0 | 1 | 0 |
| P46459 | NSF | 1.16 | 1.15 | 0 | 0 | 1 | 0 |
| P42261 | GRIA1 | 1.16 | 1.15 | 0 | 0 | 1 | 0 |
| Q14318 | FKBP8 | 1.16 | 1.15 | 0 | 0 | 1 | 0 |
| Q4G0F5 | VPS26B | 1.16 | 1.15 | 0 | 0 | 1 | 0 |
| Q8IWS0 | PHF6 | 1.16 | 1.15 | 0 | 0 | 1 | 0 |

| | | | | | | | |
|---|---|---|---|---|---|---|---|
| Q9NRX1 | PNO1 | 1.16 | 1.15 | 0 | 0 | 1 | 0 |
| P49641 | MAN2A2 | 1.16 | 1.15 | 0 | 0 | 1 | 0 |
| Q9NYH9 | UTP6 | 1.16 | 1.15 | 0 | 0 | 1 | 0 |
| P26196 | DDX6 | 1.16 | 1.15 | 0 | 0 | 1 | 0 |
| Q7L014 | DDX46 | 1.16 | 1.15 | 0 | 0 | 1 | 0 |
| Q6P1J9 | CDC73 | 1.16 | 1.15 | 0 | 0 | 1 | 0 |
| Q5XPI4 | RNF123 | 1.16 | 1.15 | 0 | 0 | 1 | 0 |
| P18583 | SON | 1.16 | 1.15 | 0 | 0 | 1 | 0 |
| P35237 | SERPINB6 | 1.16 | 1.15 | 0 | 0 | 1 | 0 |
| Q6IAN0 | DHRS7B | 1.16 | 1.15 | 0 | 0 | 1 | 0 |
| Q8IYL2 | TRMT44 | 1.16 | 1.15 | 0 | 0 | 1 | 0 |
| Q16610 | ECM1 | 1.16 | 1.15 | 0 | 0 | 1 | 0 |
| Q5T3I0 | GPATCH4 | 1.16 | 1.15 | 0 | 0 | 1 | 0 |
| Q8TC12 | RDH11 | 1.16 | 1.15 | 0 | 0 | 1 | 0 |
| Q9H936 | SLC25A22 | 1.16 | 1.15 | 0 | 0 | 1 | 0 |
| Q13045 | FLII | 1.16 | 1.15 | 0 | 0 | 1 | 0 |
| Q12824 | SMARCB1 | 1.16 | 1.15 | 0 | 0 | 1 | 0 |
| Q9HCY8 | S100A14 | 1.16 | 1.15 | 0 | 0 | 1 | 0 |
| P60763 | RAC3 | 1.16 | 1.15 | 0 | 0 | 1 | 0 |
| Q6UW68 | TMEM205 | 1.16 | 1.15 | 0 | 0 | 1 | 0 |
| Q15392 | DHCR24 | 1.16 | 1.15 | 0 | 0 | 1 | 0 |
| O15144 | ARPC2 | 1.16 | 1.15 | 0 | 0 | 1 | 0 |
| Q5UIP0 | RIF1 | 1.16 | 1.15 | 0 | 0 | 1 | 0 |
| Q9NTK5 | OLA1 | 1.16 | 1.15 | 0 | 0 | 1 | 0 |
| Q86YT6 | MIB1 | 1.16 | 1.15 | 0 | 0 | 1 | 0 |
| Q92947 | GCDH | 1.16 | 1.15 | 0 | 0 | 1 | 0 |
| Q9BW27 | NUP85 | 1.16 | 1.15 | 0 | 0 | 1 | 0 |
| O00170 | AIP | 1.16 | 1.15 | 0 | 0 | 1 | 0 |
| P55769 | SNU13 | 1.16 | 1.15 | 0 | 0 | 1 | 0 |
| O43776 | NARS | 1.16 | 1.15 | 0 | 0 | 1 | 0 |
| Q96P48 | ARAP1 | 1.16 | 1.15 | 0 | 0 | 1 | 0 |
| B7ZAQ6 | GPR89A | 1.16 | 1.15 | 0 | 0 | 1 | 0 |
| Q58FF3 | HSP90B2P | 1.16 | 1.15 | 0 | 0 | 1 | 0 |
| P48444 | ARCN1 | 1.16 | 1.15 | 0 | 0 | 1 | 0 |
| Q9NV31 | IMP3 | 1.16 | 1.15 | 0 | 0 | 1 | 0 |
| Q5VWG9 | TAF3 | 1.16 | 1.15 | 0 | 0 | 1 | 0 |
| P07954 | FH | 1.16 | 1.15 | 0 | 0 | 1 | 0 |
| Q99575 | POP1 | 1.16 | 1.15 | 0 | 0 | 1 | 0 |
| O75153 | CLUH | 1.16 | 1.15 | 0 | 0 | 1 | 0 |
| P19387 | POLR2C | 1.16 | 1.15 | 0 | 0 | 1 | 0 |
| O14792 | HS3ST1 | 1.16 | 1.15 | 0 | 0 | 1 | 0 |
| P08758 | ANXA5 | 1.16 | 1.15 | 0 | 0 | 1 | 0 |
| P31327 | CPS1 | 1.16 | 1.15 | 0 | 0 | 1 | 0 |
| P52434 | POLR2H | 1.16 | 1.15 | 0 | 0 | 1 | 0 |
| A8MXY4 | ZNF99 | 1.16 | 1.15 | 0 | 0 | 1 | 0 |
| Q15120 | PDK3 | 1.16 | 1.15 | 0 | 0 | 1 | 0 |
| Q96FW1 | OTUB1 | 1.16 | 1.15 | 0 | 0 | 1 | 0 |
| Q9H3N1 | TMX1 | 1.16 | 1.15 | 0 | 0 | 1 | 0 |
| Q6P090 | CLK4 | 1.16 | 1.15 | 0 | 0 | 1 | 0 |
| P43897 | TSFM | 1.16 | 1.15 | 0 | 0 | 1 | 0 |
| Q92973 | TNPO1 | 1.16 | 1.15 | 0 | 0 | 1 | 0 |
| Q86U42 | PABPN1 | 1.29 | 1.11 | 0 | 1 | 1 | 0 |
| O15260 | SURF4 | 1.29 | 1.11 | 0 | 1 | 1 | 0 |
| Q13242 | SRSF9 | 1.29 | 1.11 | 0 | 1 | 1 | 0 |
| P36873 | PPP1CC | 1.29 | 1.11 | 0 | 1 | 1 | 0 |
| Q10567 | AP1B1 | 1.17 | 1.07 | 0 | 1 | 0 | 0 |
| Q14692 | BMS1 | 1.17 | 1.07 | 0 | 1 | 0 | 0 |
| P61981 | YWHAG | 1.13 | 1.03 | 0 | 0 | 1 | 0 |
| P12814 | ACTN1 | 1.13 | 1.03 | 0 | 0 | 1 | 0 |
| Q9Y4P3 | TBL2 | 1.13 | 1.03 | 0 | 0 | 1 | 0 |
| O95831 | AIFM1 | 1.13 | 1.03 | 0 | 0 | 1 | 0 |
| P49406 | MRPL19 | 1.13 | 1.03 | 0 | 0 | 1 | 0 |
| P27348 | YWHAQ | 1.13 | 1.03 | 0 | 0 | 1 | 0 |
| Q04917 | YWHAH | 1.13 | 1.03 | 0 | 0 | 1 | 0 |
| Q14011 | CIRBP | 1.13 | 1.03 | 0 | 0 | 1 | 0 |
| Q8NBX0 | SCCPDH | 1.13 | 1.03 | 0 | 0 | 1 | 0 |
| P63010 | AP2B1 | 1.25 | 1.02 | 0 | 1 | 1 | 0 |
| Q9BPX3 | NCAPG | 1.14 | 0.98 | 0 | 1 | 0 | 0 |

| | | | | | | | |
|---|---|---|---|---|---|---|---|
| Q16637 | SMN1;SMN2 | 1.1 | 0.95 | 0 | 0 | 1 | 0 |
| Q14331 | FRG1 | 1.1 | 0.95 | 0 | 0 | 1 | 0 |
| O00541 | PES1 | 1.09 | 0.93 | 0 | 0 | 1 | 0 |
| Q00839 | HNRNPU | 1.47 | 0.87 | 0 | 23 | 24 | 0 |
| Q6PL18 | ATAD2 | 1.18 | 0.86 | 0 | 1 | 1 | 0 |
| Q9Y383 | LUC7L2 | 1.08 | 0.82 | 0 | 1 | 0 | 0 |
| Q04837 | SSBP1 | 1.07 | 0.81 | 0 | 1 | 0 | 0 |
| Q96PU8 | QKI | 1.06 | 0.8 | 0 | 1 | 0 | 0 |
| Q9NQ29 | LUC7L | 1.04 | 0.8 | 0 | 0 | 1 | 0 |
| P25789 | PSMA4 | 1.03 | 0.79 | 0 | 0 | 1 | 0 |
| Q13601 | KRR1 | 1.03 | 0.79 | 0 | 0 | 1 | 0 |
| P31946 | YWHAB | 1.05 | 0.76 | 0 | 1 | 0 | 0 |
| P63220 | RPS21 | 1.01 | 0.75 | 0 | 0 | 1 | 0 |
| A6NHR9 | SMCHD1 | 1.04 | 0.75 | 0 | 1 | 0 | 0 |
| P51398 | DAP3 | 1.01 | 0.74 | 0 | 0 | 1 | 0 |
| Q92769 | HDAC2 | 1.03 | 0.73 | 0 | 1 | 0 | 0 |
| Q71UM5 | RPS27L | 1.1 | 0.72 | 0 | 1 | 1 | 0 |
| P35606 | COPB2 | 1.02 | 0.71 | 0 | 1 | 0 | 0 |
| P15880 | RPS2 | 1.2 | 0.7 | 0 | 6 | 4 | 0 |
| Q9UG63 | ABCF2 | 1.08 | 0.7 | 0 | 1 | 1 | 0 |
| Q49A26 | GLYR1 | 1.01 | 0.69 | 0 | 1 | 0 | 0 |
| P62424 | RPL7A | 1.28 | 0.68 | 0 | 7 | 8 | 0 |
| Q96SB4 | SRPK1 | 0.98 | 0.68 | 0 | 0 | 1 | 0 |
| O15397 | IPO8 | 1 | 0.67 | 0 | 1 | 0 | 0 |
| P35249 | RFC4 | 0.97 | 0.67 | 0 | 0 | 1 | 0 |
| Q00341 | HDLBP | 1 | 0.67 | 0 | 1 | 0 | 0 |
| Q7L2H7 | EIF3M | 0.99 | 0.66 | 0 | 1 | 0 | 0 |
| Q13642 | FHL1 | 0.95 | 0.64 | 0 | 0 | 1 | 0 |
| P31150 | GDI1 | 0.97 | 0.63 | 0 | 1 | 0 | 0 |
| Q15233 | NONO | 1.2 | 0.6 | 0 | 6 | 4 | 0 |
| P62249 | RPS16 | 0.98 | 0.59 | 0 | 7 | 5 | 0 |
| Q07020 | RPL18 | 0.97 | 0.58 | 0 | 4 | 5 | 0 |
| Q9Y3B4 | SF3B6 | 0.92 | 0.58 | 0 | 0 | 1 | 0 |
| P07814 | EPRS | 1.38 | 0.57 | 0 | 7 | 10 | 0 |
| Q9NQ39 | RPS10P5 | 0.91 | 0.57 | 0 | 0 | 1 | 0 |
| Q9H4L4 | SENP3 | 0.92 | 0.56 | 0 | 1 | 0 | 1 |
| Q08211 | DHX9 | 1.17 | 0.55 | 0 | 16 | 21 | 0 |
| P09661 | SNRPA1 | 0.91 | 0.55 | 0 | 1 | 0 | 0 |
| P17844 | DDX5 | 1.06 | 0.55 | 0 | 8 | 8 | 0 |
| O00231 | PSMD11 | 0.89 | 0.54 | 0 | 0 | 1 | 0 |
| O95292 | VAPB | 0.89 | 0.54 | 0 | 0 | 1 | 1 |
| P28072 | PSMB6 | 0.91 | 0.54 | 0 | 1 | 0 | 0 |
| P31943 | HNRNPH1 | 0.89 | 0.52 | 0 | 7 | 5 | 0 |
| P62750 | RPL23A | 1.05 | 0.51 | 0 | 4 | 7 | 0 |
| O60506 | SYNCRIP | 0.8 | 0.51 | 0 | 7 | 6 | 0 |
| P38646 | HSPA9 | 1.04 | 0.51 | 0 | 6 | 13 | 0 |
| P10809 | HSPD1 | 1.12 | 0.51 | 0 | 10 | 14 | 0 |
| P56537 | EIF6 | 0.85 | 0.5 | 0 | 1 | 0 | 0 |
| P49207 | RPL34 | 0.84 | 0.5 | 0 | 1 | 0 | 0 |
| P62258 | YWHAE | 0.97 | 0.49 | 0 | 1 | 2 | 0 |
| P14678 | SNRPB | 1.04 | 0.48 | 0 | 2 | 5 | 0 |
| Q16629 | SRSF7 | 0.85 | 0.47 | 0 | 3 | 4 | 0 |
| P32969 | RPL9P7;RPL9P8;RPL9P | 0.88 | 0.45 | 0 | 1 | 5 | 0 |
| P19338 | NCL | 0.69 | 0.45 | 0 | 16 | 31 | 0 |
| Q32P51 | HNRNPA1L2 | 0.83 | 0.45 | 0 | 12 | 10 | 0 |
| P62266 | RPS23 | 0.93 | 0.44 | 0 | 1 | 5 | 0 |
| P51991 | HNRNPA3 | 0.83 | 0.44 | 0 | 3 | 8 | 0 |
| P62316 | SNRPD2 | 0.84 | 0.43 | 0 | 2 | 4 | 0 |
| Q99729 | HNRNPAB | 0.67 | 0.43 | 0 | 3 | 3 | 0 |
| Q9Y5Q9 | GTF3C3 | 0.8 | 0.43 | 0 | 0 | 1 | 0 |
| P61626 | LYZ | 0.85 | 0.42 | 0 | 1 | 1 | 2 |
| P33778 | HIST1H2BB | 0.83 | 0.42 | 0 | 8 | 10 | 3 |
| P50416 | CPT1A | 0.85 | 0.42 | 0 | 1 | 1 | 2 |
| P62851 | RPS25 | 0.81 | 0.42 | 0 | 3 | 3 | 0 |
| P12956 | XRCC6 | 0.89 | 0.42 | 0 | 7 | 20 | 0 |
| P35637 | FUS | 0.88 | 0.42 | 0 | 2 | 0 | 0 |
| P62913 | RPL11 | 0.72 | 0.41 | 0 | 2 | 5 | 0 |
| Q13151 | HNRNPA0 | 0.92 | 0.41 | 0 | 3 | 6 | 0 |

| | | | | | | | |
|---|---|---|---|---|---|---|---|
| P37108 | SRP14 | 0.78 | 0.41 | 0 | 0 | 1 | 0 |
| P68431 | HIST1H3A;HIST1H3B;HI. | 0.76 | 0.41 | 0 | 0 | 2 | 0 |
| P11021 | HSPA5 | 0.75 | 0.4 | 0 | 4 | 17 | 0 |
| Q15046 | KARS | 0.69 | 0.39 | 0 | 2 | 0 | 0 |
| Q13428 | TCOF1 | 0.81 | 0.39 | 0 | 1 | 1 | 0 |
| P05141 | SLC25A5 | 0.85 | 0.39 | 0 | 5 | 6 | 0 |
| P46783 | RPS10 | 0.75 | 0.39 | 0 | 1 | 0 | 0 |
| P35580 | MYH10 | 1.1 | 0.39 | 0 | 13 | 13 | 0 |
| Q12906 | ILF3 | 0.83 | 0.38 | 0 | 15 | 23 | 1 |
| P46782 | RPS5 | 0.64 | 0.38 | 0 | 1 | 1 | 0 |
| P23396 | RPS3 | 0.66 | 0.38 | 0 | 7 | 14 | 0 |
| P34931 | HSPA1L | 0.58 | 0.38 | 0 | 9 | 13 | 0 |
| P46777 | RPL5 | 0.8 | 0.38 | 0 | 7 | 8 | 0 |
| P06576 | ATP5B | 0.81 | 0.37 | 0 | 9 | 10 | 0 |
| Q14103 | HNRNPD | 0.71 | 0.36 | 0 | 1 | 5 | 0 |
| O00411 | POLRMT | 0.72 | 0.36 | 0 | 0 | 1 | 0 |
| P62333 | PSMC6 | 0.74 | 0.36 | 0 | 1 | 0 | 0 |
| P14866 | HNRNPL | 0.57 | 0.35 | 0 | 7 | 6 | 0 |
| P62273 | RPS29 | 0.75 | 0.35 | 0 | 1 | 1 | 0 |
| Q15029 | EFTUD2 | 0.86 | 0.35 | 0 | 3 | 0 | 0 |
| Q13310 | PABPC4 | 0.63 | 0.35 | 0 | 0 | 5 | 0 |
| P25705 | ATP5A1 | 0.66 | 0.35 | 0 | 5 | 6 | 0 |
| P62269 | RPS18 | 0.61 | 0.34 | 0 | 3 | 9 | 0 |
| P04908 | HIST1H2AB;HIST1H2AE | 0.64 | 0.34 | 0 | 5 | 4 | 0 |
| P62917 | RPL8 | 0.9 | 0.34 | 0 | 0 | 8 | 0 |
| P62244 | RPS15A | 0.75 | 0.34 | 0 | 0 | 9 | 0 |
| P11142 | HSPA8 | 0.59 | 0.34 | 0 | 15 | 16 | 1 |
| P26641 | EEF1G | 0.76 | 0.33 | 0 | 3 | 12 | 0 |
| P26599 | PTBP1 | 0.75 | 0.33 | 0 | 4 | 7 | 0 |
| Q01130 | SRSF2 | 0.79 | 0.33 | 0 | 1 | 3 | 0 |
| P52272 | HNRNPM | 0.77 | 0.33 | 0 | 16 | 22 | 0 |
| P62314 | SNRPD1 | 0.67 | 0.32 | 0 | 1 | 4 | 0 |
| P50395 | GDI2 | 0.77 | 0.32 | 0 | 0 | 3 | 0 |
| P68104 | EEF1A1 | 0.55 | 0.32 | 0 | 11 | 11 | 0 |
| P84098 | RPL19 | 0.62 | 0.32 | 0 | 3 | 1 | 0 |
| Q13243 | SRSF5 | 0.63 | 0.32 | 0 | 0 | 1 | 0 |
| P39023 | RPL3 | 0.62 | 0.31 | 0 | 3 | 9 | 0 |
| P61353 | RPL27 | 0.57 | 0.31 | 0 | 3 | 3 | 0 |
| P08621 | SNRNP70 | 0.67 | 0.3 | 0 | 1 | 0 | 0 |
| Q13283 | G3BP1 | 0.66 | 0.3 | 0 | 0 | 1 | 0 |
| P30050 | RPL12 | 0.58 | 0.3 | 0 | 2 | 5 | 0 |
| P06748 | NPM1 | 0.56 | 0.29 | 0 | 14 | 9 | 6 |
| P14649 | MYL6B | 0.58 | 0.29 | 0 | 1 | 0 | 0 |
| Q06830 | PRDX1 | 0.64 | 0.29 | 0 | 6 | 1 | 0 |
| P54652 | HSPA2 | 0.52 | 0.29 | 0 | 7 | 2 | 0 |
| P08238 | HSP90AB1 | 0.54 | 0.29 | 0 | 7 | 7 | 2 |
| B4DY08 | HNRNPC | 0.67 | 0.29 | 0 | 3 | 2 | 0 |
| Q8IUE6 | HIST2H2AB | 0.49 | 0.28 | 0 | 0 | 1 | 0 |
| P56192 | MARS | 0.65 | 0.28 | 0 | 3 | 4 | 0 |
| P07900 | HSP90AA1 | 0.64 | 0.27 | 0 | 11 | 9 | 5 |
| O00571 | DDX3X | 0.6 | 0.27 | 0 | 1 | 3 | 0 |
| P61247 | RPS3A | 0.54 | 0.27 | 0 | 7 | 4 | 0 |
| Q8N163 | CCAR2 | 0.69 | 0.26 | 0 | 3 | 2 | 0 |
| P52907 | CAPZA1 | 0.67 | 0.26 | 0 | 4 | 1 | 0 |
| Q96A08 | HIST1H2BA | 0.58 | 0.26 | 0 | 3 | 4 | 4 |
| P29692 | EEF1D | 0.57 | 0.26 | 0 | 2 | 0 | 0 |
| P26373 | RPL13 | 0.6 | 0.26 | 0 | 0 | 7 | 0 |
| P55795 | HNRNPH2 | 0.41 | 0.26 | 0 | 0 | 1 | 0 |
| Q07955 | SRSF1 | 0.56 | 0.25 | 0 | 2 | 1 | 0 |
| O75533 | SF3B1 | 0.58 | 0.25 | 0 | 9 | 5 | 0 |
| P09651 | HNRNPA1 | 0.4 | 0.24 | 0 | 4 | 7 | 0 |
| P23588 | EIF4B | 0.58 | 0.24 | 0 | 7 | 15 | 0 |
| P24534 | EEF1B2 | 0.5 | 0.24 | 0 | 1 | 0 | 0 |
| D6RBZ0 | HNRNPAB | 0.37 | 0.24 | 0 | 0 | 1 | 0 |
| P50990 | CCT8 | 0.59 | 0.24 | 0 | 2 | 1 | 0 |
| Q12905 | ILF2 | 0.61 | 0.24 | 0 | 2 | 14 | 0 |
| P22626 | HNRNPA2B1 | 0.39 | 0.23 | 0 | 14 | 13 | 2 |
| Q08170 | SRSF4 | 0.52 | 0.23 | 0 | 1 | 2 | 0 |

| | | | | | | | |
|---|---|---|---|---|---|---|---|
| Q58FG0 | HSP90AA5P | 0.57 | 0.23 | 0 | 1 | 1 | 4 |
| Q13200 | PSMD2 | 0.52 | 0.22 | 0 | 1 | 0 | 0 |
| P27635 | RPL10 | 0.41 | 0.22 | 0 | 1 | 0 | 0 |
| Q8NHW5 | RPLP0P6 | 0.56 | 0.22 | 0 | 0 | 9 | 0 |
| P67809 | YBX1 | 0.54 | 0.21 | 0 | 1 | 4 | 0 |
| P55072 | VCP | 0.53 | 0.21 | 0 | 0 | 1 | 0 |
| P12236 | SLC25A6 | 0.46 | 0.21 | 0 | 1 | 1 | 0 |
| P42166 | TMPO | 0.4 | 0.21 | 0 | 1 | 0 | 0 |
| P0DMV9 | HSPA1B | 0.33 | 0.2 | 0 | 12 | 14 | 2 |
| P0DMV8 | HSPA1A | 0.32 | 0.2 | 0 | 13 | 12 | 3 |
| Q13885 | TUBB2A | 0.35 | 0.2 | 0 | 13 | 16 | 3 |
| Q71U36 | TUBA1A | 0.35 | 0.2 | 0 | 12 | 15 | 2 |
| P10599 | TXN | 0.42 | 0.2 | 0 | 1 | 2 | 0 |
| P62807 | HIST1H2BC;HIST1H2BE; | 0.45 | 0.19 | 0 | 0 | 6 | 0 |
| P47756 | CAPZB | 0.46 | 0.19 | 0 | 0 | 2 | 0 |
| P16403 | HIST1H1C | 0.39 | 0.18 | 0 | 6 | 9 | 0 |
| P50914 | RPL14 | 0.4 | 0.18 | 0 | 0 | 3 | 0 |
| P35749 | MYH11 | 0.4 | 0.17 | 0 | 0 | 1 | 0 |
| P18621 | RPL17 | 0.33 | 0.17 | 0 | 0 | 1 | 0 |
| P62318 | SNRPD3 | 0.32 | 0.16 | 0 | 1 | 0 | 4 |
| P05386 | RPLP1 | 0.3 | 0.15 | 0 | 0 | 2 | 0 |
| Q13263 | TRIM28 | 0.26 | 0.15 | 0 | 0 | 1 | 0 |
| O60256 | PRPSAP2 | 0.39 | 0.15 | 0 | 1 | 3 | 0 |
| P61978 | HNRNPK | 0.3 | 0.14 | 0 | 4 | 3 | 0 |
| P98175 | RBM10 | 0.29 | 0.14 | 0 | 5 | 2 | 0 |
| P05388 | RPLP0 | 0.23 | 0.12 | 0 | 1 | 1 | 0 |
| P62987 | UBA52 | 0.36 | 0.12 | 0 | 2 | 5 | 14 |
| P62906 | RPL10A | 0.24 | 0.12 | 0 | 2 | 0 | 0 |
| P11182 | DBT | 0.28 | 0.11 | 0 | 1 | 0 | 0 |
| P35579 | MYH9 | 0.33 | 0.1 | 0 | 13 | 4 | 0 |
| Q13509 | TUBB3 | 0.2 | 0.1 | 0 | 3 | 4 | 0 |
| Q15393 | SF3B3 | 0.18 | 0.1 | 0 | 0 | 1 | 0 |
| P62736 | ACTA2 | 0.21 | 0.08 | 0 | 12 | 16 | 1 |
| Q9Y2W1 | THRAP3 | 0.2 | 0.08 | 0 | 1 | 0 | 0 |
| Q9BUF5 | TUBB6 | 0.14 | 0.07 | 0 | 0 | 1 | 0 |
| P07437 | TUBB | 0.14 | 0.07 | 0 | 8 | 4 | 0 |
| P60891 | PRPS1 | 0.19 | 0.07 | 0 | 2 | 0 | 0 |
| P10412 | HIST1H1E | 0.14 | 0.07 | 0 | 1 | 1 | 3 |
| P04350 | TUBB4A | 0.13 | 0.07 | 0 | 1 | 3 | 0 |
| Q9BQA1 | WDR77 | 0.14 | 0.06 | 0 | 0 | 3 | 0 |
| P09211 | GSTP1 | 0.12 | 0.05 | 0 | 3 | 2 | 0 |
| P21333 | FLNA | 0.11 | 0.05 | 0 | 0 | 7 | 0 |
| P68363 | TUBA1B | 0.07 | 0.04 | 0 | 0 | 1 | 0 |
| P60709 | ACTB | 0.09 | 0.04 | 0 | 5 | 17 | 0 |
| P68371 | TUBB4B | 0.06 | 0.04 | 0 | 0 | 2 | 0 |
| P16152 | CBR1 | 0.1 | 0.04 | 0 | 1 | 1 | 0 |
| O14744 | PRMT5 | 0.04 | 0.02 | 0 | 2 | 3 | 0 |

**Extended Data Table 2 | Demographic data of Crohn's Disease Patients**

| Genotype | Sex | Age of Diagnosis | Age of Collection |
|---|---|---|---|
| CD # 1 -/- SP140 SNP | M | 39 | 66 |
| CD # 2 -/- SP140 SNP | F | 35 | 54 |
| CD # 3 -/- SP140 SNP | M | 21 | 49 |
| CD # 1 +/+ SP140 SNP | M | 12 | 31 |
| CD # 2 +/+ SP140 SNP | M | 44 | 52 |
